# Aged white adipose tissue is rejuvenated in a young host through thermogenic remodeling of mature adipocytes

**DOI:** 10.64898/2025.12.28.696721

**Authors:** Alibek Moldakozhayev, Alexander Tyshkovskiy, Pasquale Nigro, Zehao Zhang, Albina Tskhay, Cecília G. de Magalhães, Shweta Yadav, Kejun Ying, Alec W. Eames, Jesse R. Poganik, Shakchhi Joshi, Ji Seul Han, Daria Kholdina, Wayne Mitchell, Joao A. Paulo, Josue A. Lopez, Kenta Ninomiya, Bohan Zhang, Erik Jacques, Sharif Iqbal, Praju Vikas Anekal, Stanislav Tikhonov, Andrei E. Tarkhov, Nicholas P. Carbone, Alexey V. Terskikh, Steven P. Gygi, Marcia Haigis, Michael F. Hirshman, Andrey A. Parkhitko, Wei Zhou, Junyue Cao, Laurie J. Goodyear, Jeremy M. Van Raamsdonk, Vadim N. Gladyshev

**Affiliations:** Division of Genetics, Department of Medicine, Brigham and Women’s Hospital, Harvard Medical School, Boston, MA, USA; Broad Institute of MIT and Harvard, Cambridge, MA, USA; Department of Neurology and Neurosurgery, McGill University, Montreal, QC, Canada; Metabolic Disorders and Complications, and Brain Repair and Integrative Neuroscience Programs, Research Institute of the McGill University Health Centre, Montreal, QC, Canada; Section on Integrative Physiology and Metabolism, Joslin Diabetes Center, Boston, MA, USA; Laboratory of Single-Cell Genomics and Population Dynamics, the Rockefeller University, New York, NY, USA; The David Rockefeller Graduate Program in Bioscience, the Rockefeller University, New York, NY, USA; Department of Health Sciences, Sargent College of Health and Rehabilitation Sciences, Boston University, Boston, MA, USA; Aging Institute of UPMC, University of Pittsburgh, Pittsburgh, PA, USA; Department of Neurology and Neurological Sciences, Stanford University School of Medicine, Stanford, CA, USA; Knight Initiative for Brain Resilience, Wu Tsai Neurosciences Institute, Stanford University, Stanford, CA, USA; Department of Cell Biology, Blavatnik Institute, Harvard Medical School, Boston, MA, USA; Department of Cancer Biology, Dana-Farber Cancer Institute, Boston, MA, USA; Department of Cell Biology, Harvard Medical School, Boston, MA, USA; Scintillon Institute, San Diego, CA, USA; Harry Perkins Institute of Medical Research, QEII Medical Centre and Centre for Medical Research, The University of Western Australia, Crawley, WA, Australia; MicRoN Core, Harvard Medical School, Boston, MA, USA; Independent researcher; Division of Endocrinology and Metabolism, Department of Medicine, University of Pittsburgh, Pittsburgh, PA, USA; Harvard Medical School, Boston, MA, USA; Division of Experimental Medicine, Department of Medicine, McGill University, Montreal, QC, Canada

**Keywords:** Aging, Rejuvenation, Longevity, Adipose tissue, Transplantation, Aging clocks, Biological age, Multi-omics, Single-nucleus RNA sequencing, Thermogenesis, Lipid droplet, Futile creatine cycle, Futile calcium cycle, Atp2a1, Lrmda, SERCA, *Drosophila*

## Abstract

Existing studies focus on endpoints, leaving unambiguous rejuvenation largely undemonstrated. To address this gap, we transplanted white adipose tissue (WAT) between young and old mice and longitudinally quantified biological age using molecular and histological approaches. Transplantation accelerated graft aging, and young WAT engrafted into old hosts acquired an aged state without rejuvenating the recipient. Conversely, old WAT engrafted into young hosts rejuvenated. Single-nucleus profiling localized rejuvenation to mature adipocytes and, within them, to chromatin, splicing, translation, and energy metabolism pathways. Rejuvenation was characterized by activation of thermogenesis and structural remodeling, and suppression of differentiation and lipid accumulation. Proteomic and metabolomic analyses supported these findings and implicated DNA repair. Histologically, rejuvenated WAT had smaller, more uniform lipid droplets, resembling young WAT. Top rejuvenation-associated genes were context-dependent modulators of lifespan in *Drosophila*. WAT’s biological age is therefore partially reversible, and the rejuvenation program components regulate longevity.

**Highlights:**

- Aged mouse adipose tissue is longitudinally rejuvenated in a young host
- Single-nucleus profiling localizes age reversal to mature adipocytes
- Rejuvenation activates thermogenesis and restores healthy lipid droplet morphology
- Rejuvenation-associated genes modulate lifespan in *Drosophila*

## Introduction

Aging is characterized by the progressive accumulation of deleterious molecular transformations that ultimately drive disease onset and organismal death ^1–4^. While slowing aging may delay the onset of age-related diseases, rejuvenation offers the potential for more radical anti-aging approaches by not only treating existing conditions but also preventing their future emergence. Heterochronic parabiosis (HPB), in which the circulatory systems of a young and an old animal are surgically joined, has long been studied as a powerful experimental model of rejuvenation. Studies have demonstrated that exposure of an old animal to a youthful systemic environment can improve molecular and functional features of aging in multiple tissues, such as muscle, liver, heart, brain, and bone, highlighting the potential of a young system to ameliorate age-associated decline ^5–11^. The most recent research employing epigenetic and transcriptomic profiling to assess biological age has shown that HPB can restore youthful gene expression profiles, reduce epigenetic and transcriptomic ages, enhance physiological functions, and ultimately extend the lifespan of older mice, even after the animals were separated ^12,13^.

Despite evidence from HPB that systemic factors can partially reverse organismal biological age, it remains unclear whether all tissues share comparable rejuvenation potential. Most studies have focused on whole-organism phenotypes or "canonical" tissues such as brain and liver, leaving tissue-specific rejuvenation largely unexplored. A direct way to address this gap is to transplant individual old organs or tissues into young hosts and assess the extent to which their biological age can be reset. Importantly, rejuvenation is often reported but rarely demonstrated. To show that a tissue has become younger, one must know how old it was to begin with, but almost no study makes measurements of a baseline state. Comparison of treated animals against untreated ones at a single time point cannot distinguish rejuvenation from slower aging. Longitudinal studies in humans treat unhealthy individuals and simply return them to a healthy state. Direct evidence that rejuvenation is possible therefore remains scarce.

To address these gaps, we focused on subcutaneous white adipose tissue (WAT) - a non-vital, easily accessible tissue that can be sampled prior to transplantation without substantial harm. This feature makes WAT uniquely suited for longitudinal assessment of biological age and provides a tractable model to study the intrinsic rejuvenation capacity of aged tissues *in vivo*. WAT is a highly dynamic endocrine organ essential for energy homeostasis and systemic metabolic regulation ^14,15^. In addition to serving as a primary site of lipid storage, WAT secretes a broad array of adipokines that influence organismal metabolism and coordinate the function of other tissues. With age, WAT undergoes marked dysfunction characterized by impaired progenitor activity, increased infiltration of senescent and pro-inflammatory cells, redistribution of fat from subcutaneous to visceral depots, and thermogenic capacity decline, driven by the gradual loss of beige and brown adipocytes – the cell populations associated with better metabolic health ^14^. This results in insulin resistance, chronic low-grade inflammation, and ectopic lipid accumulation that may result in obesity.

We established a longitudinal murine adipose tissue transplantation model in which old WAT was engrafted into young hosts, and young WAT into old hosts, to directly test the impact of systemic environment on tissue and cellular aging trajectories. We quantified biological age using epigenetic and transcriptomic clocks, resolved cell-type-specific changes in biological age at single-nucleus resolution, and performed molecular characterization of transcriptomic, proteomic, and metabolomic profiles. Histological assessment revealed structural correlates of the observed effects at tissue and organelle resolution. Top candidate genes emerging from this framework were tested in *D. melanogaster* to establish their relevance to longevity. Finally, we examined the capacity of heterochronic transplantation to reshape the systemic aging landscape of the host - asking not only whether old tissue can be rejuvenated, but whether transplanted young tissue can itself confer rejuvenating effects or *vice versa* for the transplanted old tissue. Our multimodal framework allowed rigorous evaluation of whether a young systemic environment can rejuvenate aged WAT, dissected the molecular pathways and structural outcomes underlying this process, and established functional links between adipose rejuvenation programs and lifespan regulation, extending the paradigm of young-blood–mediated rejuvenation to a central metabolic and endocrine organ.

## Results

### Experimental design

To explore the potential for adipose tissue rejuvenation, we transplanted subcutaneous inguinal WAT into the subcutaneous compartment in the interscapular region of recipient mice. We collected WAT samples from (1) donor animals prior to transplantation (WAT Before), (2) the same WAT retrieved from the graft site 10 weeks post-transplantation (WAT After), and (3) the untouched inguinal WAT of recipients (WAT Native), enabling within-animal longitudinal comparisons of aging dynamics. To assess whether adipose tissue extraction or its transplantation influenced systemic aging of the host, we additionally collected endpoint livers and blood from donor and recipient animals. Tissues were subjected to comprehensive multi-modal profiling: WAT Before and After were characterized using Mammalian DNA Methylation Array ^16^; ImAge, a single-nucleus chromatin imaging-based measure of epigenetic age ^17^; bulk and single-nucleus RNA sequencing; TMT proteomics; LC-MS metabolomics of polar and non-polar fractions; and histology, comprising hematoxylin and eosin (H&E), Masson’s trichrome, and mitochondrial immunofluorescence staining (**Figure 1A**). Native WAT and livers were profiled by the Mammalian DNA Methylation Array and bulk RNA sequencing. Finally, blood was profiled by the Mammalian DNA Methylation Array, and complete blood counts (CBC) were acquired using a Hemavet 950FS.

**Figure 1.**
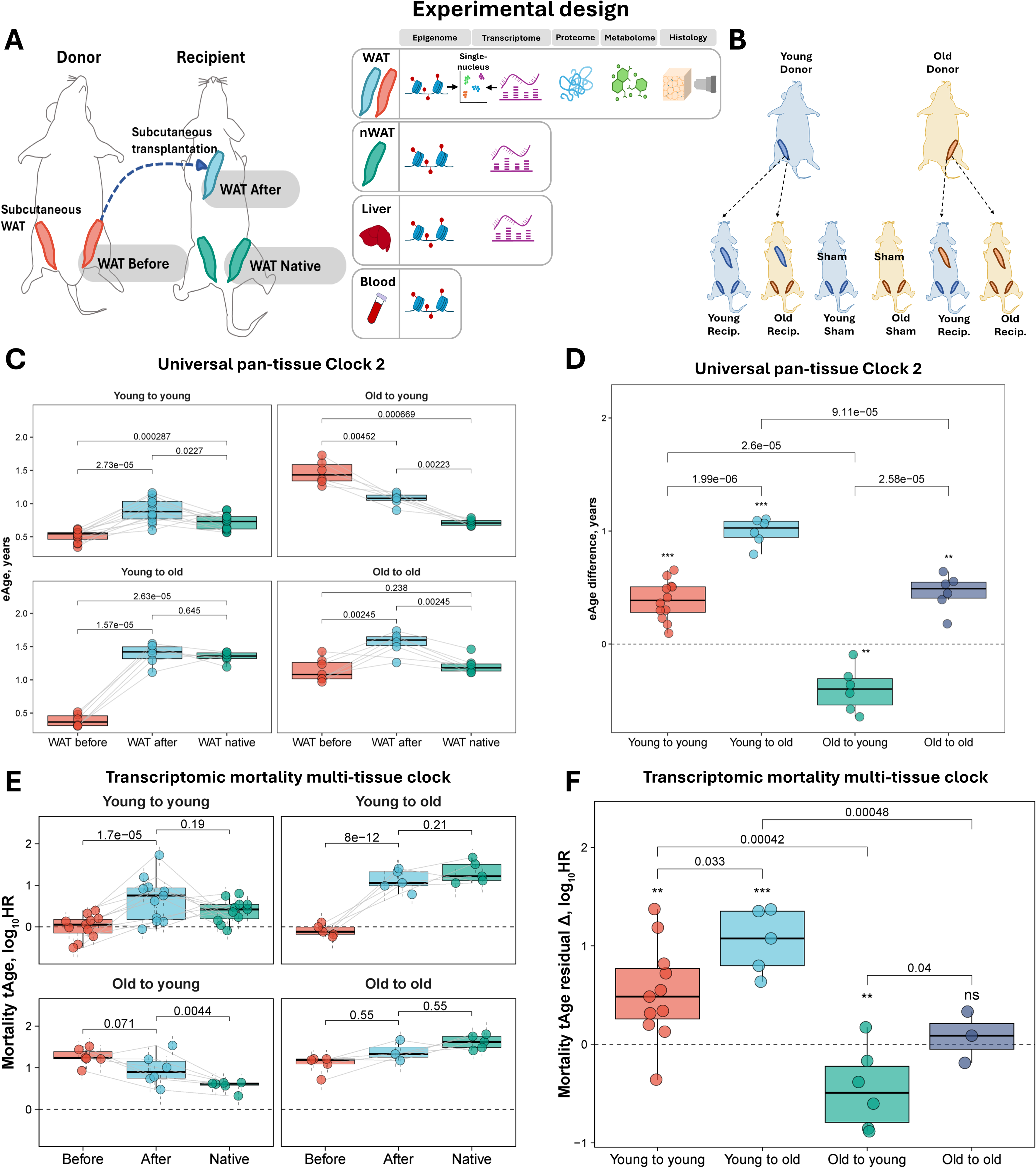
Epigenetic and transcriptomic aging clocks reveal rejuvenation of aged WAT in young hosts. **(A)** Experimental overview. Subcutaneous inguinal white adipose tissue (WAT) was surgically excised from donor mice and transplanted subcutaneously into recipient animals. To enable a longitudinal, within-animal assessment of biological aging, donor WAT was sampled immediately prior to transplantation (WAT Before) and recovered from the graft site 10 weeks post-transplantation (WAT After); both were subjected to comprehensive multi-omic profiling, comprising epigenetic, transcriptomic, proteomic, and metabolomic analyses, as well as histological characterization. Epigenetic and transcriptomic analyses were conducted at both bulk-tissue and single-nucleus levels. Endogenous, untouched inguinal WAT from recipient animals (WAT Native) was collected at the end of the study to assess the systemic influence of the transplantation on host tissue aging using epigenetic and transcriptomic clocks. Liver and blood were additionally collected from donor and recipient animals and subjected to epigenetic clock analyses to complement the evaluation of host-wide aging trajectories following WAT removal or implantation. **(B)** Schematic of the experimental setup. Heterochronic (old-to-young, young-to-old) and isochronic (young-to-young, old-to-old) transplantations were performed, with tissues collected before and after grafting. Blue color represents young, and yellow color represents old animals/tissues. **(C)** Epigenetic age (eAge) of WAT samples before and after transplantation across all experimental groups according to the Universal Clock 2 predictions. Mouse ID was included in the model as a covariate to ensure paired comparison of eAge dynamics. Each dot represents a sample collected from individual mice, n = 6-12 per group. **(D)** Comparison of epigenetic age (eAge) changes of WAT samples before and after transplantation across all experimental groups according to Universal Clock 2. Asterisks reflect statistical significance (BH-adjusted p-values) of eAge changes during transplantation within individual groups, whereas the significance of pairwise comparisons in eAge dynamics between groups is denoted in text. **(E)** Expected mortality (in log scale) of WAT samples before and after transplantation across all experimental groups according to the Bayesian Ridge (BR) rodent multi-tissue transcriptomic mortality clock. Statistical significance was assessed with mixed-effects models. Mouse ID was included in the model as a covariate to ensure paired comparison of tAge dynamics. Data are tAges ± s.d. Each dot represents a sample collected from individual mice, n = 3-11 per group. **(F)** Comparison of age-adjusted changes in expected mortality (in log scale) of WAT samples before and after transplantation across all experimental groups, calculated by the rodent multi-tissue transcriptomic mortality clock. Statistical significance was assessed with mixed-effects models. Asterisks reflect statistical significance (BH-adjusted p-values) of expected mortality changes during transplantation within individual groups, whereas the significance of pairwise comparisons in expected mortality dynamics between groups is denoted in text. Boxplots: center line indicates median; box limits, interquartile range; whiskers, ±1.5× IQR. Unless specified otherwise, statistical differences between groups are assessed with ANOVA and adjusted for multiple comparisons with the Benjamini-Hochberg approach. *** p.adjusted< 0.001, ** p.adjusted < 0.01, * p.adjusted < 0.05, ^ p.adjusted < 0.1

To rigorously evaluate the rejuvenation potential of WAT and isolate the reciprocal contributions of donor and host age, we established four experimental groups: the primary heterochronic conditions - old WAT transplanted into young recipients and young WAT transplanted into old recipients alongside isochronic controls of young-to-young and old-to-old transplantations. We also performed endpoint tissue collection from age-matched sham-surgery controls and adipose tissue donors in both young and old animals to delineate the effects of adipose tissue removal or implantation on the systemic aging of the host (**Figure 1B**).

### Epigenetic and transcriptomic aging clocks reveal rejuvenation of aged WAT in young hosts

To quantify biological age changes in transplanted adipose tissue, we applied two advanced pan-tissue mammalian epigenetic age estimators, Universal Clocks 2 and 3 ^18^, to collected WAT samples (**Figure 1C, Supplementary Figure 1A;** sample n = 6-12 per group). In isochronic transplantations (young-to-young and old-to-old), we observed a significantly increased predicted age in post-transplant grafts (WAT After) relative to both their baseline states (WAT Before) and the untouched adipose tissue from recipient animals (WAT Native), across both clocks. In heterochronic young-to-old transplants, young adipose tissue placed into old hosts acquired a biological age indistinguishable from that of the host’s own WAT Native, indicating that the aged systemic environment rapidly imposes an old epigenetic state on young grafts. In contrast, old adipose tissue implanted into young recipients underwent substantial rejuvenation, reflected by a marked and significant reduction in predicted age.

The longitudinal design of our study, which included pre-transplant clock measurements, allowed us to assess the magnitude and direction of biological age change for individual grafts using Universal Clocks 2 (**Figure 1D**) and 3 (**Supplementary Figure 1B**). Subtracting predicted values of WAT Before from those of WAT After showed that isochronic grafts aged significantly and to a similar extent, regardless of donor and host age. Young tissue implanted into old hosts underwent greater age acceleration than either isochronic group, underscoring the stronger influence of the old systemic environment on young tissue. In contrast, old adipose tissue transplanted into young hosts was the only group with a significant reversal of the aging trajectory, and these changes were significantly different from those of the old-to-old and young-to-young groups, indicating the capacity of a young environment to rejuvenate old tissue. A similar pattern emerged with the SVM Universal Mortality pan-tissue Clock ^19^, a pan-mammalian epigenetic predictor of expected mortality: expected hazard rose in isochronic grafts, although not significantly above native recipient WAT; young grafts in old hosts remained indistinguishable from recipient native tissue; and old grafts in young hosts showed a marked reduction, with within-sample observations mirroring those of the Universal Clocks (**Supplementary Figure 1C, 1D**). These results indicate that a young systemic environment can reverse both chronological- and mortality-linked epigenetic features of aged adipose tissue.

Because epigenetic age predictions may be influenced by global shifts in methylation, whether due to biological factors or technical variability such as coverage of probe hybridization, we compared average methylation levels across all groups. No significant differences were observed between any comparisons (**Supplementary Figure 1E**). We also evaluated whether transplantation affected epigenetic noise, as measured by Shannon entropy, which has been associated with aging and molecular disorder ^20^. We observed a general non-significant trend toward increased entropy in all post-transplant adipose tissue samples, with old adipose tissue transplanted into old recipients reaching a marginal significance (**Supplementary Figure 1F**). Notably, baseline Shannon entropy values were similar between young and old samples.

To assess whether the rejuvenation observed at the epigenetic level extended to gene expression, we applied rodent multi-tissue transcriptomic clocks of chronological age and expected mortality (sample n = 6-11 per group) ^13^. With the chronological clock (**Supplementary Figure 1G**), young-to-young grafts showed a significant increase in predicted transcriptomic age (tAge) over their pre-transplant baseline and a modest, non-significant elevation over recipient WAT Native; the corresponding increases in old-to-old grafts did not reach significance. Young grafts in old hosts converged on host tAge, mirroring the epigenetic clocks, whereas old grafts in young hosts exhibited a marginally significant decrease in tAge compared to baseline. Subtracting WAT Before from WAT After values and adjusting for chronological age (i.e., calculating age deviations) (**Supplementary Figure 1H**) showed that young tissue aged to the same extent in young and old hosts, although old hosts accelerated aging of young grafts more than that of old grafts. Old tissue in young hosts was again the only group to decelerate significantly, with changes significantly different from both isochronic controls.

The mortality clock produced nearly identical results across WAT Before, WAT After, and WAT Native (**Figure 1E**), suggesting that these expression changes also track features associated with mortality risk. In the age-adjusted within-graft analysis (**Figure 1F**), young tissue in old hosts aged significantly more than either isochronic group, whereas old tissue in young hosts again decelerated significantly, with changes significantly different from both isochronic controls.

To delineate shared and distinct biological pathways underlying the transcriptomic responses to transplantation, we applied module-specific rodent multi-tissue transcriptomic clocks of chronological age and mortality, which were designed to predict tAge at the level of individual cellular subsystems (modules) ^13^. We did not detect any modules that showed significant rejuvenation or aging in transplantation scenarios involving old graft tissue (**Supplementary Figure 1I**). In contrast, transplantation involving young grafts, regardless of recipient age, was associated with a strong pro-aging signal according to multiple functional modules. Modules that exhibited the strongest tAge acceleration for both young-to-young and young-to-old transplantations, supported by both chronological and mortality clocks, included adaptive immunity/T cell signaling, innate immunity/inflammation, cholesterol metabolism/platelet degranulation, and respiration/mitochondrial translation. These results suggest that these metabolic and immune domains are particularly sensitive to stress following the transplantation of young adipose tissue.

Overall, our results show that the biological age of a tissue is strongly influenced by both the transplantation procedure and the age of the host.

### Single-nucleus profiling and module clocks localize and characterize rejuvenation in adipocytes

Most aging clocks trained on bulk tissue data cannot distinguish whether a molecular age signal reflects cell-intrinsic states or shifts in cell composition ^21,22^. This limitation is problematic for rejuvenation studies based on contact with a young systemic environment, where compositional change may mean infiltration by young host immune cells - a process that could mimic rejuvenation and confound the interpretation. To determine whether the rejuvenation of old WAT transplanted into young hosts arose within post-mitotic mature donor adipocytes themselves, we performed single-nucleus RNA sequencing (snRNA-seq) of WAT Before and WAT After across all four transplantation groups (sample n = 1-3 per group). We recovered 165,013 nuclei that resolved into fourteen cell populations: fat cells (mature adipocytes), macrophages, preadipocytes, dendritic cells, B cells, T cells, mesothelial cells, monocytes, endothelial cells, neutrophils, pericyte cells, mast cells, smooth muscle cells, and epithelial cells (**Figure 2A**). As suspected, WAT After tissues showed enhanced immune cell infiltration, particularly by macrophages (**Supplementary Figure 2A, 2B**). Principal component analysis of pseudo-bulk meta-cell profiles grouped major sequenced populations into three superclusters — adipocytes with preadipocytes and endothelial cells, macrophages with monocytes, and T cells with B cells (**Figure 2B**).

**Figure 2.**
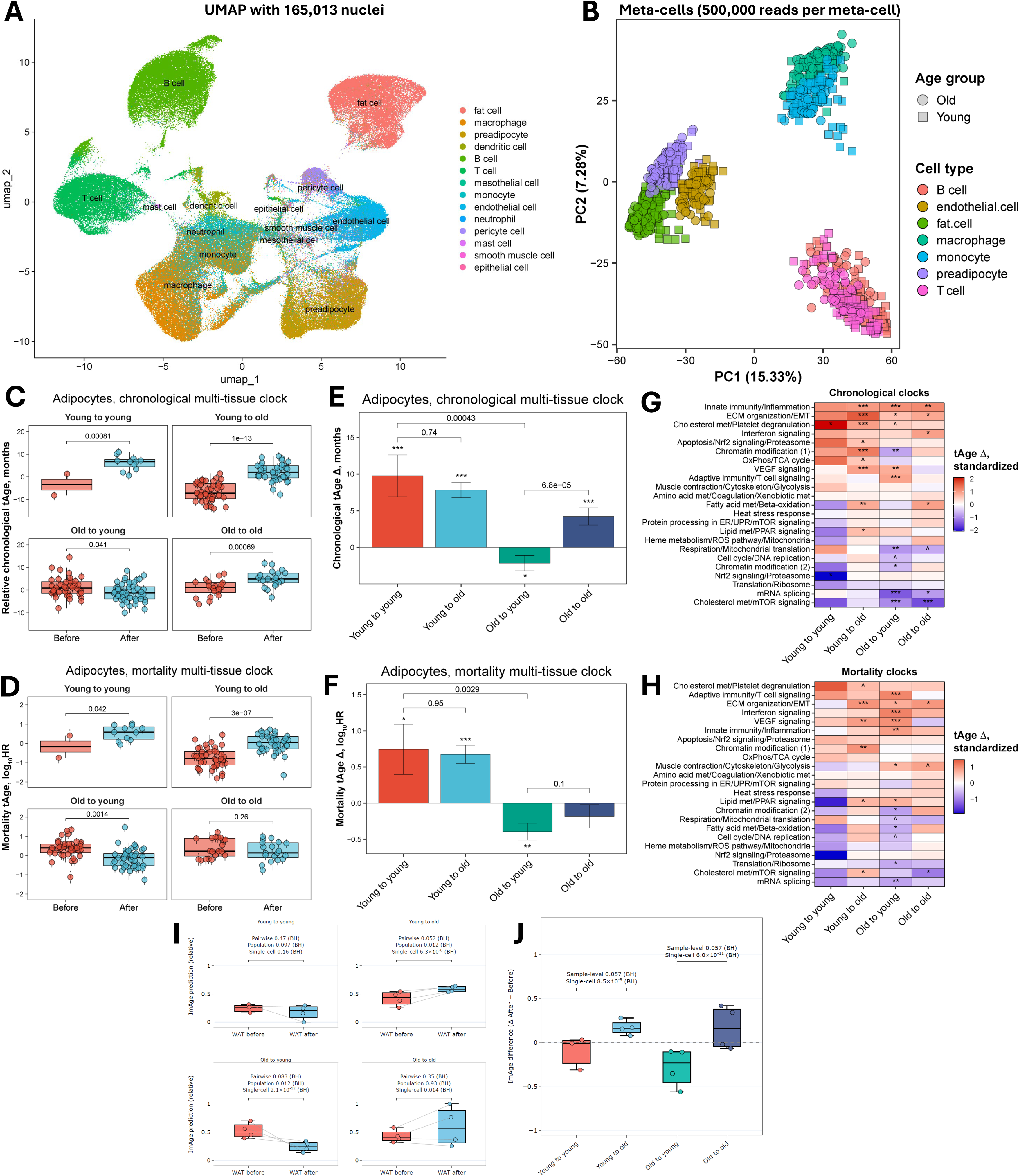
Single-nucleus profiling and module clocks localize and characterize rejuvenation in adipocytes. **(A)** UMAP visualization of single-nucleus RNA sequencing (snRNA-seq) data from subcutaneous white adipose tissue of young and old donor mice before and after transplantation (165,013 nuclei pooled across all four experimental groups). Each dot represents a single nucleus, colored by cell type identity. Fourteen major cell populations were identified, including fat cells (mature adipocytes), macrophages, preadipocytes, dendritic cells, B cells, T cells, mesothelial cells, monocytes, endothelial cells, neutrophils, pericyte cells, mast cells, smooth muscle cells, and epithelial cells. **(B)** Principal component analysis (PCA) of meta-cell transcriptomic profiles derived from snRNA-seq data, computed for seven major cell types. Each point represents a meta-cell (500,000 reads aggregated), colored by cell type and shaped by age group (circle = old, square = young). PC1 (15.33%) and PC2 (7.28%) together capture the dominant sources of transcriptomic variance across cell types. Three broad super-clusters are apparent: macrophages and monocytes cluster together in the upper right, fat cells, preadipocytes, and endothelial cells co-cluster on the left, and T cells and B cells form a distinct cluster in the lower right. **(C)** Transcriptomic age (tAge) of adipocytes before and after transplantation across all experimental groups according to the Bayesian Ridge (BR) mouse multi-tissue chronological clock. Statistical significance was assessed with mixed-effects models. Mouse ID was included in the model as a covariate to ensure paired comparison of tAge dynamics. Data are tAges ± s.d. Each dot represents a meta-cell compiled from individual mouse samples, n = 1-3 per group. **(D)** Comparison of chronological tAge dynamics in adipocytes over 10 weeks after transplantation across groups. For each transplantation group, the slope was estimated using a mixed-effects model with donor fat identity included as a covariate; p-values testing whether the slope differs from 0 are indicated with asterisks. Pairwise comparisons of tAge changes between groups were performed using mixed-effects model, with the p-value for the interaction term shown in the plot. P-values were adjusted for multiple comparisons using the Benjamini-Hochberg approach. Data are slope estimate ± SE. **(E)** Standardized change of tAge in adipocytes following transplantation across all experimental conditions, according to module-specific mouse multi-tissue transcriptomic chronological clocks. Red color indicates increased tAge, while blue color indicates decreased tAge after transplantation relative to their respective pre-transplantation baselines. Asterisks reflect statistical significance (BH-adjusted p-values) of changes, assessed with ANOVA with donor fat identity included as a covariate. Individual mouse samples, n = 1-3 per group. **(F)** Transcriptomic age (tAge) of adipocytes before and after transplantation across all experimental groups according to the BR mouse multi-tissue mortality clock. Statistical significance was assessed with mixed-effects models. Mouse ID was included in the statistical model as a covariate to ensure paired comparison of tAge dynamics. Data are tAges ± s.d. Each dot represents a meta-cell compiled from individual mouse samples, n = 1-3 per group **(G)** Comparison of mortality tAge dynamics in adipocytes over 10 weeks after transplantation across groups. For each transplantation group, the slope was estimated using a mixed-effects model with donor fat identity included as a covariate; p-values testing whether the slope differs from 0 are indicated with asterisks. Pairwise comparisons of tAge changes between groups were performed using mixed-effects model, with the p-value for the interaction term shown in the plot. P-values were adjusted for multiple comparisons using the Benjamini-Hochberg approach. Data are slope estimate ± SE. **(H)** Standardized change of tAge of adipocytes following transplantation across all experimental conditions, according to module-specific mouse multi-tissue transcriptomic mortality clocks. Red color indicates increased tAge, while blue color indicates decreased tAge after transplantation relative to their respective pre-transplantation baselines. Asterisks reflect statistical significance (BH-adjusted p-values) of changes, assessed with ANOVA with donor fat identity included as a covariate. Individual mouse samples, n = 1-3 per group. **(I)** Within-condition before→after changes in adipocyte ImAge across transplant conditions. Three levels of comparison are shown: Pairwise (paired t-test across transplant units), Population (Mann–Whitney U, graft sample medians versus the donor pool of the corresponding age), and Single-cell (Mann–Whitney U on the individual nuclei). Shown are BH-adjusted values. Tests are one-sided for the conditions with a pre-specified direction (Old to young, rejuvenation; Young to old, aging) and two-sided for the environment-matched conditions (Old to old, Young to young). All values are leave-one-mouse-out cross-validated estimates. Each dot represents a sample collected from individual mice, n = 3-4 per group. **(J)** Comparison of changes in adipocyte ImAge before and after transplantation. Sample level: Mann–Whitney U on the per-transplant Δ. Single-cell level: Mann–Whitney U on the per-cell Δ, each graft nucleus expressed relative to the median of its own donor’s pre-transplant nuclei. Both tests are two-sided; BH-adjusted values are shown. Boxplots: center line indicates median; box limits, interquartile range; whiskers, ±1.5× IQR. Unless specified otherwise, statistical differences between groups are assessed with ANOVA and adjusted for multiple comparisons with the Benjamini-Hochberg approach. *** p.adjusted< 0.001, ** p.adjusted < 0.01, * p.adjusted < 0.05, ^ p.adjusted < 0.1

We first asked whether either immune supercluster could account for the rejuvenation detected in bulk data, assuming that immune cells in WAT Before were donor-derived, whereas most immune cells in WAT After were recipient-derived. Applying the rodent transcriptomic chronological and mortality multi-tissue clocks to pooled macrophages and monocytes, we found that these cell populations aged significantly only in young-to-old transplants (both clocks) and old-to-old transplants (chronological clock; **Supplementary Figure 2C, 2D**). No population rejuvenation was observed in old-to-young transplants. Comparison of the change magnitudes showed that macrophage and monocyte populations generally aged more in old than in young hosts, although this trend was largely non-significant and reached marginal significance only for the young-to-old versus young-to-young comparison (**Supplementary Figure 2E, 2F**). Therefore, macrophages and monocytes would, if anything, bias the bulk signal toward older values, excluding infiltration of these cells as a source of the rejuvenation signal observed in bulk-analyzed old-to-young grafts.

Pooled T and B cell populations aged significantly by both chronological and mortality clocks only in transplantations involving young grafts (**Supplementary Figure 2G, 2H**). No significant rejuvenation of T and B cell populations occurred in old-to-young transplants, although tAge trended below baseline. Comparison of the change magnitudes indicated that T and B cell populations generally aged more in old than in young hosts (**Supplementary Figure 2I, 2J**), consistent with recent observations ^23^. Among old grafts, the difference between old-to-young and old-to-old changes was significant by the chronological clock, whereas the young-to-old versus young-to-young comparison reached marginal significance by the mortality clock. The young-to-young versus old-to-young comparison was highly significant by both clocks, indicating that the direction of tAge change differs markedly between young and old grafts placed into young hosts. Unlike macrophages and monocytes, T and B cells may therefore contribute to the transcriptomic rejuvenation signal detected at the bulk level.

Most importantly, we applied the same clocks to mature adipocyte nuclei (**Figure 2C, 2D**). Adipocytes aged significantly by both the chronological and mortality clocks in transplants involving young grafts, and in old-to-old transplants by the chronological clock. In contrast, old adipocytes transplanted into young hosts were the only population to undergo a significant reduction in predicted tAge, according to both clocks. Comparison of the change magnitudes showed that the direction of tAge change in old-to-young transplants differed significantly from that in young-to-young transplants (both clocks) and old-to-old transplants (chronological clock; **Figure 2E, 2F**). Together, these results localize the rejuvenation detected at the bulk transcriptomic level to the donor’s mature adipocytes themselves.

Extensive differential expression accompanied observed tAge shifts, ranging from several hundred genes in young-to-young grafts to over a thousand in the other groups (**Supplementary Figure 2K**). To resolve gene regulatory programs underlying adipocyte aging and rejuvenation, we applied module-specific chronological and mortality clocks to adipocyte nuclei (**Figure 2G, 2H**). The modules that aged in old-to-young adipocytes by both clocks were immune, inflammatory, vascular, and ECM/EMT programs. ECM organization/EMT module also aged by both clocks in young-to-old and old-to-old transplants and trended in the pro-aging direction in young-to-young transplants by the chronological clock, suggesting that changes in this module reflect the detrimental nature of the transplantation procedure itself.

A distinct set of modules rejuvenated in old adipocytes transplanted into young hosts. Chromatin modification (2) and mRNA splicing modules rejuvenated most consistently, reaching significance by both clocks; respiration/mitochondrial translation rejuvenated significantly by the chronological and marginally by the mortality clock, and cell cycle/DNA replication rejuvenated marginally by both clocks. Chromatin modification (1) and cholesterol metabolism/mTOR signaling modules rejuvenated by the chronological clock alone, whereas fatty acid metabolism/beta-oxidation and translation/ribosome modules rejuvenated by the mortality clock alone. Two of these modules also significantly rejuvenated in old-to-old transplants: mRNA splicing by the chronological clock alone, and the cholesterol metabolism/mTOR signaling module by both clocks. The latter module rejuvenation therefore appears to be a general response of old tissue to transplantation rather than a feature of the young host environment.

To test whether the transcriptomic rejuvenation of chromatin-related modules in adipocytes extended to chromatin organization itself, we applied ImAge, a chromatin imaging-based measure of epigenetic age in single nuclei ^17^, to WAT Before and WAT After across all transplantation conditions (sample n = 3-4 per group). The analysis was restricted to mature adipocytes, gated as C/EBPα-positive and PU.1-negative.

Isochronic conditions showed no strong aging signal (**Figure 2I**). Young adipocytes in young recipients did not change significantly at any level of comparison, whereas old adipocytes in old recipients aged significantly only at the single-cell level. Young adipocytes transplanted into old recipients aged significantly at the population sample-level average and single-cell levels, with the pairwise sample-level average comparison reaching marginal significance after adjustment for multiple hypothesis testing. Old adipocytes implanted into young recipients underwent significant rejuvenation at the population sample-level average and single-cell levels, with the pairwise sample-level average comparison reaching marginal significance.

Next, we tested whether these changes in ImAge differed from those observed in the corresponding isochronic control conditions (**Figure 2J**). Young adipocytes transplanted into old recipients aged significantly more than young adipocytes transplanted into young recipients at the single-cell level, whereas the comparison of sample-level average dynamics reached marginal significance. Likewise, old adipocytes transplanted into young recipients changed in the direction opposite to that of old adipocytes transplanted into old recipients at the single-cell level, with the comparison of sample-level average dynamics reaching marginal significance.

Overall, our findings localize rejuvenation to the aged adipocytes themselves, showing that a young systemic environment is capable of intrinsically resetting the biological age of these post-mitotic cells.

### Transcriptomic signature of WAT rejuvenation

To gain functional insight into the molecular changes associated with aging and rejuvenation, we analyzed bulk transcriptional responses within grafts across transplantation conditions (**Figure 3A**; sample n = 3-11 per group). All transplantations, especially those involving either a young graft or a young recipient, elicited numerous gene expression changes in the grafted adipose tissue. The most extensive response was observed in old-to-young transplants, with over 2,000 genes (p.adjusted < 0.05) significantly up- or downregulated after transplantation. Changes observed in the old-to-young group were highly and significantly correlated (Pearson’s r > 0.77 for all transplantation groups) with those from all other transplant conditions (**Figure 3B**). The weakest correlation (Pearson’s r = 0.4) was observed between the isochronic groups (young-to-young vs. old-to-old).

**Figure 3.**
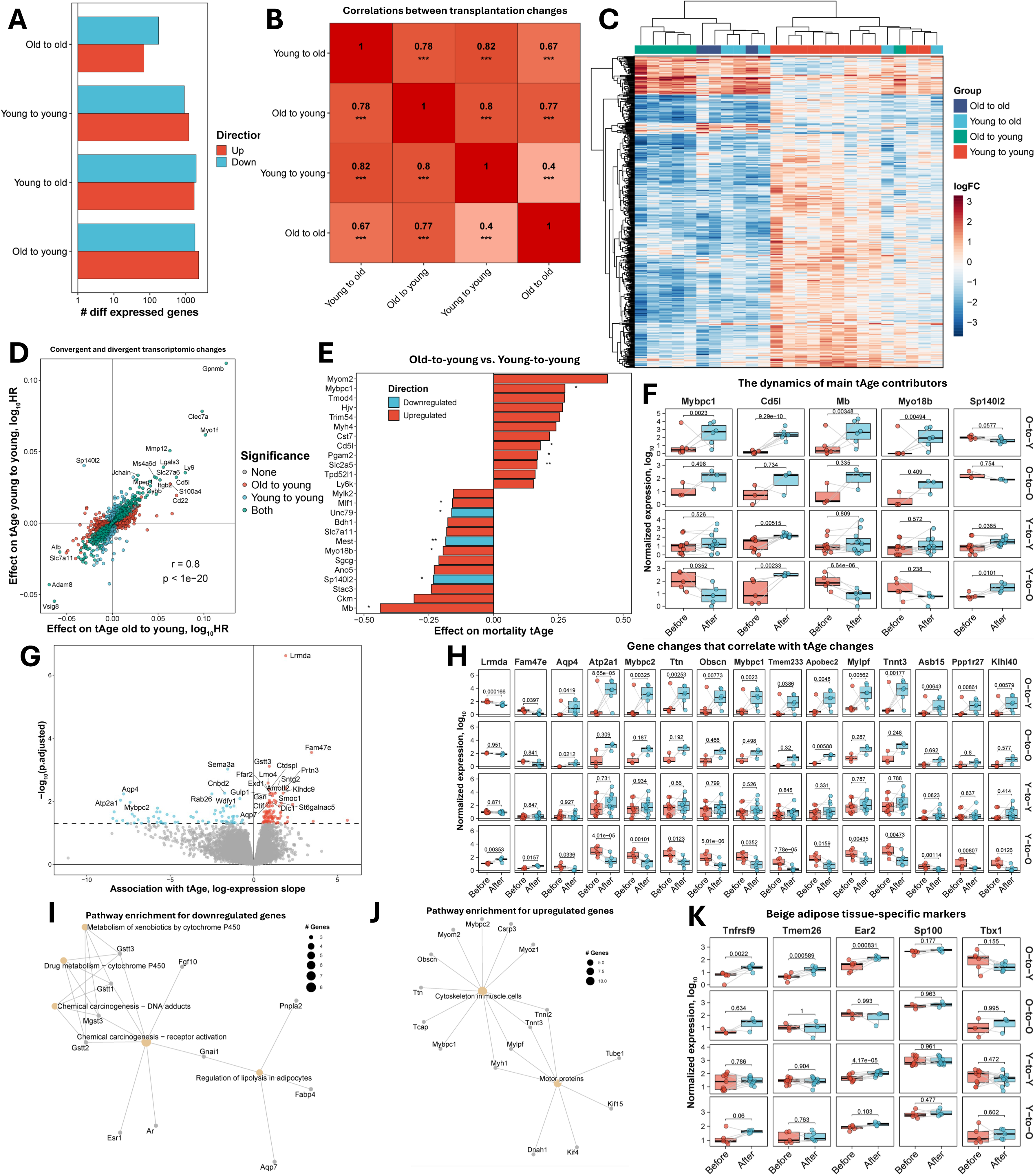
WAT rejuvenation engages thermogenic and muscle-like transcriptional programs. **(A)** Number of differentially expressed genes (p.adjusted < 0.05) in a donor’s WAT after transplantation relative to their respective pre-transplantation baselines for each experimental group. Barplots indicate the numbers of differentially expressed genes, and colors denote the direction of changes. **(B)** Pearson’s correlations of gene expression changes in WAT samples after transplantation across all experimental groups. Red indicates positive correlation, and the number indicates the correlation coefficient (Pearson’s r). Asterisks reflect statistical significance (BH-adjusted p-values). **(C)** Hierarchical clustering of transcriptional responses in WAT samples before and after transplantation across all experimental conditions. The direction of logFC is reflected with color. **(D)** Convergent and divergent gene expression changes induced by transplantation in a donor’s WAT that contribute to the change in tAge for young-to-young and old-to-young groups. Pearson’s correlation coefficient and its p-value are indicated in text. **(E)** Top 25 genes contributing to increased and decreased tAge following transplantation in old-to-young vs young-to-young group, according to the rodent multi-tissue transcriptomic mortality clock. Color and asterisks denote direction and statistical significance (BH-adjusted p-value) of difference in expression dynamics between groups, respectively. **(F)** Normalized log-expression of top genes, contributing to increased and decreased tAge of WAT samples following transplantation in old-to-young vs young-to-young group according to the rodent multi-tissue transcriptomic mortality clock. **(G)** Volcano plot displaying genes, whose upregulation during transplantation in a donor’s WAT is associated with higher (red) or lower (blue) tAge across all transplantation conditions according to the rodent multi-tissue transcriptomic mortality clock. **(H)** Top 15 genes whose normalized log-expression changes are most significantly associated with tAge changes following the transplantation across all experimental groups, according to the rodent multi-tissue transcriptomic mortality clock, ranked by statistical significance (BH-adjusted p-value). See Supplementary File 1 for the full gene list. **(I-J)** Pathway enrichment for genes negatively **(I)** and positively **(J)** associated with tAge dynamics during transplantation according to the rodent multi-tissue transcriptomic mortality clock. Grey nodes represent individual genes, and yellow nodes represent enriched pathways. Yellow node size reflects the number of significantly associated genes in each pathway, and edges indicate semantic similarity between terms. Only statistically significant terms (BH-adjusted p < 0.05, based on Fisher’s exact test) are shown. **(K)** Normalized log-expression dynamics of established beige adipose tissue-specific markers in WAT samples before and after transplantation across all experimental groups. Boxplots: center line indicates median; box limits, interquartile range; whiskers, ±1.5× IQR. Number of samples from individual mice = 3-11 per group. Unless specified otherwise, statistical differences between groups are assessed with ANOVA and adjusted for multiple comparisons with the Benjamini-Hochberg approach. *** p.adjusted< 0.001, ** p.adjusted < 0.01, * p.adjusted < 0.05, ^ p.adjusted < 0.1.

To identify biological pathways underlying these correlations, we performed a functional enrichment analysis of differentially expressed genes across transplant groups (**Supplementary Figure 3A, Supplementary File 1**). All groups displayed upregulation of inflammatory and immune pathways, such as interleukin signaling and general inflammatory response - established hallmarks of aging and mortality in mice ^24^. This shared response likely reflects immune infiltration triggered by transplantation of exogenous tissue. The young-to-young group also experienced a pronounced suppression of metabolic and energy pathways, such as fatty acid metabolism, mitochondrial translation, oxidative phosphorylation, and respiratory electron transport.

Next, we performed a clustering analysis of gene expression changes for the genes across all transplantation events. The strongest transcriptomic divergence was observed between old-to-young and young-to-young groups (**Figure 3C**), consistent with the largest number of differentially expressed genes identified between these two conditions (>2,000 genes with p.adjusted < 0.05; **Supplementary Figure 3B**). A closer examination of mortality-associated genes revealed an extensive overlap between young-to-young and old-to-young groups, with the same genes increasing and decreasing transcriptomic age in both groups, yielding a strong correlation in expression changes (Pearson’s r = 0.8; **Figure 3D**).

To understand why a young systemic environment rejuvenates aged adipose tissue but accelerates aging in young grafts, we compared young-to-young and old-to-young groups to identify the key differences in gene expression changes contributing to transcriptomic clock prediction differences across these groups. The most consistent top contributors to higher tAge predictions in the old-to-young group compared to young-to-young animals, according to both chronological (**Supplementary Figure 3C**) and mortality (**Figure 3E**) clocks, were upregulated *Mybpc1* and *Cd5l*, while the most consistent contributors to decreased tAge in the old-to-young group were upregulated *Mb* and *Myo18b*, and downregulated *Sp140l2*.

Next, we examined the expression dynamics of the top consistent contributing genes across all transplantation conditions (**Figure 3F**). In old adipose tissues implanted into young recipients, all top genes, except for *Sp140l2*, demonstrated a significant upregulation relative to the baseline levels. *Cd5l* was significantly upregulated in all groups except old-to-old, suggesting that strong macrophage recruitment ^25–27^ to the graft requires at least one young component, either the graft or the host environment. Interestingly, *Mybpc1*, *Mb*, and *Myo18b*, all muscle-related genes ^28^, exhibited coordinated regulation across samples, with expression changes occurring in the same direction and magnitude for all samples. Moreover, the directionality of changes for these genes was opposite between old-to-young and young-to-old conditions and absent in isochronic comparisons, suggesting that these genes may be specifically linked to rejuvenation or aging of adipose tissue rather than to general wound healing or graft stabilization.

### WAT rejuvenation engages thermogenic and muscle-like transcriptional programs

To identify genes associated with adipose tissue rejuvenation, we performed a transcriptomic screen by incorporating all experimental groups into a joint analysis. Specifically, we searched for genes whose expression changes correlated with changes in predicted expected hazard values derived from the transcriptomic mortality clock. In total, we identified 172 genes (p.adjusted < 0.05) whose expression changes significantly correlated with the changes in biological age (**Figure 3G**). The top 15 genes included *Lrmda*, *Fam47e*, *Aqp4*, *Atp2a1* (SERCA1), *Mybpc2*, *Ttn*, *Obscn*, *Mybpc1*, *Tmem233*, *Apobec2*, *Mylpf*, *Tnnt3*, *Asb15*, *Ppp1r27*, *Klhl40* (**Figure 3H**, full list available in **Supplementary File 1**).

Closer inspection of gene expression dynamics (**Figure 3H**) revealed coordinated changes among sets of genes, similar to those observed for *Mybpc1*, *Mb*, and *Myo18b* in previous analysis. These genes changed in opposite directions in old-to-young versus young-to-old transplantations, while showing minimal change in the isochronic transplantations (old-to-old and young-to-young).

To determine whether these rejuvenation-associated genes shared common biological functions, we performed a pathway enrichment analysis. Genes downregulated during rejuvenation were significantly enriched for pathways associated with chemical carcinogenesis, metabolism of xenobiotics by cytochrome P450, and regulation of lipolysis in adipocytes pathways (**Figure 3I**). Many of the genes related to cytochrome P450 and chemical carcinogenesis were glutathione transferases (*Gstt1/ Gstt2/Gstt3*, and *Mgst3*), involved in cellular detoxification and protection from oxidative damage, particularly lipid peroxidation in adipocytes ^29–31^. Among the lipolysis-related genes downregulated during rejuvenation were *Aqp7*, *Fabp4*, and *Pnpla2*. Finally, downregulated genes associated with rejuvenation included *Gnai1*, which connected chemical carcinogenesis and regulation of lipolysis pathways. In contrast, genes upregulated during rejuvenation were enriched for cytoskeletal and motor protein pathways commonly expressed in muscle cells (**Figure 3J**).

To investigate macrophage infiltration suggested by single-nuclei data, and elevation of *Cd5l* levels in all groups except the old-to-old scenario, we examined the expression dynamics of immune and adipocyte lineage markers. Specifically, we assessed *Cd68* ^32^, *Mrc1* ^33^, *Cd8a* ^34^, and the adipocyte marker *Pparg* ^35^ to infer shifts in cellular composition following transplantation (**Supplementary Figure 3D**). We hypothesized that increased *Cd68* and *Cd8a* expression, coupled with reduced *Pparg* levels, would indicate heightened immune cell infiltration and a relative decline in adipocyte-specific signals within bulk RNA-seq profiles. Consistent with the *Cd5l* findings, *Cd68* expression was significantly upregulated in all transplantation conditions except old-to-old, confirming robust macrophage infiltration when either the graft or recipient was young. *Mrc1* levels were selectively elevated in the old-to-young and young-to-young groups, reflecting enhanced recruitment of anti-inflammatory macrophages in young environments. *Cd8a* was significantly upregulated only in young grafts implanted into old recipients, indicating increased cytotoxic T-cell infiltration. Finally, *Pparg* expression was significantly reduced in both old-to-young and young-to-old conditions, suggesting the mitigation of the signal from adipocytes for these groups during bulk adipose tissue sequencing. Consequently, decreased or unchanged expression of adipocyte-specific genes and increased expression of immune-related genes in the bulk molecular data may partially reflect these compositional changes, masking true transcriptional alterations within cell types.

To determine whether established thermogenesis markers were altered in aged adipose tissue following transplantation into young recipients, we first examined expression changes of well-characterized brown and beige adipocyte thermogenic markers, including *Ebf2*, *Prdm16*, *Ppara*, *Cidea*, *Dio2*, *Cox8b*, *Cox7a1*, *Ucp1*, and *Atp2a2* ^36–40^ (**Supplementary Figure 3E**). *Ebf2* and *Prdm16* were suppressed in both old-to-young and young-to-old transplantations, while *Ppara* was significantly downregulated only in the old-to-young group. *Cidea* levels decreased in every group except the old-to-old and *Dio2* levels increased only in the young-to-old group. *Cox8b* significantly decreased in young grafts regardless of recipient ages, and *Cox7a1* remained unchanged across conditions. Finally, neither *Ucp1*, the most well-known marker of thermogenic activation, nor *Atp2a2*, encoding a calcium pump involved in UCP1-independent thermogenesis, were significantly altered in any transplantation scenario. Together, these findings indicate no activation of the canonical thermogenic program, mainly related to brown adipose tissue (BAT), during adipose tissue rejuvenation.

To further explore the possibility that rejuvenation engages thermogenic programs, we assessed the expression of established beige adipocyte-specific markers (**Figure 3K)**. *Tnfrsf9* (also frequently reported as CD137) and *Tmem26* - the most robust and validated markers of beige adipocytes ^37^ were markedly and exclusively upregulated in the old-to-young group, providing strong molecular evidence for beige phenotype induction upon rejuvenation. We further evaluated *Ear2*, *Sp100*, and *Tbx1*, a subset of beige-specific genes proposed to be activated in response to *Ebf2* downregulation ^39^. *Ear2* was significantly upregulated in both old-to-young and young-to-young groups, whereas *Sp100* and *Tbx1* showed no significant changes in any transplantation condition.

Collectively, our findings suggest that the transcriptional signature of WAT rejuvenation is characterized by partial engagement of known thermogenic programs and activation of a muscle-like program.

### Proteomic and metabolomic profiling corroborate and extend the transcriptomic signature of WAT rejuvenation

To corroborate the transcriptomic signatures at the protein level, we performed quantitative proteomic profiling of old adipose tissue transplanted into young (old-to-young) and old (old-to-old) recipients (sample n = 3 per group) using tandem mass tags ^41^. Similarly to transcriptomic, protein-level changes were strongly and positively correlated between the two contrasts (Pearson’s r = 0.769, **Figure 4A**), defining a convergent response shared by both. The concordantly upregulated proteins were dominated by immune system response markers, including MMP12, GPNMB, CTSS, CTSD, LGALS3, and LYZ2 ^42^, whereas the concordantly downregulated proteins were adipocyte and lipid-handling markers such as LEP, CIDEC, and MOGAT1 ^43–45^. This convergent signature supports transcriptomic findings, indicating that increased immune infiltration and consequent dilution of adipocyte signals accompany adipose tissue transplantation, regardless of host age.

**Figure 4.**
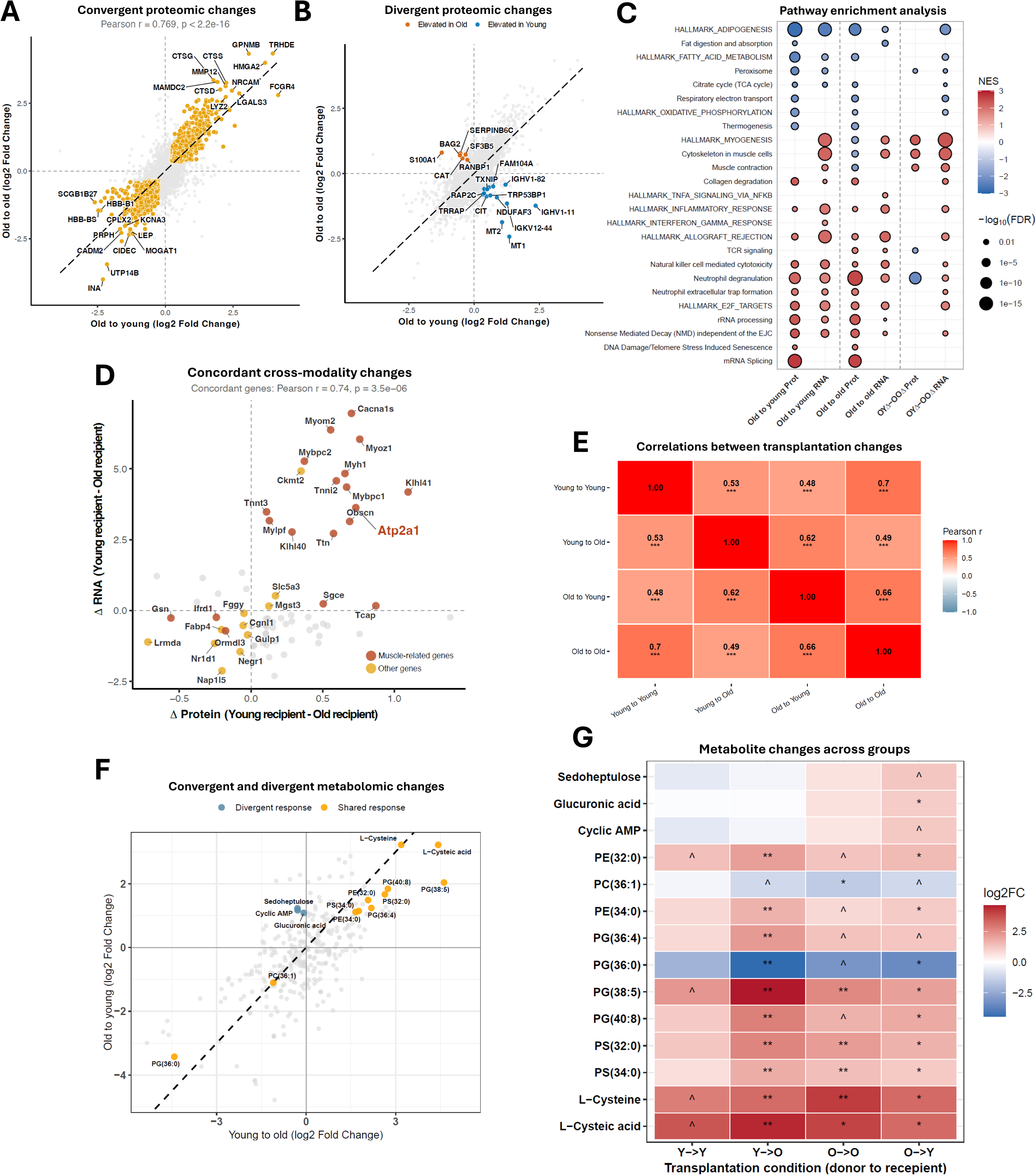
Proteomic and metabolomic profiling corroborate and extend the transcriptomic signature of WAT rejuvenation. **(A)** Convergent proteomic changes induced in old-to-old and old-to-young transplantation groups. Scatter plot of protein-level logFC for the rejuvenation contrast (Old to young; x-axis) versus the old host contrast (Old to old; y-axis). Orange points indicate proteins that significantly changed in both groups in the same direction after transplantation (concordant; p.adjusted < 0.05 in both). Grey points represent discordant or non-significantly changing proteins. A linear regression line is shown (dashed black). Top concordant proteins are labeled by gene symbol. Pearson’s r indicates the correlation coefficient, and p indicates the statistical significance (p-value) of the observed correlation. **(B)** Divergent proteomic changes induced in the old-to-young transplantation group compared to the old-to-old transplantation group. Scatter plot as in the shared response panel, highlighting proteins with significant changes in opposite directions between the old-to-young and old-to-old transplantations (p.adjusted < 0.1 in both). Blue indicates individual proteins that increase following old-to-young transplantation but decrease following old-to-old transplantation, while orange indicates proteins that decrease following old-to-young transplantation but increase following old-to-old transplantation. **(C)** Gene set enrichment analysis (GSEA) comparing proteomics and transcriptomics signatures across transplantation contrasts. Dot color indicates the normalized enrichment score (NES; red = positive enrichment, blue = negative enrichment), and dot size reflects statistical significance (−log10 p.adjusted). Pathways are grouped by functional category: adipose metabolism and lipid handling (top), energy production and mitochondria, muscle and structural remodeling, inflammation and immunity, and cell cycle, RNA processing, and protein synthesis (bottom). See Supplementary File 1 for full enrichment results. **(D)** Concordance of protein-level and transcript-level delta-delta changes (Young recipient minus Old recipient logFC) for genes belonging to the WAT rejuvenation signature. Each point represents a gene present in both the proteome and transcriptome datasets. Pearson’s r indicates the correlation coefficient, and p indicates the statistical significance (p-value) of the observed correlation, computed among concordantly changing genes. Red points indicate muscle-related genes; yellow points indicate other concordant genes; grey points indicate discordant genes. *Atp2a1* is highlighted as a key rejuvenation-associated gene. **(E)** Pearson’s correlations of metabolite abundance changes in WAT samples after transplantation across all experimental groups. Red indicates positive correlation, and the number indicates the correlation coefficient (Pearson’s r). Asterisks reflect statistical significance (BH-adjusted p-values). **(F)** Convergent and divergent metabolomic changes induced in young-to-old and old-to-young transplantation groups. Scatter plot of metabolite-level logFC for the rejuvenation contrast (Old to young; y-axis) versus the aging contrast (Young to old; x-axis). Orange points indicate metabolites that were significantly changed in the same direction in both contrasts (concordant (shared); p.adjusted < 0.1 in both). Blue points indicate metabolites that changed significantly in the opposite direction to aging (Young to old contrast) for the rejuvenation contrast (Old to young; p.adjusted < 0.1). Grey points represent discordant or non-significantly changing metabolites for the rejuvenation contrast. A linear regression line is shown (dashed black). **(G)** Changes in identified convergent and divergent metabolites (y-axis) in each transplantation condition (x-axis). Color indicates the direction of change following transplantation (logFC; red = increased abundance, blue = decreased abundance), and an asterisk indicates statistical significance (p.adjusted). Number of samples from individual mice in both proteomic and metabolomic assays = 3 per group. *** p.adjusted< 0.001, ** p.adjusted < 0.01, * p.adjusted < 0.05, ^ p.adjusted < 0.1.

Because both the proteome and the transcriptome analyses pointed to shifts in cellular composition, we estimated adipocyte and macrophage content directly from cell-type-specific marker proteins (**Supplementary Figure 4A**). Macrophage scores increased significantly in old donor WAT after transplantation into both old and young recipients (**Supplementary Figure 4B**), while adipocyte scores decreased correspondingly (**Supplementary Figure 4C**), further confirming macrophage infiltration and a decline in adipocyte signal in the bulk proteome, and bulk and single-nuclei transcriptomic data. Slightly higher immune infiltration was observed for the WAT placed into old bodies.

A smaller set of proteins changed in opposite directions between the old-to-young and old-to-old contrasts, defining a divergent, rejuvenation-specific proteomic signature (**Figure 4B**). Proteins elevated in old-to-young grafts and lowered in old-to-old grafts included TXNIP, the metallothioneins MT1 and MT2, RAP2C, NDUFAF3, TRP53BP1, CIT, TRRAP, FAM104A, IGHV1-11, IGKV12-44, and IGHV1-82. On the other hand, CAT, S100A1, BAG2, SERPINB6C, RANBP1, and SF3B5 were lowered during rejuvenation and elevated in old-to-old grafts.

To test the relationship of such divergence with aging, we correlated the proteomic contrasts with established transcriptomic signatures of aging and mortality across tissues and species ^24^ (**Supplementary Figure 4D**). The old-to-old and old-to-young responses both correlated significantly and positively with aging and mortality signatures, consistent with their shared transplantation component, whereas the rejuvenation-specific contrast (OYΔ–OOΔ) were significantly negatively correlated, suggesting that the divergent proteome reflects a reversal of molecular age.

To resolve the pathways underlying these changes and to assess their concordance with the transcriptome, we performed gene set enrichment analysis across both transcriptomic and proteomic modalities and all contrasts (**Figure 4C, Supplementary File 1**). Adipogenesis pathway was suppressed, and inflammatory response, allograft rejection, natural killer cell mediated cytotoxicity, neutrophil degranulation, E2F targets, rRNA processing, and Nonsense Mediated Decay (NMD) independent of the EJC pathways were activated across both modalities in both old-to-young and old-to-old grafts marking them as part of the shared transplantation response. By contrast, peroxisome pathway suppression, as well as activation of myogenesis, cytoskeleton in muscle cells, and muscle contraction pathways, were specific to old-to-young transplantation relative to the old isochronic group (OYΔ–OOΔ) and were observed at both the protein and transcript modalities, resolving rejuvenation-specific cross-modality pathway signature.

Next, we asked whether the transcriptomic changes associated with rejuvenation were mirrored at the protein level (**Figure 4D**). For each gene of the WAT rejuvenation signature, we computed the difference between the old-to-young and old-to-old logFC (delta-delta) separately in the proteome and the transcriptome, such that positive values denote a greater increase after transplantation into a young than into an old host, and classified genes as concordant when the two modalities agreed in sign. Among concordant genes, protein- and transcript-level delta-deltas were strongly correlated (Pearson’s r = 0.74). The concordant set was dominated by muscle-related genes, including *Atp2a1* (SERCA1), *Cacna1s*, *Myom2*, *Myoz1*, *Myh1*, *Mybpc1*, *Mybpc2*, *Tnnt3*, *Tnni2*, *Mylpf*, *Ttn*, *Obscn*, *Klhl40*, *Klhl41*, *Sgce*, and *Tcap*, together with *Ckmt2*, *Slc5a3*, and *Mgst3*, all of which were positive in both modalities. *Lrmda*, *Fabp4*, *Nap1l5*, *Negr1*, *Nr1d1*, *Gulp1*, *Cgnl1*, *Fggy*, *Ormdl3*, and the muscle-related *Ifrd1* and *Gsn* were concordantly negative.

### Metabolomic signature of WAT rejuvenation

We next profiled the WAT metabolome across all four transplantation conditions (sample n = 3 per group), quantifying 320 metabolites after quality filtering and internal standard removal. Similar to the transcriptome and proteome, metabolite changes after transplantation were positively and significantly correlated across all groups (**Figure 4E**), although here the weakest correlation was between old-to-young and young-to-young grafts (Pearson’s r = 0.48) and the strongest between the two isochronic groups (Pearson’s r = 0.70).

To identify the metabolic changes accompanying rejuvenation, we performed moderated differential analysis of the old-to-young contrast, which yielded 14 metabolites of significantly altered abundance (p.adjusted < 0.10). Eleven were phospholipids: four phosphatidylglycerols (PG(36:0), PG(36:4), PG(38:5), PG(40:8)), two phosphatidylethanolamines (PE(32:0), PE(34:0)), two phosphatidylserines (PS(32:0), PS(34:0)), and one phosphatidylcholine (PC(36:1)). The remaining five were cysteine and its oxidation product cysteic acid, cyclic AMP, sedoheptulose, and glucuronic acid.

To determine which of these changes were specific to the rejuvenation, we classified each of the 16 metabolites by comparing their behavior in the old-to-young and young-to-old transplantation conditions (**Figure 4F**). 11 of the 14 changed significantly (p.adjusted < 0.10) and in the same direction in both contrasts: nine increased (PE(32:0), PE(34:0), PG(36:4), PG(38:5), PG(40:8), PS(32:0), PS(34:0), cysteine, and cysteic acid) and two decreased (PC(36:1) and PG(36:0)). We classified these as a shared transplant response. The remaining three, sedoheptulose, glucuronic acid, and cyclic AMP, changed in the opposite direction during old-to-young transplantation and were classified as divergent.

Mapping the 14 metabolites across all four transplantation conditions further confirmed this distinction (**Figure 4G**). All shared metabolites also changed significantly (p.adjusted < 0.10) and in the same direction in old-to-old grafts, suggesting they reflect a response to the transplantation procedure. The three divergent metabolites changed significantly (p.adjusted < 0.10) in no arm other than old-to-young, with increasing sedoheptulose, glucuronic acid, and cyclic AMP defining a rejuvenation-associated metabolic signature.

To summarize, the proteome and metabolome corroborate and extend the transcriptomic signature of WAT rejuvenation.

### WAT rejuvenation model and validation of findings at the structural level of organization

To synthesize the molecular features of rejuvenation into a single framework, we integrated the transcriptomic, proteomic, and metabolomic changes observed in old adipose tissue transplanted into young recipients into a working model by conducting an extensive analysis of the existing literature (**Figure 5A**). Features were grouped by the biological process they inform, colored by their direction of change, and, where available, by their concordance across the RNA and protein levels. The model places activated thermogenesis, including futile calcium and creatine fluxes, at the center of the rejuvenation program. In addition, our working model identifies the suppression of differentiation and glutathione metabolism as downstream consequences of thermogenic programs activation. Further, we observe downregulation of genes and proteins positively associated with various adipose tissue disorders. Lastly, our findings point to activation of cellular homeostasis maintenance programs, driven by upregulated genes and proteins responsible for providing structural support and DNA repair within the cells. **Table 1** lists these features and describes their functions and roles during WAT rejuvenation.

**Figure 5.**
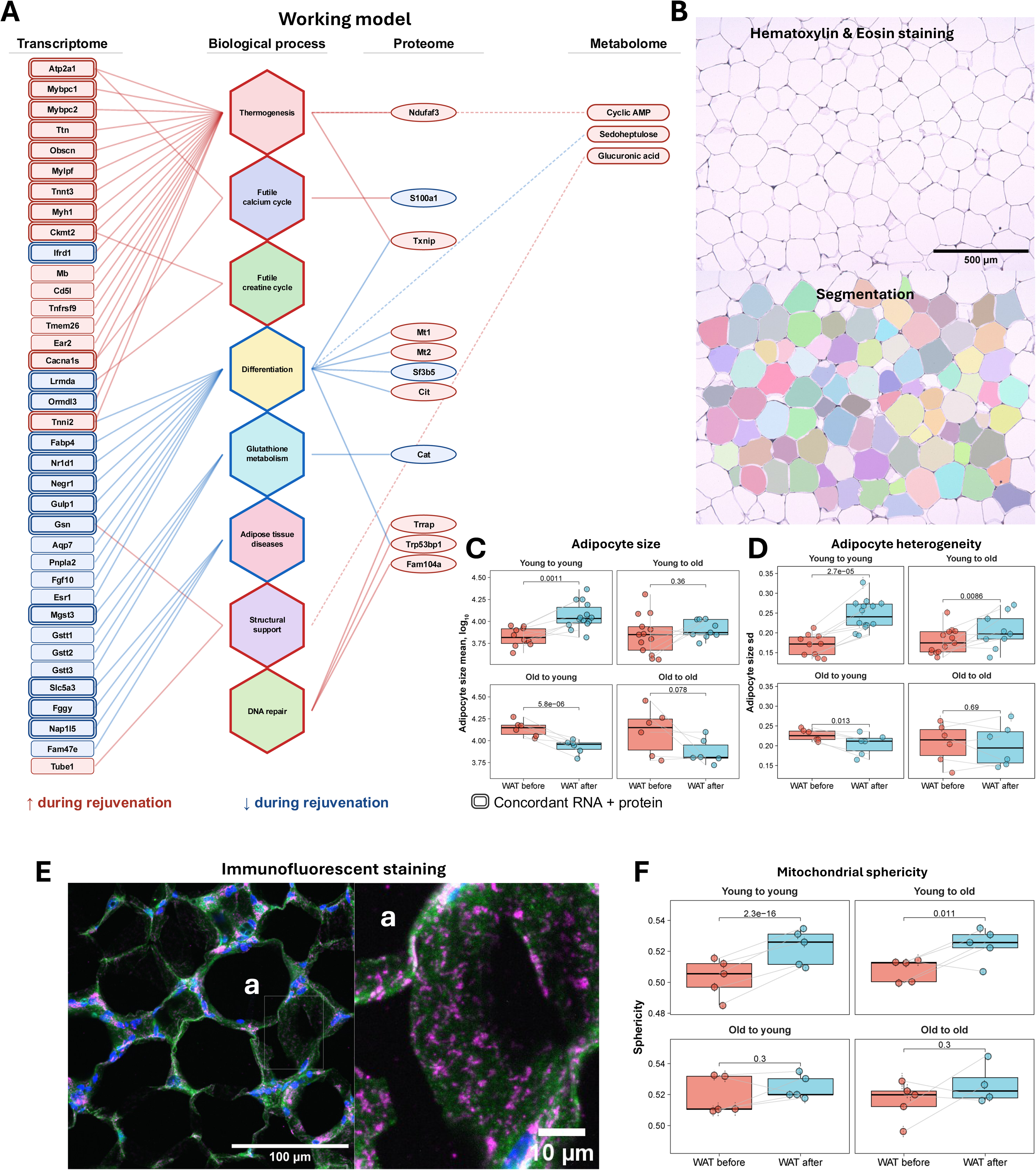
WAT rejuvenation model and validation of findings at the structural level of organization. **(A)** Integrative working model of molecular changes underlying WAT rejuvenation, combining transcriptomic, proteomic, and metabolomic findings. Columns represent distinct omics layers (transcriptome, proteome, metabolome), connected to their respective biological process. Node colors indicate the direction of change in old WAT transplanted into young recipients: red, increased; blue, decreased. Transcriptomic nodes with a double border indicate features with concordant changes at both the RNA and protein levels. Biological processes are organized by functional domain, encompassing thermogenesis, futile calcium and creatine cycles, differentiation, glutathione metabolism, adipose tissue diseases, structural support, and DNA repair. **(B)** Example hematoxylin and eosin (H&E)-stained section of inguinal white adipose tissue (top) and selected adipocytes used for lipid droplet size quantification (bottom). **(C)** Quantification of adipocyte lipid droplet size ratio (in log scale) in WAT samples before and after transplantation across all experimental groups. Differences in mean droplet size before and after transplantation were assessed with mixed-effects models. Mouse ID was included in the statistical model as a covariate to ensure paired comparison of lipid droplet size ratios. Data are mean droplet size (in log scale) ± s.e.m. Each dot represents a sample collected from individual mice, n = 6-13 per group. **(D)** Quantification of adipocyte lipid droplet size (in log scale) standard deviation in all WAT samples before and after transplantation across all experimental groups. Differences in droplet size standard deviation before and after transplantation were assessed with mixed-effects models. Mouse ID was included in the statistical model as a covariate to ensure paired comparison of lipid droplet size standard deviation ratios. Data are mean standard deviation of adipocyte size (in log scale) ± s.e.m. Each dot represents a sample collected from individual mice, n = 6-13 per group. **(E)** Immunofluorescent staining of 15 μm adipose tissue sections for mitochondria (Tomm20, magenta), adipocyte cell membrane (autofluorescence, green) and nuclei (DAPI, blue); panel (a) shows higher magnification highlighting mitochondrial labeling. **(F)** Quantification of mitochondrial sphericity in WAT samples before and after transplantation across all experimental groups. Changes of mitochondrial sphericity following transplantation were assessed with mixed-effects models. Mouse ID was included in the statistical model as a covariate to ensure paired comparison of mitochondrial sphericities. Data are mean ± s.e.m. Each dot represents a sample collected from individual mice, n = 4-5 per group. Boxplots: center line indicates median; box limits, interquartile range; whiskers, ±1.5× IQR. Unless specified otherwise, statistical differences between groups are assessed with ANOVA and adjusted for multiple comparisons with the Benjamini-Hochberg approach. *** p.adjusted< 0.001, ** p.adjusted < 0.01, * p.adjusted < 0.05, ^ p.adjusted < 0.1.

**Table 1.**
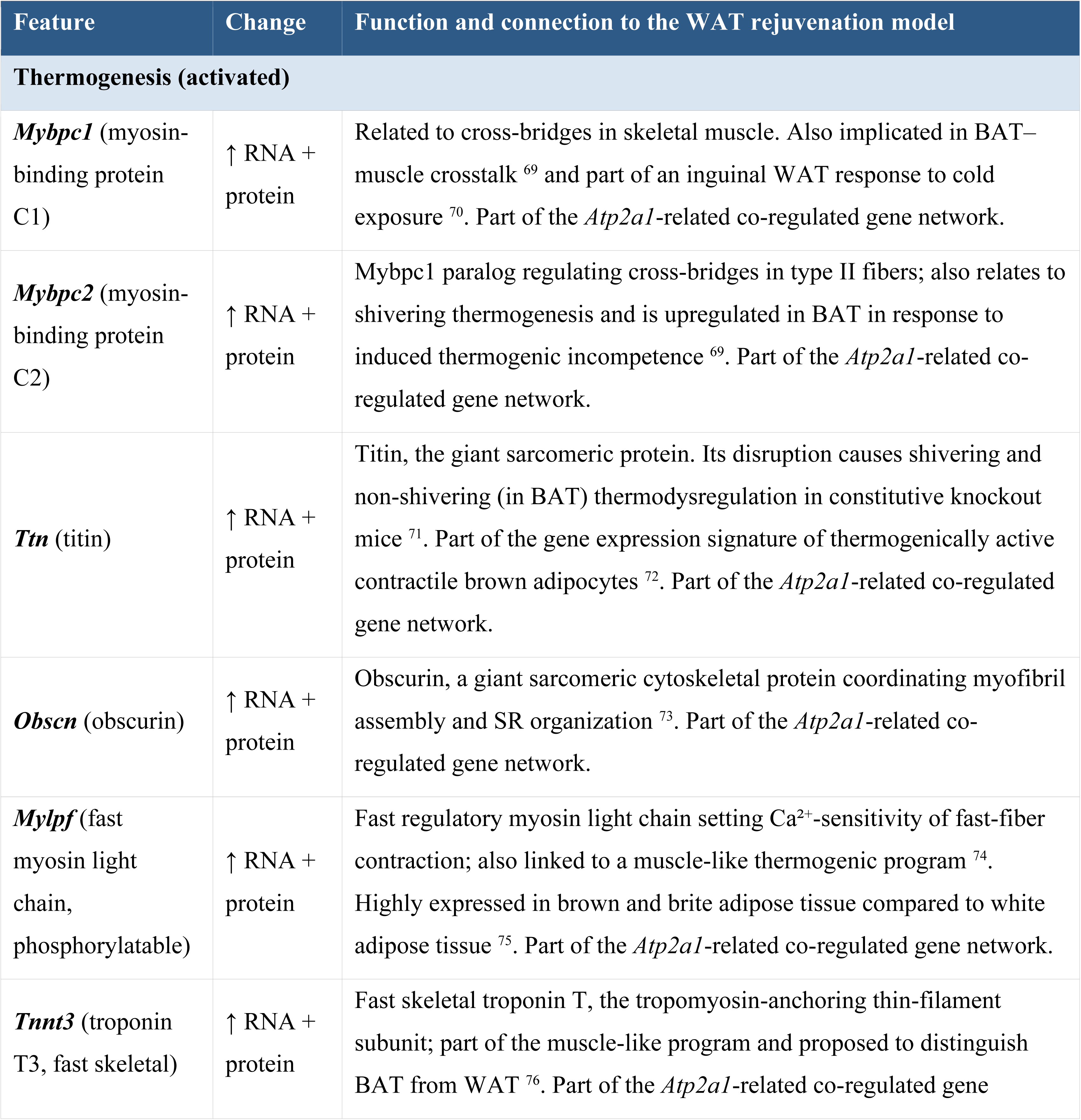
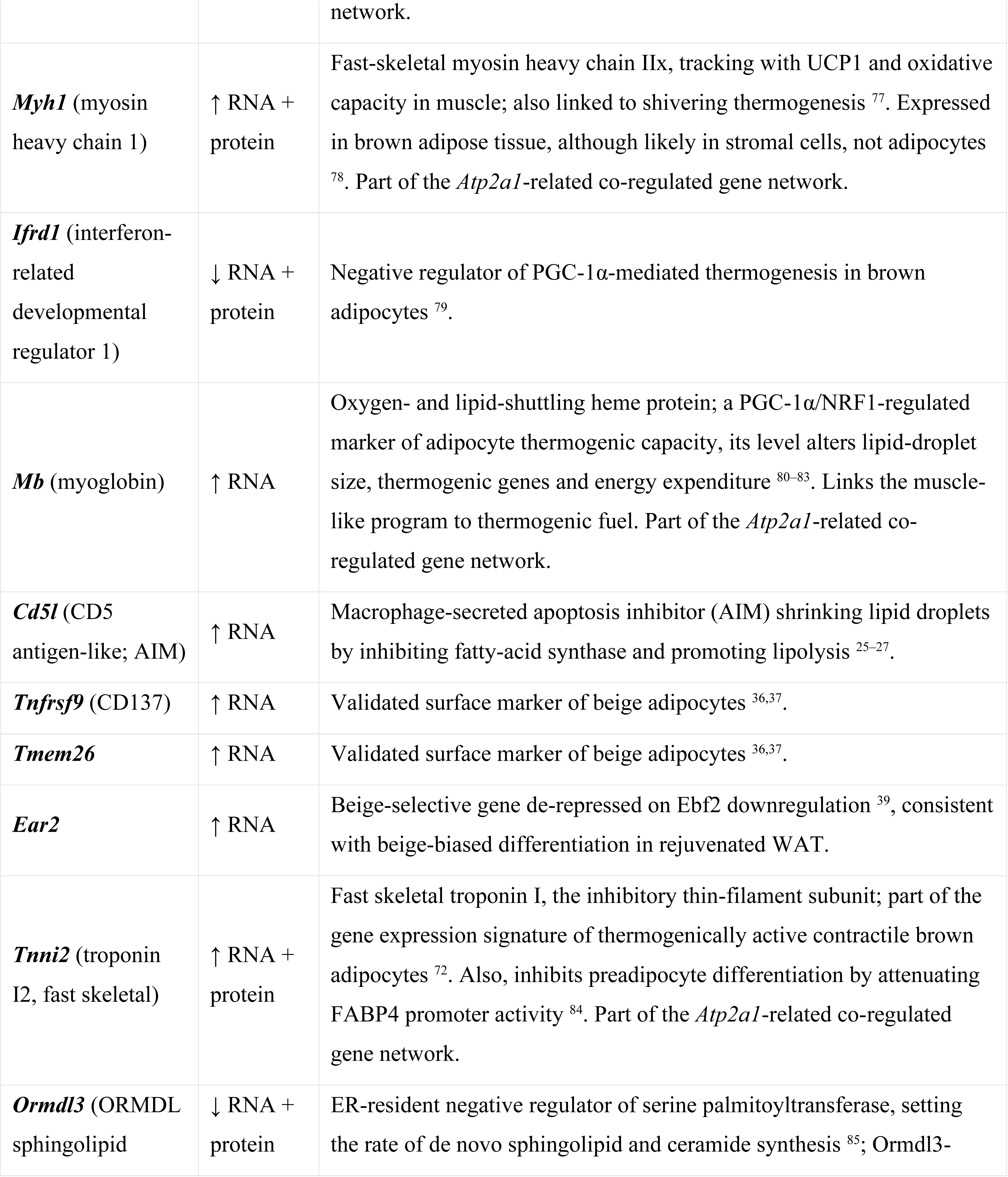
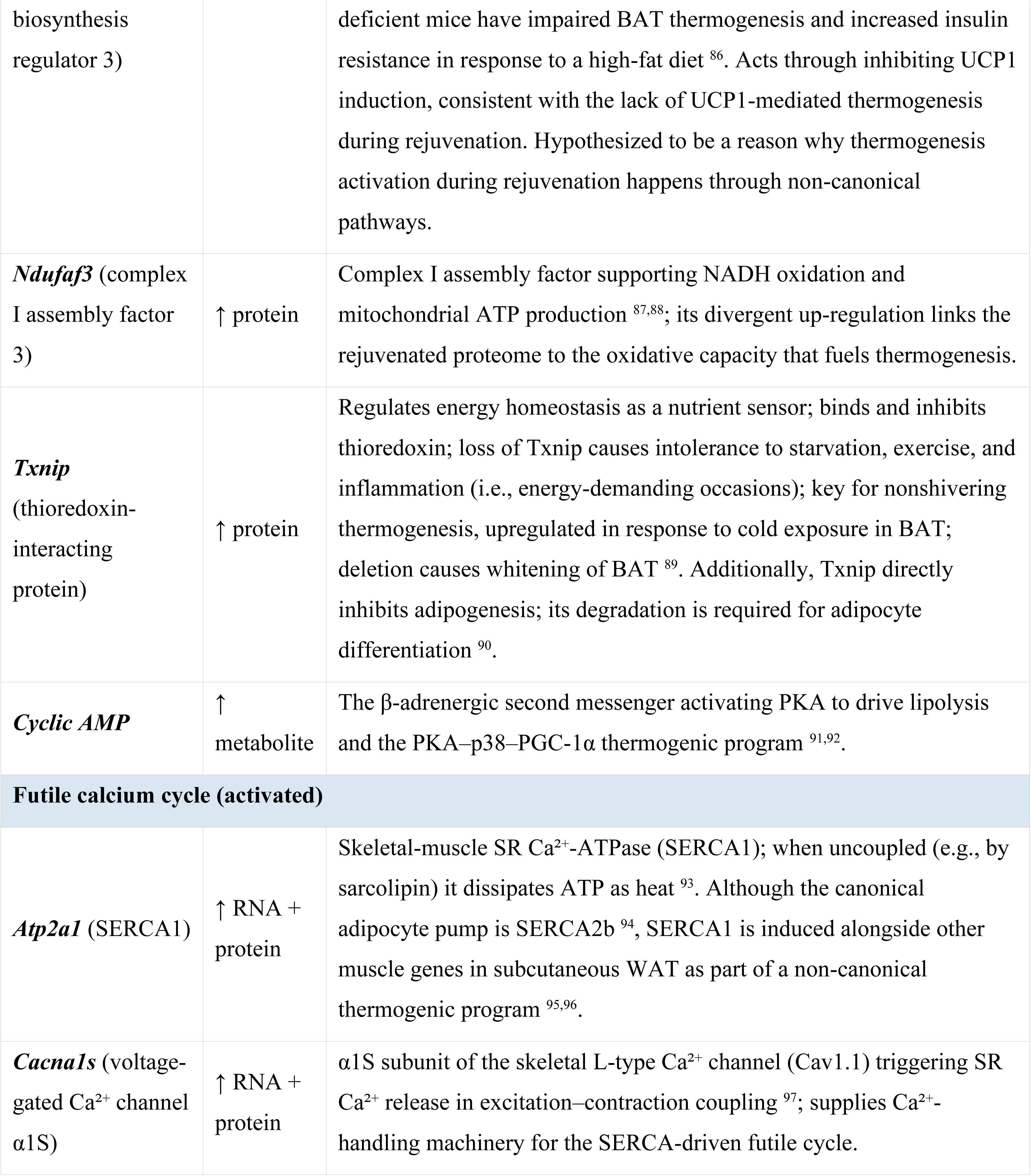
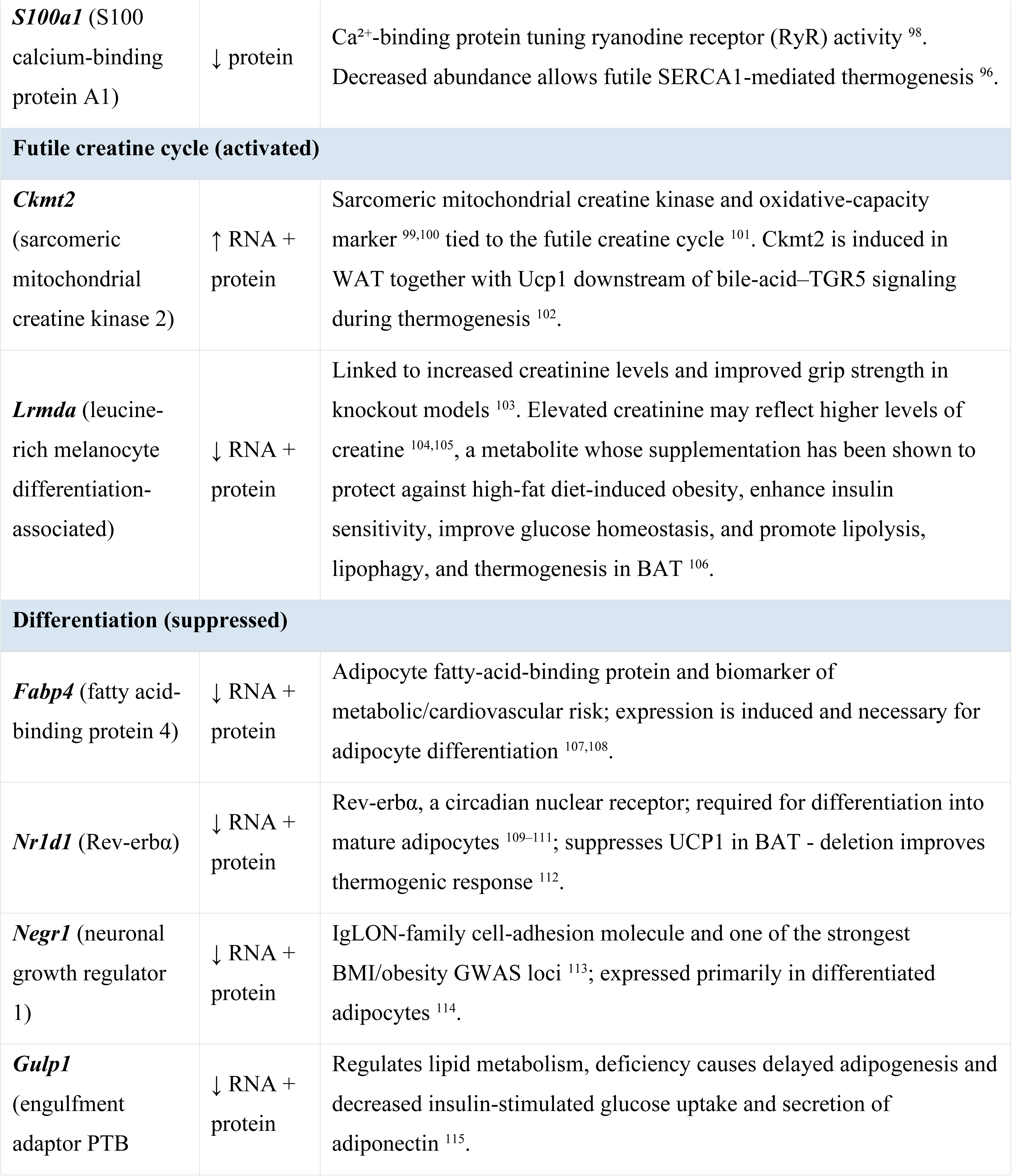
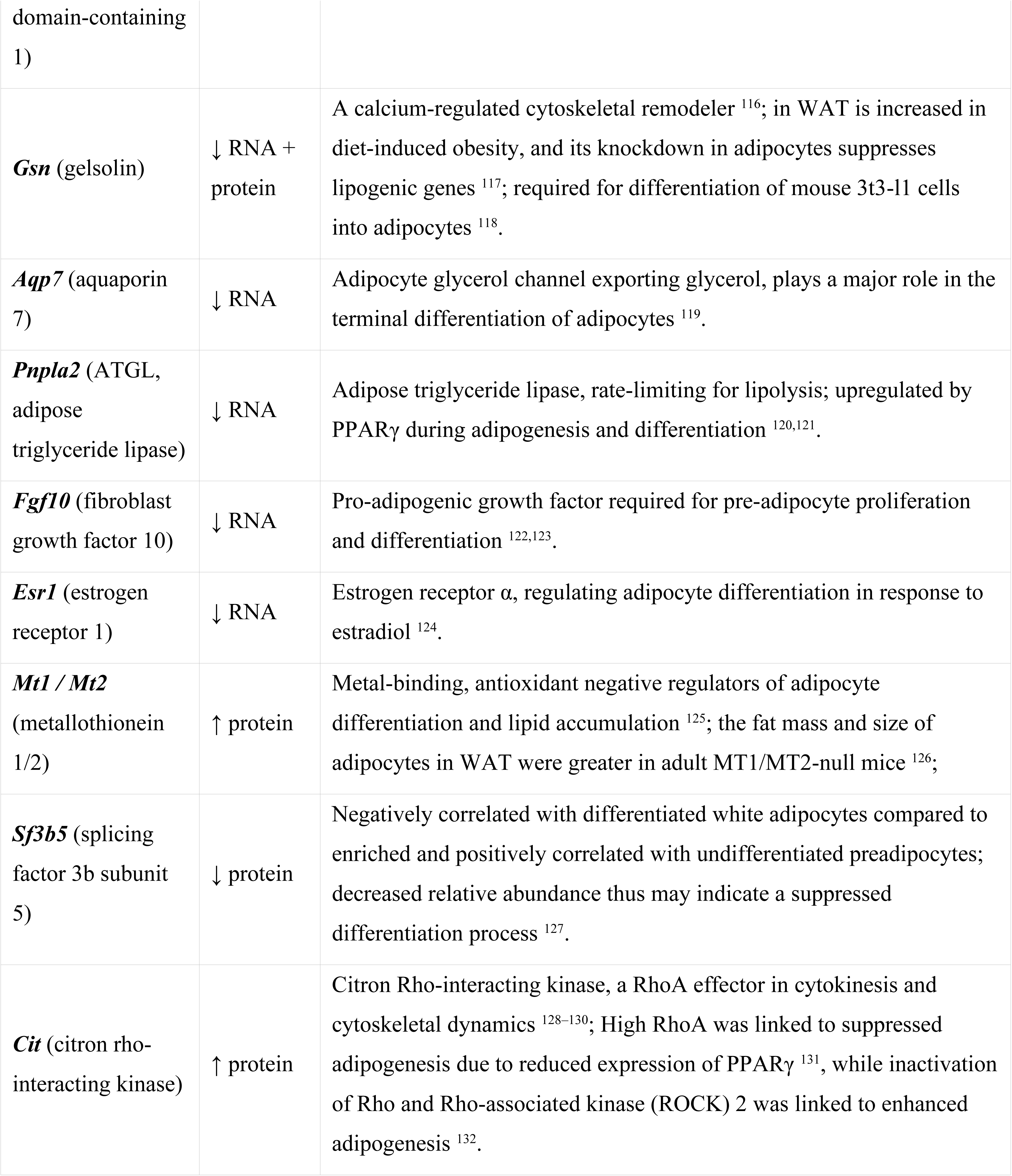
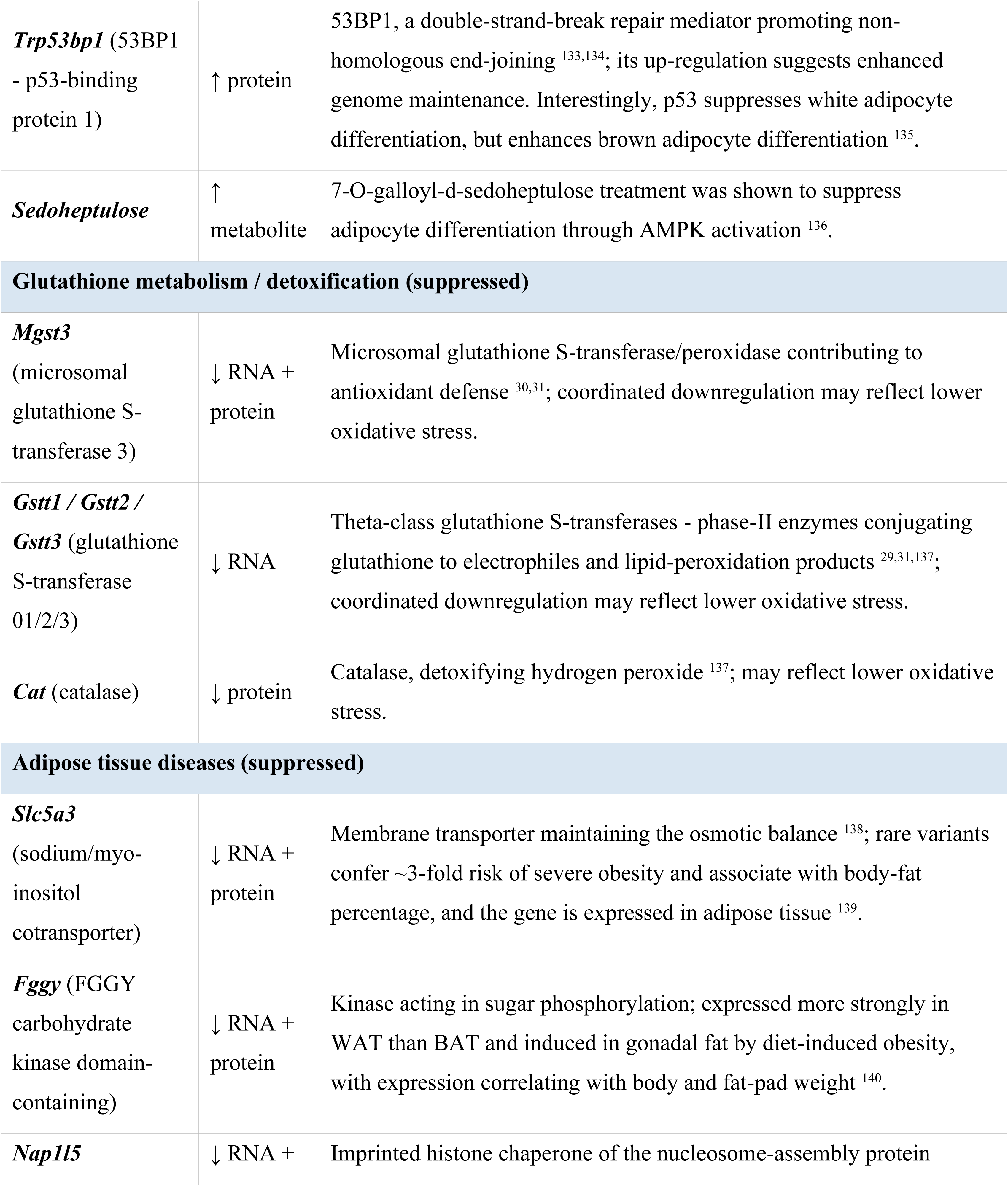
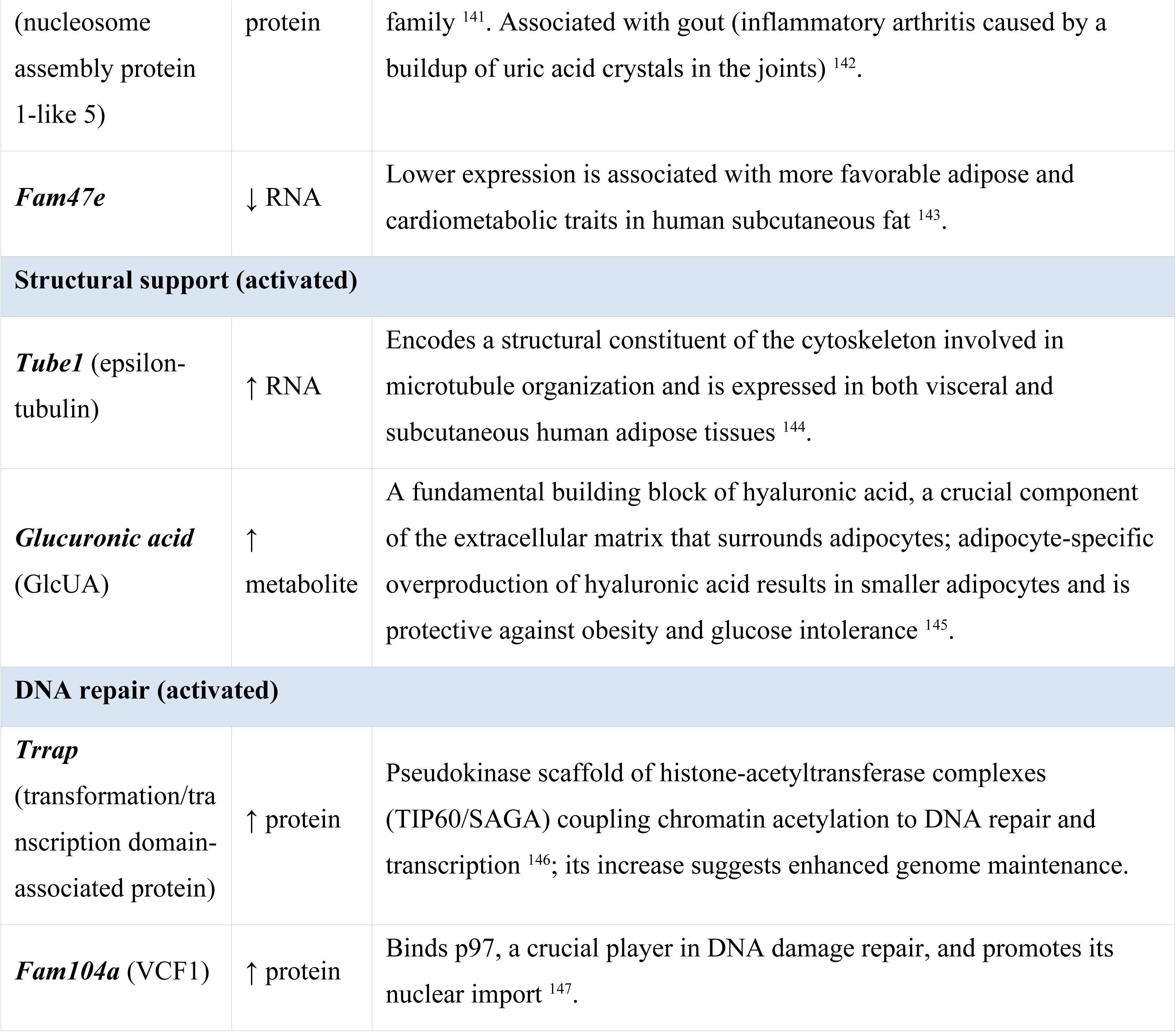
Molecular features of WAT rejuvenation. *Direction of change refers to old WAT transplanted into young recipients (rejuvenation). “RNA + protein” indicates features concordant at both the transcript and protein level (double-bordered nodes in* ***Figure 5A)***.

Given that the rejuvenation-associated genes, proteins, and metabolites were highly involved in lipid droplet size regulation and activation of thermogenesis, we decided to validate our findings by assessing adipocyte lipid droplet sizes across transplantation groups. Specifically, we performed Segment Anything Model (SAM)-based quantification of lipid droplet sizes from Hematoxylin and eosin (H&E)-stained sections of WAT collected before and after transplantation (**Figure 5B**). Consistent with prior reports that aging is associated with adipocyte hypertrophy ^46^, baseline measurements showed that old WAT generally exhibited larger lipid droplets compared to young WAT, although some variability was present (**Figure 5C**). Following transplantation, we observed an increase in lipid droplet size in young adipocytes implanted into young recipients, and conversely, a significant reduction in droplet size in old adipocytes implanted into young hosts. At the same time, an increase in size for young adipocytes implanted into old recipients did not reach statistical significance, and old adipocytes implanted into old recipients exhibited a marginally significant decrease in lipid droplet size.

We next quantified intra-tissue heterogeneity in adipocyte size, using the standard deviation of lipid droplet area (in log scale) as a proxy. Overall, variation in adipocyte size mirrored the patterns observed for mean lipid droplet size, where old adipose tissue had a higher cellular heterogeneity compared to young. Young grafts in young recipients exhibited increased heterogeneity following the transplantation (**Figure 5D**), consistent with expansion of the largest adipocyte subpopulations and a possible shift toward a hypertrophic phenotype. In contrast, old adipose tissues transplanted into young hosts showed a marked reduction in heterogeneity, indicating not only a global reduction in adipocyte size but also its stabilization across the population. Young adipose tissue transplanted into old recipients displayed increased adipocyte size heterogeneity, suggesting the emergence of hypertrophic subpopulations even in the absence of significant mean size change. Lastly, the old-to-old group showed no changes in heterogeneity.

To further characterize tissue remodeling following transplantation, we evaluated collagen deposition in WAT using Masson’s Trichrome staining (**Supplementary Figure 5A)**. Elevated collagen levels have been linked to macrophage recruitment and adipose pathologies such as diet-induced obesity, hypoxia, and fibrosis ^47–50^. Post-transplantation, collagen levels increased significantly across all groups except the young-to-young controls, and the largest increase in collagen content was observed in old grafts implanted into young hosts (**Supplementary Figure 5B**). The analysis of collagen content revealed that young tissues contained higher baseline levels of collagen than old tissues prior to transplantation, suggesting that collagen accumulation during fibrosis may reflect adaptive remodeling rather than degenerative changes. Alternatively, the apparent increase could result from enhanced visibility of collagen in regions with smaller or less densely packed adipocytes.

Fibrosis increased post-transplantation in all groups, but changes were not significantly different between any of the compared transplantation types (**Supplementary Figure 5C**). Overall, the strongest fibrotic response occurred in young-to-old grafts, while old-to-young grafts showed the least increase. These findings suggest that transplantation broadly activates fibrotic remodeling, with host age exerting a modest influence on its extent.

We next examined whether mitochondrial abundance and morphology were altered in WAT following transplantation (**Supplementary Figure 5D)**. Mitochondrial fragmentation and increased sphericity are established features of aging ^51^, although recent studies suggest that increased fragmentation can be uncoupled from aging, potentially representing changes that are neither adaptive nor deleterious, but instead reflect alternative cellular processes ^52^. 15 μm-thick tissue sections used for quantitative analysis (**Figure 5E**) revealed that old and young adipocytes before transplantation had comparable numbers of mitochondria, and mitochondrial abundance did not significantly change post-transplantation across any transplantation groups, regardless of donor or host age (**Supplementary Figure 5E**). Mitochondria in young adipocytes on average exhibited greater volumes and lower sphericity, consistent with prior observations of more elongated and connected mitochondrial networks in young cells ^52^. Upon transplantation into either young or old hosts, young adipocytes displayed increased mitochondrial sphericity (**Figure 5F**) and reduced volume (**Supplementary Figure 5F**), indicative of enhanced mitochondrial fragmentation. In contrast, old adipocytes transplanted into young or old recipients showed no significant changes in mitochondrial volume or shape, suggesting that age-related changes in mitochondrial morphology are resistant to rejuvenation. These findings imply that only select age-related features, potentially those most relevant to function, are reset by transplantation, while others may remain unaffected, at least within the timeframe of the conducted study.

Collectively, our results position a non-canonical thermogenic program as a central feature of adipose rejuvenation, with reduced adipocyte lipid droplet size as its principal structural consequence.

### Heterochronic WAT transplantation has a limited impact on host systemic aging

Having characterized rejuvenation within the graft, we asked whether heterochronic transplantation reciprocally affected the systemic aging of the host. We estimated the biological age of three host compartments - the recipient’s native WAT, liver, and blood (**Figure 1A**). Native WAT was not present in donors at the end of the study, so only liver and blood were characterized for this group. Donor and recipient animals were compared against age-matched sham-operated controls, and recipients of old and young adipose tissue were additionally compared against each other. Comparisons were done using a panel of epigenetic (Universal pan-tissue Clocks 2 and 3, SVM Universal Mortality pan-tissue Clock) and transcriptomic (chronological and mortality rodent multi-tissue) clocks. Molecular clock analyses extended to comparing complete blood count (CBC) compositions, blood-based dynamic frailty indicator (dFI) and its embedding modifications ^53^, and body weight. For every clock and dFI embedding, comparisons were done for residuals computed after regressing out chronological age, study, and sequencing batch when present so that positive values of molecular clocks denote age acceleration beyond that expected for the animal’s chronological age, and the opposite is true for the values below zero.

In the young recipients’ native WAT, the SVM Universal Mortality pan-tissue Clock detected epigenetic age (eAge) acceleration of both recipients relative to sham (**Figure 6A**). The effect was stronger in young recipients of old fat, but no statistically significant difference was detected between recipients. No difference was observed using Universal pan-tissue Clocks 3 and 2 (**Supplementary Figure 6A, 6B**). Similarly, WAT Native of both old recipients had elevated eAge compared to shams according to SVM Universal Mortality pan-tissue Clock and Universal pan-tissue Clock 2, with no differences observed for Universal pan-tissue Clock 3 (**Figure 6B; Supplementary Figure 6C, 6D**). Surprisingly, old recipients of old fat had significantly lower eAge compared to old recipients of young fat according to the SVM Universal Mortality pan-tissue Clock (**Figure 6B)**. Lastly, both Chronological and Mortality Transcriptomic Clocks did not detect any differences between Native WATs of any groups (**Supplementary Figure 6E, 6F, 6G, 6H**).

**Figure 6.**
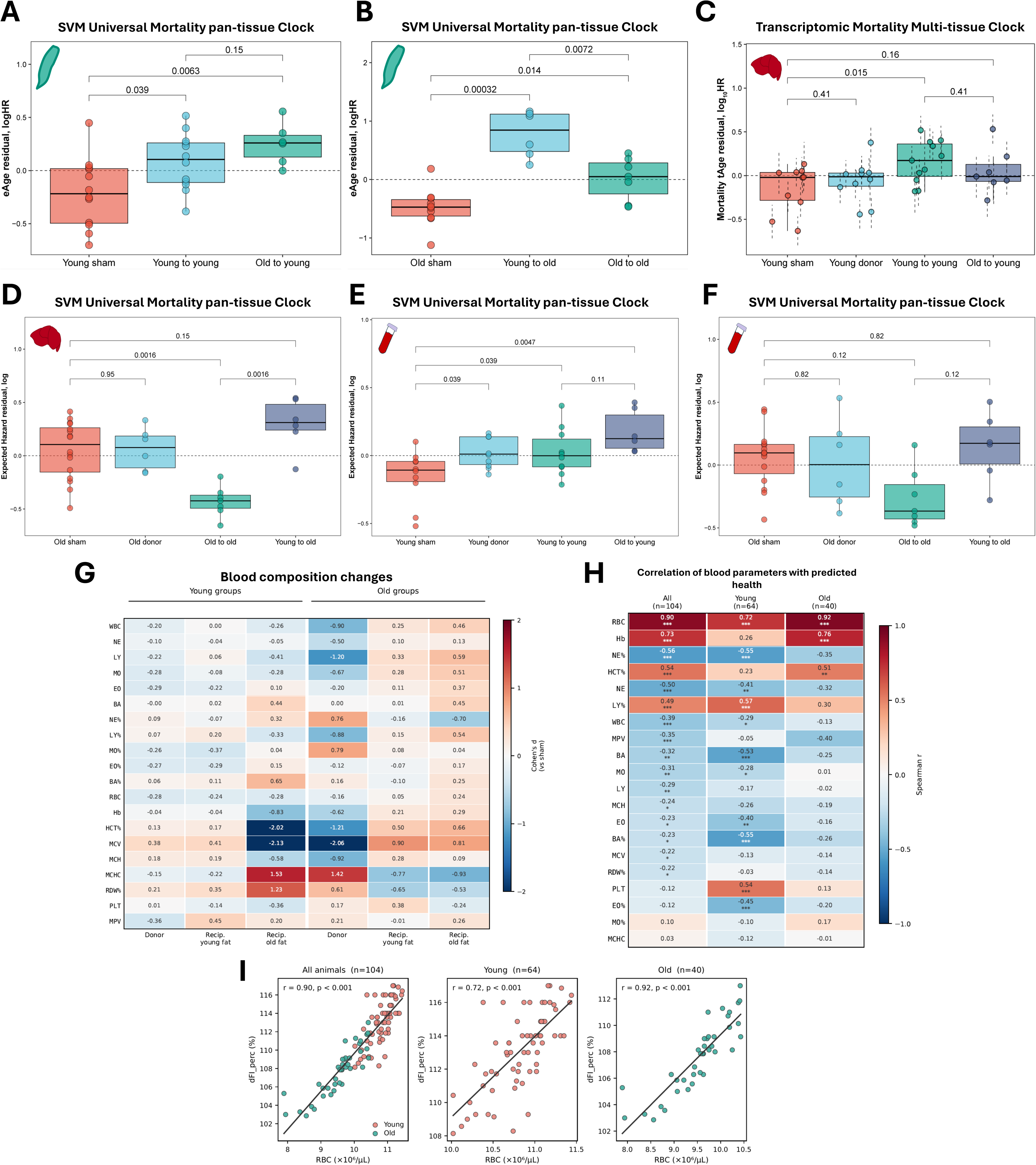
Heterochronic WAT transplantation has a limited impact on host systemic aging. **(A)** Expected hazard ratio residuals (in log scale) (eAge residual, logHR) of native WATs from young recipient mice (young-to-young and old-to-young transplantation groups) compared to young sham controls, estimated using SVM Universal Mortality pan-tissue Clock. Residuals were obtained by regressing out chronological age, sequencing batch, and study as covariates; positive values indicate epigenetic age acceleration relative to the value expected for the given chronological age. BH-adjusted p-values reflect pairwise comparisons between groups. Each dot represents a sample collected from individual mice, n = 6-15 per group. **(B)** eAge expected hazard ratio residuals (in log scale) (eAge residual, logHR) of native WATs from old recipient mice (young-to-old and old-to-old transplantation groups) compared to old sham controls, estimated using SVM Universal Mortality pan-tissue Clock. Residuals were obtained by regressing out chronological age, sequencing batch, and study as covariates; positive values indicate epigenetic age acceleration relative to the value expected for the given chronological age. BH-adjusted p-values reflect pairwise comparisons between groups. Each dot represents a sample collected from individual mice, n = 6-10 per group. **(C)** tAge expected hazard ratio residuals (in log scale) (Mortality tAge residual, log_10_HR) of livers from young donor and recipient mice (young-to-young and old-to-young transplantation groups) compared to young sham controls, estimated using Bayesian Ridge (BR) rodent multi-tissue mortality clock. Residuals were obtained by regressing out chronological age, sequencing batch, and study as covariates; positive values indicate transcriptomic age acceleration relative to the value expected for the given chronological age. BH-adjusted p-values reflect pairwise comparisons between groups, assessed with mixed-effects models. Each dot represents a sample collected from individual mice, n = 7-11 per group. **(D)** Expected hazard ratio residuals (in log scale) (eAge residual, logHR) of livers from old donor and recipient mice (young-to-old and old-to-old transplantation groups) compared to old sham controls, estimated using SVM Universal Mortality pan-tissue Clock. Residuals were obtained by regressing out chronological age and study as covariates; positive values indicate epigenetic age acceleration relative to the value expected for the given chronological age. BH-adjusted p-values reflect pairwise comparisons between groups. Each dot represents a sample collected from individual mice, n = 6-16 per group. **(E)** Expected hazard ratio residuals (in log scale) (eAge residual, logHR) of blood from young donor and recipient mice (young-to-young and old-to-young transplantation groups) compared to young sham controls, estimated using SVM Universal Mortality pan-tissue Clock. Residuals were obtained by regressing out chronological age and study as covariates; positive values indicate epigenetic age acceleration relative to the value expected for the given chronological age. BH-adjusted p-values reflect pairwise comparisons between groups. Each dot represents a sample collected from individual mice, n = 6-10 per group. **(F)** Expected hazard ratio residuals (in log scale) (eAge residual, logHR) of blood from old donor and recipient mice (young-to-old and old-to-old transplantation groups) compared to old sham controls, estimated using SVM Universal Mortality pan-tissue Clock. Residuals were obtained by regressing out chronological age and study as covariates; positive values indicate epigenetic age acceleration relative to the value expected for the given chronological age. BH-adjusted p-values reflect pairwise comparisons between groups. Each dot represents a sample collected from individual mice, n = 6-15 per group. **(G)** Heatmap of Cohen’s d values for 20 CBC parameters comparing each intervention group against the biologically appropriate sham reference (Young sham for young groups, Old sham for old groups). Positive values (red) indicate higher values in the intervention group relative to sham; negative values (blue) indicate lower values. Cell values show Cohen’s d; asterisks indicate BH-adjusted p-values. All comparisons were performed using ANCOVA with study batch and chronological age as covariates. Mice per group: young sham = 17; young donor = 15; young recipient of young fat = 20; young recipient of old fat = 12; old sham = 15; old donor = 11; old recipient of young fat = 8; old recipient of old fat = 6. **(H)** Heatmap of Spearman rank correlation coefficients (r) between each of the 20 CBC parameters and dFI_perc, computed for all animals combined (n = 104), young animals only (n = 64), and old animals only (n = 40). Parameters are sorted by absolute r in the all-animals subset (largest at the top). Red indicates positive correlation, blue indicates negative correlation. Cell values show r; asterisks indicate BH-adjusted p-values. Column headers indicate subset and sample size. Mice per group same as in G. **(I)** Scatter plots of red blood cell count (RBC, ×10⁶/µL) against dFI_perc (%) for (left) all animals (n = 104), (center) young animals only (n = 64), and (right) old animals only (n = 40). Individual data points are colored by age group (young: salmon, old: teal). The overlaid line shows the ordinary least-squares regression fit. Spearman r and BH-adjusted p-value are displayed in the upper left of each panel. Mice per group same as in G. Boxplots: center line indicates median; box limits, interquartile range; whiskers, ±1.5× IQR. Unless specified otherwise, statistical differences between groups are assessed with ANOVA and adjusted for multiple comparisons with the Benjamini-Hochberg approach. *** p.adjusted< 0.001, ** p.adjusted < 0.01, * p.adjusted < 0.05, ^ p.adjusted < 0.1

Adipose tissue implantation or removal resulted in small and inconsistent shifts in liver biological age. In particular, no differences were observed across any epigenetic clocks applied to livers of tested young groups (**Supplementary Figure 6I, 6J, 6K**). At the same time, both mortality and chronological transcriptomic multi-tissue clocks indicated that tAge of young recipients of young fat livers was higher relative to young sham (**Figure 6C**; **Supplementary Figure 6L**). No other groups were different from each other. Analysis of old groups showed that livers of old recipients of old fat had significantly lower eAge compared to sham-operated animals and old recipients of young fat, according to the SVM Universal Mortality pan-tissue Clock (**Figure 6D**). Further, the old-to-old group had significantly lower eAge compared to shams, and marginally significant eAge reduction compared to old recipients of young fat, according to Universal pan-tissue Clock 3 (**Supplementary Figure 6M**). A similar, albeit non-significant for any comparison, trend was observed using Universal pan-tissue Clock 2 (**Supplementary Figure 6N**). No differences in old liver tAge were observed using Transcriptomic Clocks (**Supplementary Figure 6O, 6P**).

In blood, the SVM Universal Mortality Clock detected eAge elevation in every young animal intervention group (**Figure 6E**). Young recipients of old fat experienced the strongest aging-accelerating effect, potentially due to a release of old immune cells from the old graft, but no statistically significant difference was detected compared with the young-to-young group. Elevated eAge residuals compared to shams were observed for the young recipients of old fat group using Universal pan-tissue Clocks 3 and 2 as well (**Supplementary Figure 6Q, 6R**). Blood eAge of old animals mirrored the eAge predictions for livers, although the effect was weaker and reached only marginal statistical significance for the eAge reduction in the old recipients of the old fat group compared to sham-operated controls according to the Universal pan-tissue Clock 2 (**Figure 6F**, **Supplementary Figure 6S, 6T**).

Next, we assessed hematological changes induced by adipose tissue implantation or removal by analyzing complete blood counts (CBC) of animals collected at study endpoint. First, we applied principal component analysis (PCA) to the CBC data to identify and remove potential outliers (**Supplementary Figure 7A**). PC1 was dominated by immune-cell parameters, and PC3 by red-blood-cell parameters, whereas PC2 captured both.

Comparison of hematological parameters revealed that the most pronounced changes (|Cohen’s d| > 1) were in young recipients of old fat and old donors (**Figure 6G**). Young recipients of old fat had reduced mean corpuscular volume (MCV) and hematocrit (HCT) as well as increased red cell distribution width (RDW) and mean corpuscular hemoglobin concentration (MCHC) relative to sham-operated controls. Old donors had a lower absolute lymphocyte count, lower HCT and MCV, and elevated MCHC relative to shams. None of these changes reached significance after correction for multiple comparisons likely owing to the limited number of animals per group. At the same time, the lack of profound blood composition changes implies that the observed effects on blood eAges are not explained by cell composition changes and may reflect intrinsic changes in cellular aging.

To assess whether observed blood composition changes corresponded to a shift in predicted health, we applied a blood-based dynamic frailty indicator (dFI) ^53^ and related predictors that we derived from dFI embeddings. To choose the predictor most sensitive to age-related changes we tested 4 versions of dFI for their ability to distinguish between young and old sham controls: original dFI (**Supplementary Figure 7B**), dFI_age, trained to predict chronological age (**Supplementary Figure 7C**), dFI_loghaz, trained to predict the logarithm of hazard (**Supplementary Figure 7D**), and dFI_perc, trained to predict the hazard percentile (**Supplementary Figure 7E**). dFI_age showed no significant difference between old and young shams, although old shams had slightly higher predicted values. The strongest separation was observed for dFI_perc (Cohen’s d = - 2.19), with old animals having lower values; dFI_perc was therefore used for subsequent comparisons of donor and recipient animals against shams, and of recipients against each other.

No significant differences were observed within young (**Supplementary Figure 7F**) or old (**Supplementary Figure 7G**) groups using dFI_perc. Among the 20 measured CBC parameters, red blood cell count (RBC) was the strongest correlate of dFI_perc, with correlations of Spearman r = 0.90 across all animals, r = 0.72 in young animals, and r = 0.92 in old animals (**Figure 6H, 6I**), identifying age-related decline in RBC as the dominant hematological component of this health metric. No other CBC parameter significantly correlated with predicted health across both young and old groups, although many of them correlated in the same direction: higher hemoglobin (Hb), HCT, lymphocyte percentage (LY%), and platelet numbers (PLT), as well as lower neutrophil percentage (NE%) and absolute number (NE), basophil and eosinophil percentage (BA%, EO%), and their absolute number (BA, EO) were highly significantly (p < 0.01) correlated with better predicted health in at least one of the age groups. Finally, body weight changes over the 10-week study period did not differ between young (**Supplementary Figure 7H**) and old (**Supplementary Figure 7I**) groups.

In sum, our assessment of recipient health metrics indicates that, in contrast to the pronounced changes within grafts themselves, the systemic aging trajectory of the host was largely unaffected by the presence of a heterochronic graft.

### Rejuvenation-associated genes *Lrmda* and *SERCA* modulate lifespan in Drosophila

To test whether genes linked to rejuvenation of adipose tissue in our transplantation model may also modulate longevity, we manipulated two of the top conserved genes exclusively in the fat bodies of *Drosophila melanogaster*. Specifically, we utilized the inducible, whole-body fat-body GeneSwitch-GAL4 (WB-FB-GS-GAL4) system activated by RU-486 feeding in adult flies ^54,55^ to knockdown *Lrmda* (downregulated during rejuvenation), knockdown *SERCA* (the single *Drosophila* SERCA orthologue, upregulated during rejuvenation), and overexpress *SERCA*, assessing survival in each sex by Kaplan-Meier, quantile regression (90th percentile) for maximum lifespan, Cox proportional-hazards, and time-varying hazard analyses (**Figure 7A**; flies per experimental condition = 97-204; vials per condition = 5-9).

**Figure 7.**
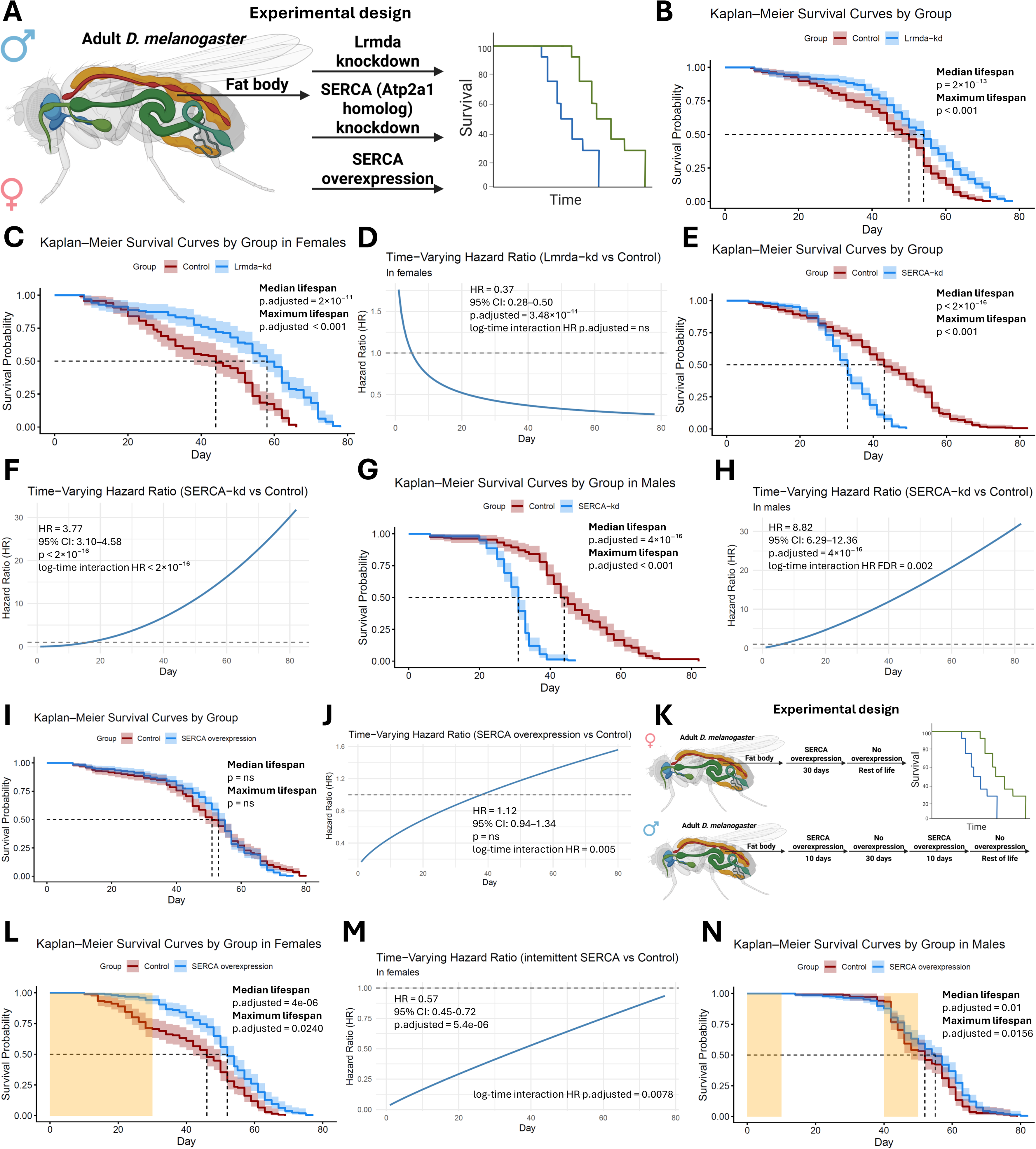
Rejuvenation-associated genes *Lrmda* and *SERCA* modulate lifespan in Drosophila. **(A)** Schematic of fat body-specific genetic manipulation in adult male and female *D. melanogaster*. Lrmda knockdown, knockdown of the SERCA homolog (the Drosophila ortholog of Atp2a1), and SERCA overexpression were each induced in the fat body of adult flies using the GeneSwitch-GAL4 (GS-GAL4) system, with transgene expression activated by feeding flies RU-486-containing food. Survival differences were assessed in both sexes by Kaplan-Meier, quantile regression (90th percentile), Cox proportional hazard, and time-varying hazard analyses. **(B)** Kaplan-Meier survival curves for Lrmda knockdown (Lrmda-kd) and control flies, with sexes pooled and sex included as a covariate. Median lifespan was compared using the log-rank test; maximum lifespan was compared using the quantile regression (90th percentile) test. Shaded bands indicate 95% confidence intervals. **(C)** Kaplan-Meier survival curves for Lrmda knockdown (Lrmda-kd) and control females. Median lifespan was compared using the log-rank test; maximum lifespan was compared using the quantile regression (90th percentile) test. BH-adjusted p-values are shown. Shaded bands indicate 95% confidence intervals. **(D)** Time-varying hazard ratio of Lrmda knockdown (Lrmda-kd) females relative to control flies. Hazards were estimated using a Cox proportional hazards model and a Cox model with a time-varying coefficient. Values above 1 indicate increased mortality risk, and values below 1 indicate decreased mortality risk relative to controls at the corresponding timepoint. HR, 95% CI, and BH-adjusted p-value reflect the overall Cox proportional hazards model estimates; the log-time interaction HR tests whether the hazard ratio changes significantly over time. **(E)** Kaplan-Meier survival curves for SERCA knockdown (SERCA-kd) and control flies, with sexes pooled and sex included as a covariate. Median lifespan was compared using the log-rank test; maximum lifespan was compared using the quantile regression (90th percentile) test. Shaded bands indicate 95% confidence intervals. **(F)** Time-varying hazard ratio of SERCA knockdown (SERCA-kd) males and females relative to control flies, with sexes pooled and sex included as a covariate. Hazards were estimated using a Cox proportional hazards model and a Cox model with a time-varying coefficient. Values above 1 indicate increased mortality risk, and values below 1 indicate decreased mortality risk relative to controls at the corresponding timepoint. HR, 95% CI, and p-value reflect the overall Cox proportional hazards model estimates; the log-time interaction HR tests whether the hazard ratio changes significantly over time. **(G)** Kaplan-Meier survival curves for SERCA knockdown (SERCA-kd) and control males. Median lifespan was compared using the log-rank test; maximum lifespan was compared using the quantile regression (90th percentile) test. BH-adjusted p-values are shown. Shaded bands indicate 95% confidence intervals. **(H)** Time-varying hazard ratio of SERCA knockdown (SERCA-kd) males relative to control flies. Hazards were estimated using a Cox proportional hazards model and a Cox model with a time-varying coefficient. Values above 1 indicate increased mortality risk, and values below 1 indicate decreased mortality risk relative to controls at the corresponding timepoint. HR, 95% CI, and BH-adjusted p-value reflect the overall Cox proportional hazards model estimates; the log-time interaction HR tests whether the hazard ratio changes significantly over time. **(I)** Kaplan-Meier survival curves for SERCA overexpression and control flies, with sexes pooled and sex included as a covariate. Median lifespan was compared using the log-rank test; maximum lifespan was compared using the quantile regression (90th percentile) test. Shaded bands indicate 95% confidence intervals. **(J)** Time-varying hazard ratio of SERCA overexpression males and females relative to control flies, with sexes pooled and sex included as a covariate. Hazards were estimated using a Cox proportional hazards model and a Cox model with a time-varying coefficient. Values above 1 indicate increased mortality risk, and values below 1 indicate decreased mortality risk relative to controls at the corresponding timepoint. HR, 95% CI, and p-value reflect the overall Cox proportional hazards model estimates; the log-time interaction HR tests whether the hazard ratio changes significantly over time. **(K)** Schematic of fat body-specific SERCA overexpression in adult male and female *D. melanogaster* using the GeneSwitch-GAL4 (GS-GAL4) system, with transgene expression activated by feeding flies RU-486-containing food. In females, SERCA was overexpressed continuously for the first 30 days of adult life, followed by no overexpression for the remainder of life. In males, a pulsed protocol was used: overexpression for 10 days, withdrawal for 30 days, re-induction for 10 days, followed by no overexpression for the remainder of life. Survival was assessed by Kaplan-Meier, quantile regression (90th percentile), Cox proportional hazard, and time-varying hazard analyses. **(L)** Kaplan-Meier survival curves for intermittent SERCA overexpression and control females. Median lifespan was compared using the log-rank test; maximum lifespan was compared using the quantile regression (90th percentile) test. BH-adjusted p-values are shown. Shaded bands indicate 95% confidence intervals, and the yellow shaded portion shows the period of SERCA overexpression. **(M)** Time-varying hazard ratio of intermittent SERCA overexpression females relative to control flies. Hazards were estimated using a Cox proportional hazards model and a Cox model with a time-varying coefficient. Values above 1 indicate increased mortality risk, and values below 1 indicate decreased mortality risk relative to controls at the corresponding timepoint. HR, 95% CI, and BH-adjusted p-value reflect the overall Cox proportional hazards model estimates; the log-time interaction HR tests whether the hazard ratio changes significantly over time. **(N)** Kaplan-Meier survival curves for intermittent SERCA overexpression and control males. Median lifespan was compared using the log-rank test; maximum lifespan was compared using the quantile regression (90th percentile) test. BH-adjusted p-values are shown. Shaded bands indicate 95% confidence intervals, and the yellow shaded portion shows the period of SERCA overexpression. Number of flies per experimental condition = 97-204; number of vials per condition = 5-9.

Consistent with the direction of change during rejuvenation, fat body-specific *Lrmda* knockdown extended lifespan. With sexes pooled and sex included as a covariate, *Lrmda*-knockdown flies showed increased median (log-rank p = 2 × 10⁻¹³) and maximum (p < 0.001) lifespan (**Figure 7B**), corresponding to a reduced and time-stable mortality risk (hazard ratio (HR) = 0.57, 95% CI 0.47–0.70, p = 2.38 × 10⁻⁸; **Supplementary Figure 8A**). The effect was female-specific: *Lrmda*-knockdown females lived markedly longer median (log-rank p.adjusted = 2 × 10⁻¹¹), and maximum (p.adjusted < 0.001; **Figure 7C**) lifespan with a consistently reduced HR (HR = 0.37, 95% CI 0.28–0.50, p.adjusted = 3.48 × 10⁻¹¹; **Figure 7D**), whereas males showed no change in median or maximum lifespan with a HR indistinguishable from controls (HR = 0.95, 95% CI 0.72–1.25; **Supplementary Figure 8B, 8C**).

Conversely, knockdown of *SERCA* strongly shortened lifespan in both sexes. With sexes pooled and sex included as a covariate, *SERCA*-knockdown reduced median (log-rank p < 2 × 10⁻¹⁶) and maximum (p < 0.001) lifespan (**Figure 7E**), with a HR that significantly worsened with age (HR = 3.77, 95% CI 3.10–4.58, p < 2 × 10⁻¹⁶, time-varying interaction p < 2 × 10⁻¹⁶; **Figure 7F**). The effect was most pronounced in males (log-rank p.adjusted = 4 × 10⁻¹⁶, maximum lifespan p.adjusted < 0.001; HR = 8.82, 95% CI 6.29–12.36, p.adjusted = 4 × 10⁻¹⁶, time-varying interaction p.adjusted = 0.002; **Figure 7G, 7H**) but also apparent, especially for the time-varying interaction, in females (log-rank p.adjusted = 2 × 10⁻¹⁴, maximum lifespan p.adjusted < 0.001; HR = 2.68, 95% CI 2.09–3.43, p.adjusted = 6 × 10⁻¹⁵, time-varying interaction p.adjusted = 7.66×10⁻¹⁵; **Supplementary Figure 8D, 8E**). The opposing directions of the *Lrmda* and *SERCA* knockdowns, protective and detrimental, respectively, mirrored the directions of their expression changes during WAT rejuvenation.

Given that *SERCA* was upregulated during rejuvenation, we next asked whether its overexpression could extend lifespan. Constitutive fat body-specific *SERCA* overexpression, however, conferred no median or maximum lifespan benefit with sexes pooled and sex included as a covariate (**Figure 7I**), and its HR, although not significantly different overall (HR = 1.12, 95% CI 0.94–1.34), rose steadily over time (time-varying interaction p = 0.005; **Figure 7J**). The same time-dependent pattern was observed in females (HR = 0.94, 95% CI 0.73–1.20, time-varying interaction p.adjusted = 0.04; **Supplementary Figure 8F, 8G**), while in males constitutive overexpression decreased median (log-rank p.adjusted = 0.06) and maximum lifespan (p.adjusted = 0.0012) and also raised the overall hazard with no significant interaction with time (HR = 1.39, 95% CI 1.07–1.81, p.adjusted = 0.028; **Supplementary Figure 8H, 8I**). Together, these results indicated that sustained SERCA overexpression confers no clear longevity benefit and becomes increasingly detrimental with continued induction.

Taking into account the exposure time dependency of the *SERCA* overexpression effects and its benefit at the beginning of the treatment, we reasoned that the transient overexpression of *SERCA* might still be beneficial. Therefore, we tested an intermittent overexpression protocol: continuous induction for the first 30 days of adult life in females (corresponding to reaching a HR of approximately 0.75 during constitutive overexpression), and two 10-day pulses (also corresponding to reaching a HR of approximately 0.75 during constitutive overexpression) separated by a 30-day withdrawal in males, each followed by no further induction (**Figure 7K**). This protocol extended lifespan in both sexes. Females showed increased median (log-rank p.adjusted = 4e-06) and maximum (p.adjusted = 0.0240) lifespan (**Figure 7L**) with a lower early-life HR that never reached the HR of controls due to timely withdrawal (HR = 0.57, 95% CI 0.45–0.72, p.adjusted = 5.4e-06, time-varying interaction p = 0.0078; **Figure 7M**). Males similarly showed extended median (log-rank p.adjusted = 0.01) and maximum (p.adjusted = 0.0156) lifespan (**Figure 7N**) with a reduced hazard that did not change significantly with time (HR = 0.74, 95% CI 0.6–0.93, p.adjusted = 8.2 × 10⁻³; **Supplementary Figure 8J**).

Together, these experiments identify *Lrmda* and *SERCA* as fat body-specific modulators of longevity whose effects are both sex- and exposure-time-dependent. *Lrmda* knockdown was beneficial only in females; *SERCA* was required for normal lifespan in both sexes, but extended life only when overexpressed transiently, not constitutively, with the effect being stronger in females.

## Discussion

We sought to test whether an aged tissue retains an intrinsic capacity for rejuvenation and, if so, which molecular programs mediate it. By transplanting subcutaneous WAT between young and old mice and tracking biological age and molecular changes longitudinally, we found that old adipose tissue placed in a young host undergoes rejuvenation across epigenetic and transcriptomic clocks of both chronological age and expected mortality, whereas young tissue placed in an old host rapidly ages. To our knowledge, this is the first study where the ability of the young environment to rejuvenate old tissues was assessed longitudinally, providing direct evidence for the existence of rejuvenation by showing that the biological age endpoint estimate becomes lower than its own baseline value.

Notably, rejuvenated grafts did not fully reach the age of young hosts. Elevated epigenetic age in isochronic settings suggests that the transplantation procedure induces measurable aging acceleration. Strong transcriptomic, proteomic, and metabolomic correlations across changes in heterochronic and isochronic experimental groups further suggest the presence of a shared pro-aging injury response program, likely encompassing implantation, vascularization, healing, and immune activation, confounding the interpretation of host-specific effects. Despite this shared graft-survival response, old adipose tissue transplanted into young hosts still underwent detectable rejuvenation. Targeting the deleterious nature of graft implantation, for example, with pharmacological dampening of post-transplant inflammation, may help amplify the rejuvenation observed in old-to-young contexts.

A central obstacle to interpreting bulk aging clock predictions is that they capture a signal from both cell-intrinsic changes and shifts in cell composition. This issue is especially acute during assessment of aging and rejuvenation following heterochronic transplantations. Single-nucleus profiling resolved this problem: the age reversal was intrinsic to mature, post-mitotic adipocytes, whereas the infiltrating immune cells had a minor or even opposite contribution to the age-reversal signal. Module-specific clocks further localized the composite adipocyte rejuvenation signal to the age reversal in chromatin modification, mRNA splicing, translation, and energy metabolism domains, characterizing a cell-type- and pathway-level anatomy of adipocyte rejuvenation. Interestingly, we did not observe accelerated aging of young adipocytes implanted into old compared with young recipients by either single-nucleus RNA sequencing or histological assessment, suggesting that the rapid aging signal detected by bulk aging clocks is largely attributable to infiltration of the graft by the aged immune cells of the recipient. If young cells can resist assimilation to the age of their environment, replacement of selected cell types by their young counterparts may represent a viable strategy for slowing systemic aging in old organisms.

ImAge, used to quantify the age of adipocyte chromatin, detected accelerated aging of young adipocytes transplanted into old hosts relative to the young-to-young control. This agrees with the transcriptomic data: the chromatin modification modules aged significantly in young-to-old adipocytes by both module clocks, but not in young-to-young adipocytes. Chromatin is one important layer of biological age among others, so our conclusions about which groups aged and which rejuvenated rest on composite clocks that integrate many layers.

Transcriptomic, proteomic, and metabolomic evidence converged on an energy balance rewiring as the core feature of the adipose tissue rejuvenation program. Therefore, we hypothesized that the energy metabolism rewiring should have changed a core phenotypic hallmark of adipocyte biology - the size of the lipid droplet. Lipid droplets are evolutionarily conserved organelles that serve as intracellular reservoirs of neutral lipids and play critical roles in maintaining energy, structural, and signaling homeostasis across diverse cell types ^56–59^. Mature adipocytes champion lipid storage function, typically harboring a single (unilocular), large lipid droplet that expands markedly in states of energy excess. In response to insulin, adipocytes mobilize lipids through lipolysis and release non-esterified fatty acids into the blood circulation. However, during aging or in the context of persistent caloric surplus, adipocytes may enter a hypertrophic state characterized by enlarged lipid droplets, loss of thermogenic function, and metabolic dysfunctions such as insulin resistance, obesity, and elevated risk of type 2 diabetes ^60–63^.

By contrast, brown and beige adipocytes are characterized by small, often fragmented (multilocular) lipid droplets dispersed throughout the cytoplasm ^64^. Individuals with higher levels of brown adipose tissue exhibit a markedly reduced prevalence of cardiometabolic diseases, including cardiovascular diseases, hypertension, and type 2 diabetes ^65^. This protective association is underpinned by the unique capacity of thermogenic adipocytes to increase energy expenditure by actively utilizing glucose and lipids - the process that improves systemic metabolic flexibility, reduces lipid burden, and collectively buffers against age- and obesity-associated metabolic decline.

The reduction of lipid droplet size in old adipocytes transplanted into young hosts is consistent with a shift toward the smaller, multilocular droplets that characterize thermogenic adipocytes. This morphological change was accompanied by the upregulation of established positive regulators of thermogenesis, including *Mb*, the validated beige markers *Tnfrsf9* and *Tmem26*, and the increased abundance of the β-adrenergic second messenger and a thermogenesis regulator, cAMP (**Table 1**). Of particular interest was the upregulation of *Atp2a1* (SERCA1), which encodes the skeletal-muscle calcium pump that has recently begun to appear in reports of UCP1-independent thermogenesis in subcutaneous WAT (**Table 1**). Our data extend this picture by placing *Atp2a1* (SERCA1) within a set of genes, including *Mybpc1*, *Mybpc2*, *Ttn*, *Obscn*, and others, whose expression changed in the same direction, with comparable magnitude, and concordantly at the transcript and protein levels. Given SERCA1’s critical role in regulating calcium homeostasis, we hypothesize that activation of SERCA1-mediated futile calcium cycling is the key step for initiation of the thermogenic program and rejuvenation in WAT.

Beyond thermogenesis, the rejuvenation program was associated with five further processes (**Table 1**). Adipocyte differentiation was strongly suppressed, which we interpret as a downstream consequence of thermogenic activation: as lipid droplets shrank, the demand for storage capacity would be expected to fall, and reduced differentiation is the anticipated response in both mature adipocytes and preadipocytes. Glutathione metabolism and detoxification pathways were suppressed together with the peroxisome pathway, which declined at both the transcript and protein levels. Because antioxidant defenses are induced in proportion to oxidant exposure, their coordinated decline may indicate a reduced oxidative burden, although we did not measure oxidative damage directly ^66^. Genes positively associated with adipose tissue disorders were downregulated, indicating an improved health status of the tissue, which we attribute to active thermogenesis. Structural support programs were upregulated, and glucuronic acid, a precursor of hyaluronic acid whose overproduction yields smaller adipocytes, increased in abundance; we interpret both as consequences of the reduced lipid droplet size. Finally, DNA repair factors were upregulated, which may have allowed the removal of accumulated DNA damage, potentially contributing to the reduction in biological age.

We note that the interpretations above should be read as testable hypotheses rather than conclusions. In summary, our data establishes the nature and direction of the molecular changes that accompany adipose tissue rejuvenation, together with a defined set of candidates for the perturbation experiments.

Aged adipose tissue also showed higher adipocyte size heterogeneity than young tissue before transplantation, and this heterogeneity decreased in old grafts placed into young hosts, suggesting that the dispersion of adipocyte size, and not only its mean, tracks adipocyte aging. Moreover, the dynamics of adipocyte population size heterogeneity changes perfectly mirrored the dynamics of the aging clock changes applied to the adipocytes. The alignment between these two parameters indicates that the heterogeneity feature is responsive to interventions aimed at reducing the biological age of adipose tissue, making it a novel, cheap, interpretable, and straightforward biomarker for assessing adipocyte biological age. The decrease in adipocyte population heterogeneity following the transplantation of old WAT into the young body could suggest either reversal of hypertrophy in large, dysfunctional adipocytes or selective elimination of the most hypertrophic cells, either outcome aligning with a rejuvenation phenotype. Overall, the parallels observed between lipid droplet sizes and molecular age of adipocytes suggest that obesity itself may represent a state of accelerated adipocyte aging. Under such a framework, targeting adipocyte aging, potentially through the induction of identified and/or known thermogenesis programs, could restore youthful adipocyte function and, consequently, alleviate obesity as a downstream outcome.

Rejuvenation was partial and selective. Mitochondrial morphology, an established aging feature, was not reset in old adipocytes, and although transplantation non-specifically restored collagen levels to youthful values, it also broadly increased fibrosis across all groups. The fact that some age-associated features reverse while others persist suggests that biological age reversal reflects the resetting of specific, likely function-relevant programs or an emergence of a new functional state rather than a complete return to the original youthful state ^67^, at least within the timeframe we examined.

The graft-host interaction was largely unidirectional, potentially due to size differences of interacting tissues. Molecular age shifts in the recipients’ native WAT, liver, and blood were small and inconsistent across clocks, tissues, and phenotypic measures. This asymmetry is critical - young adipose tissue was not sufficient to rejuvenate the body that received it, challenging the evolving replacement strategies ^68^. At the same time, the fact that aged cells retain the capacity to rejuvenate implies that reprogramming such cells in place may be a more tractable goal than replacing them.

Moving from rejuvenation to longevity, we tested two strongest conserved candidates in *Drosophila*, using adult-onset, fat body–restricted manipulation so that any effect would be causally related to the fly’s adipose tissue. Both genes behaved as the mouse data predicted. Knockdown of *Lrmda*, downregulated during rejuvenation, extended female lifespan, whereas knockdown of *SERCA*, upregulated during rejuvenation, shortened it, with a mortality risk worsening progressively with age. Importantly, for *SERCA*, the effect on lifespan was bidirectional, though only under the right schedule: sustained overexpression conferred no benefit and became harmful over time, whereas transient, intermittent induction extended lifespan in both sexes. The dependence of lifespan on fat body-specific *SERCA* levels matching the direction observed during WAT rejuvenation suggests a causal role of this pump in adipose tissue aging. The timing dependence demonstrates that the effect of interventions can vary based on when and for how long it happens, not in its direction alone - an important consideration when designing strategies aimed at lowering the biological age.

Collectively, our data establish that the biological age of white adipose tissue is partially reversible, that this reversal is intrinsic to mature adipocytes and organized around a non-canonical thermogenic program, with components of that program being conserved fat tissue-specific regulators of longevity. As aging and obesity increasingly converge to drive metabolic disease, the adipocyte-intrinsic circuitry defined here offers a foundation for interventions to restore energy balance, improve metabolic flexibility, slow systemic physiological decline, and extend life of an aging population.

### Limitations of the study

This study focused on male animals of the same genetic background. Opting for the same genetic background was crucial to mitigate potential challenges related to the immunocompatibility of transplanted fat. While these issues could have been addressed with the use of immunosuppressants, incorporating them in our study might have introduced a confounding variable. The decision to select only one sex was influenced by the unavailability of old female mice when the study was conducted. It would therefore be important to replicate the study in females and in genetically heterogeneous mice that more closely resemble the diversity found in human populations. Furthermore, the duration of exposure to the recipient environment is likely to critically influence the magnitude and trajectory of aging or rejuvenation, underscoring the importance of testing additional time points to better characterize the temporal dynamics of adipose tissue aging plasticity. We infer a reduced oxidative burden from the coordinated downregulation of glutathione metabolism, detoxification, and peroxisome pathways, but we did not quantify reactive oxygen species or oxidative damage. The metabolomic and proteomic analyses of divergence were performed on three pairs per group at a 10% false discovery threshold, so the rejuvenation candidates require further investigation and validation. Our *Drosophila* experiments carry constraints inherent to the model. The fly genome encodes a single SERCA, so these experiments cannot resolve whether the mouse phenotype depends specifically on Atp2a1 or on other SERCA paralogues. In addition, the fat body serves adipose, hepatic, and immune functions, and fat body–restricted manipulation therefore cannot separate these contributions. Finally, functional validation beyond fly models, using targeted perturbation strategies such as genetic knockout or RNA interference, will be essential to delineate the causal mechanistic contributions of candidate genes and pathways to adipose tissue aging, rejuvenation, and thermogenic potential.

## Resource availability

### Lead contact

Requests for further information should be directed to and will be fulfilled by the lead contact, Vadim N. Gladyshev

### Materials availability

No new plasmids, cell lines, or other unique reagents were generated in this study.

### Data and code availability

All the raw and processed data will be publicly available after publication.

## Acknowledgments

This work was supported by the National Institute on Aging (NIA) grants to V.N.G. The Project was supported by a Longevity Impetus Grant from Norn Group to A.M. This work was supported by NIH grants (R01DK099511 to L.J.G.) and the Joslin Diabetes Center DRC (P30 DK36836). A.M. and A. Tskhay are recipients of Doctoral Training Scholarships from the Fonds de Recherche du Québec–Santé (FRQS). The authors gratefully acknowledge the MicRoN (Microscopy Resources on the North Quad) Core for their support and assistance in this work.

We thank John Tower for the GeneSwitch fly stocks. We thank Norbert Perrimon for the CantonS fly stocks. We thank Dr. Roderick Bronson for the histopathological evaluation of white adipose tissue slides for the presence and degree of fibrosis. We are grateful to members of the Gladyshev and Van Raamsdonk laboratories for valuable discussions and insights throughout this work.

## Author contributions

A.M. and V.N.G. conceived and designed the study. A.M. and P.N. conducted mouse experiments with assistance from N.P.C. and M.F.H. Single-nucleus RNA sequencing was performed by Z.Z. S.Y. did fly experiments. S.J. and J.S.H. performed metabolomic profiling of tissues. J.A.P. performed proteomic profiling of tissues. J.L. and K.N. did ImAge profiling and analyses. Tissue processing was performed by A.M., C.G.d.M., A.W.E., J.R.P., W.M., B.Z., and S.I. C.G.d.M. carried out mitochondrial immunostaining, and C.G.d.M., A.W.E., P.V.A., and A.M. performed imaging of fixed tissues. Data analysis was conducted by A. Tyshkovskiy, A. Tskhay, A.M., K.Y., D.K., E.J., S.T., and A.E.T. Figures were made by A. Tyshkovskiy, A. Tskhay, A.M., C.G.d.M., and K.N. Data interpretation and manuscript drafting were performed by A.M., A. Tyshkovskiy, and V.N.G. V.N.G., J.M.V.R., L.J.G., J.C., W.Z., A.A.P., M.F.H., M.H., S.P.G., and A.V.T. supervised the project and provided materials and methods. V.N.G. secured funding. All authors reviewed, edited, and approved the manuscript.

## STAR★Methods

Key resources table

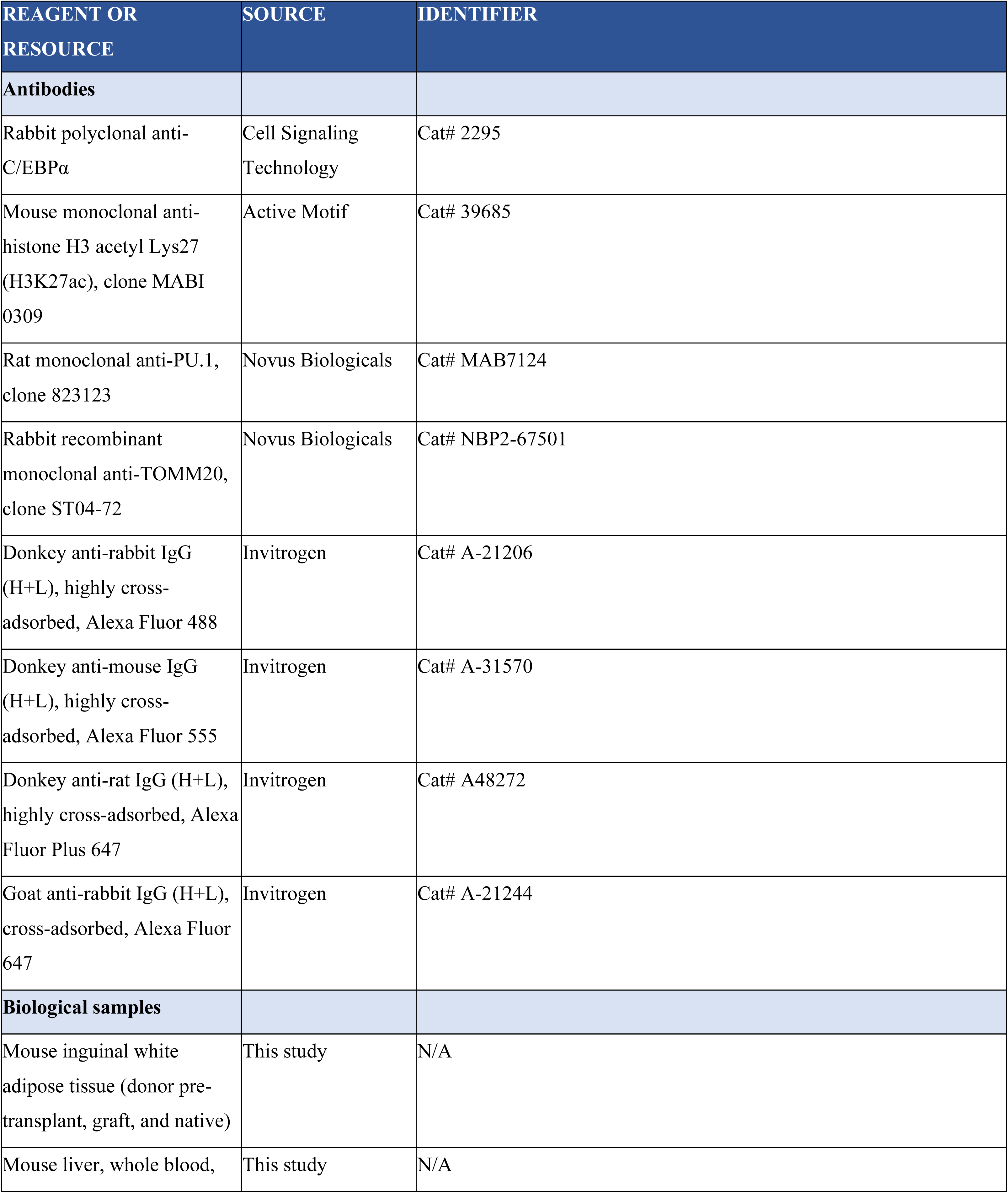
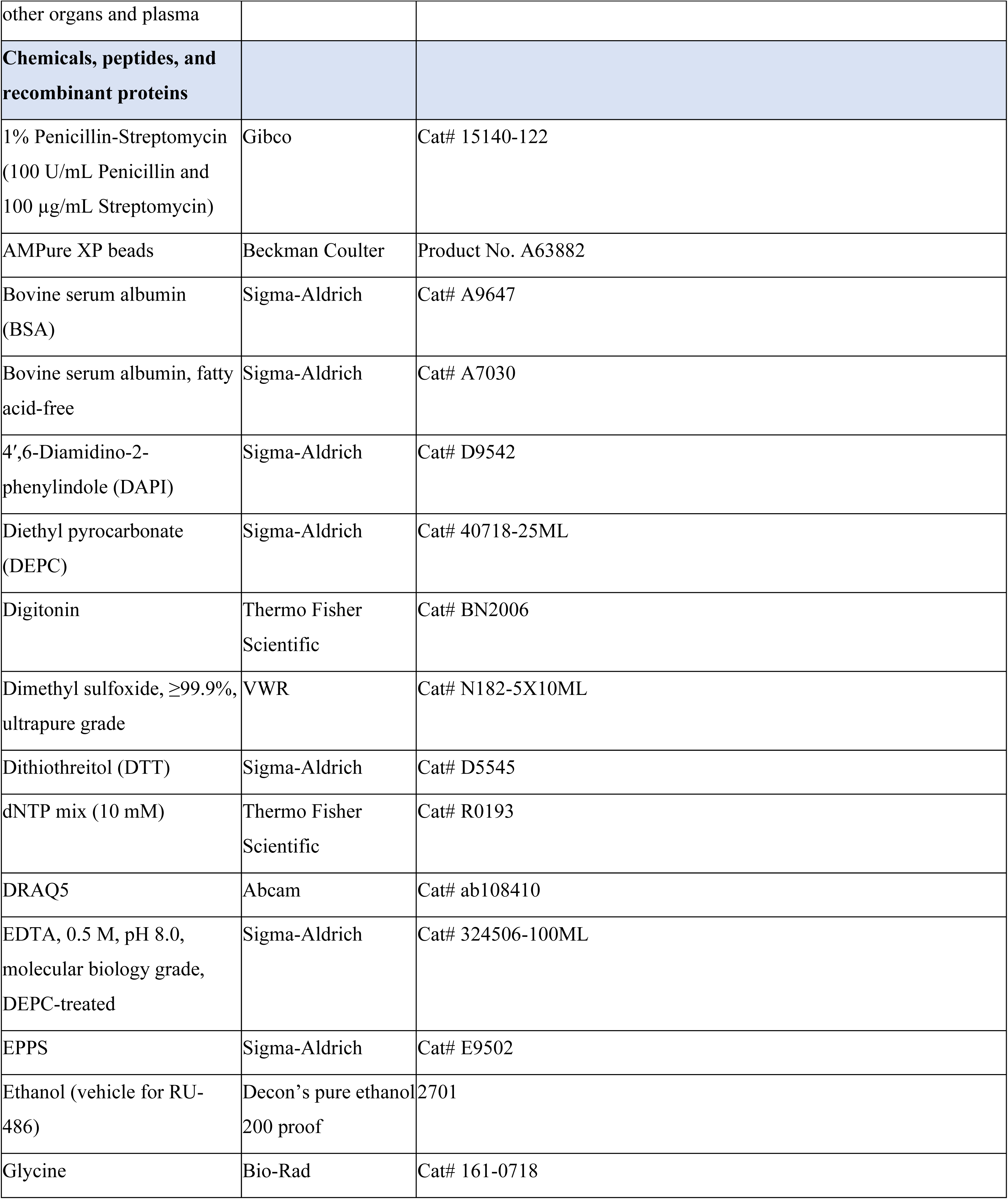
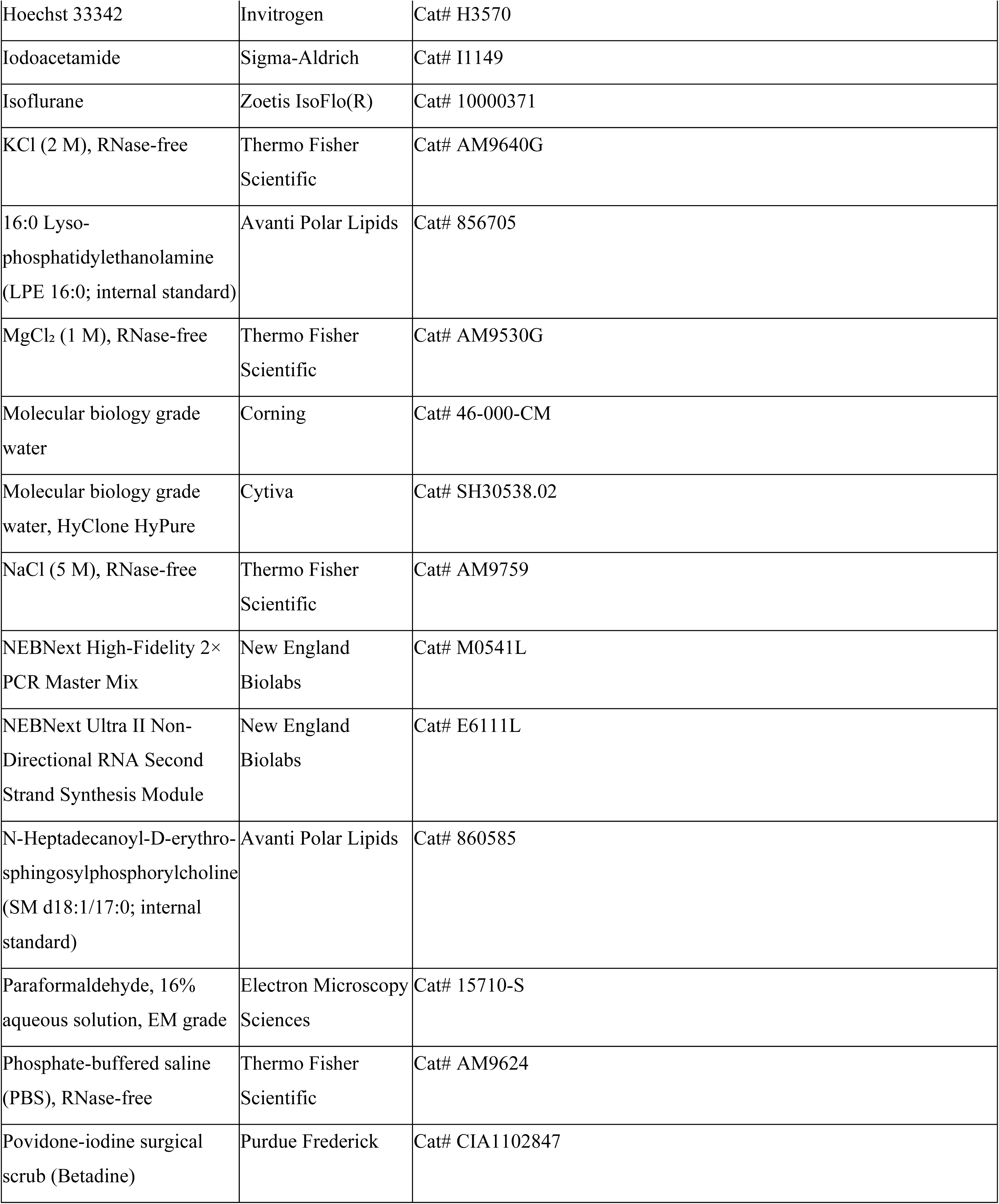
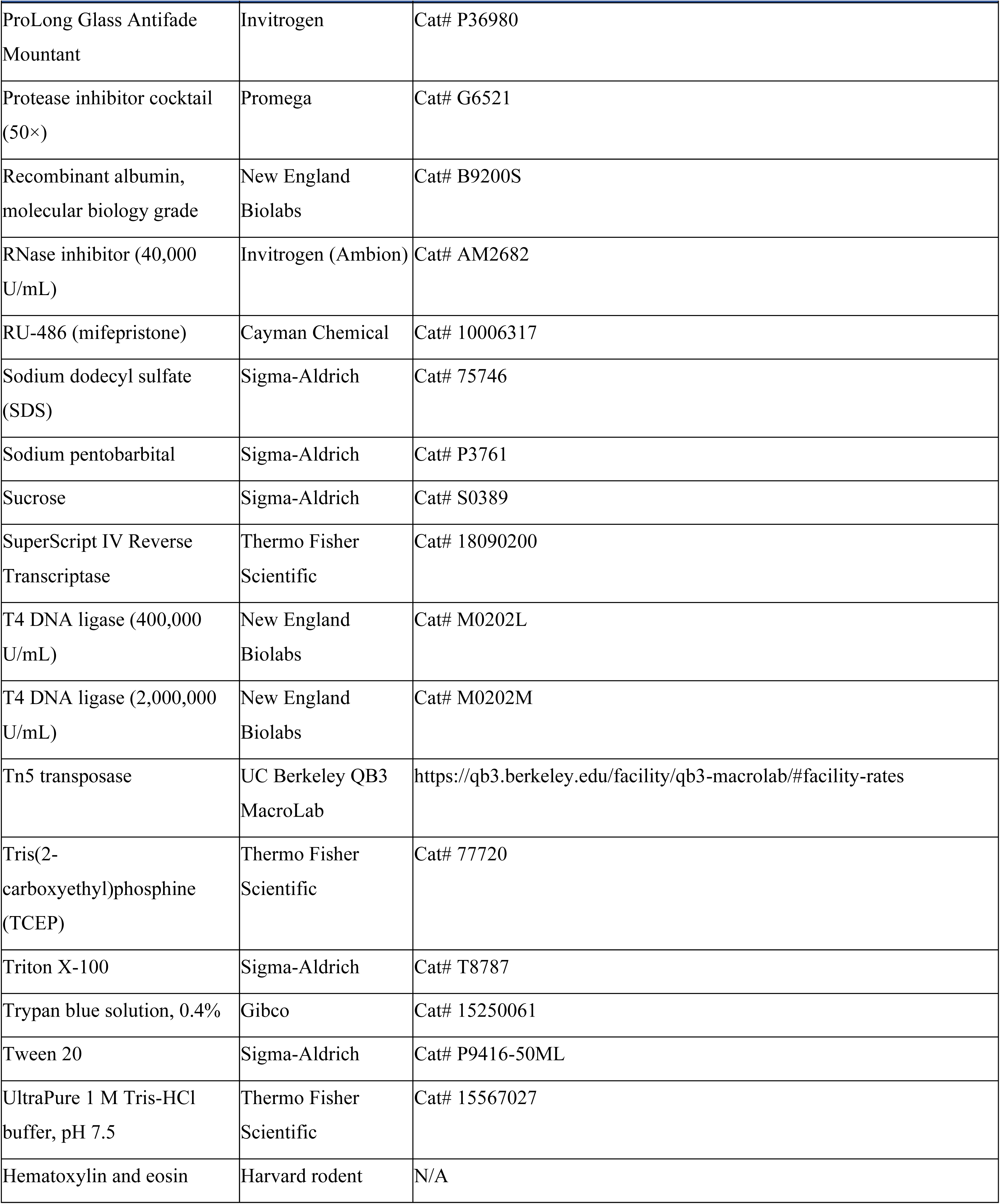
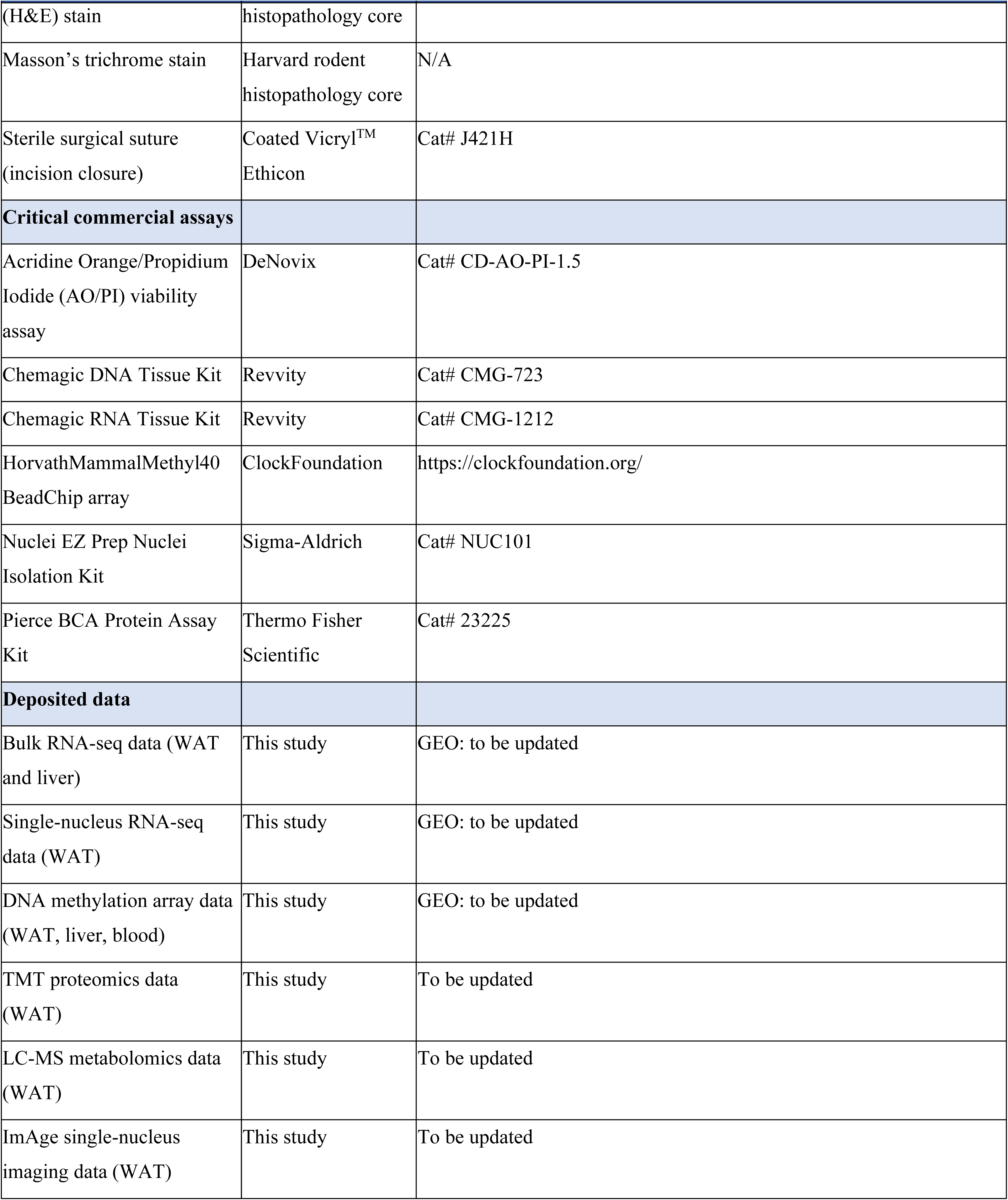
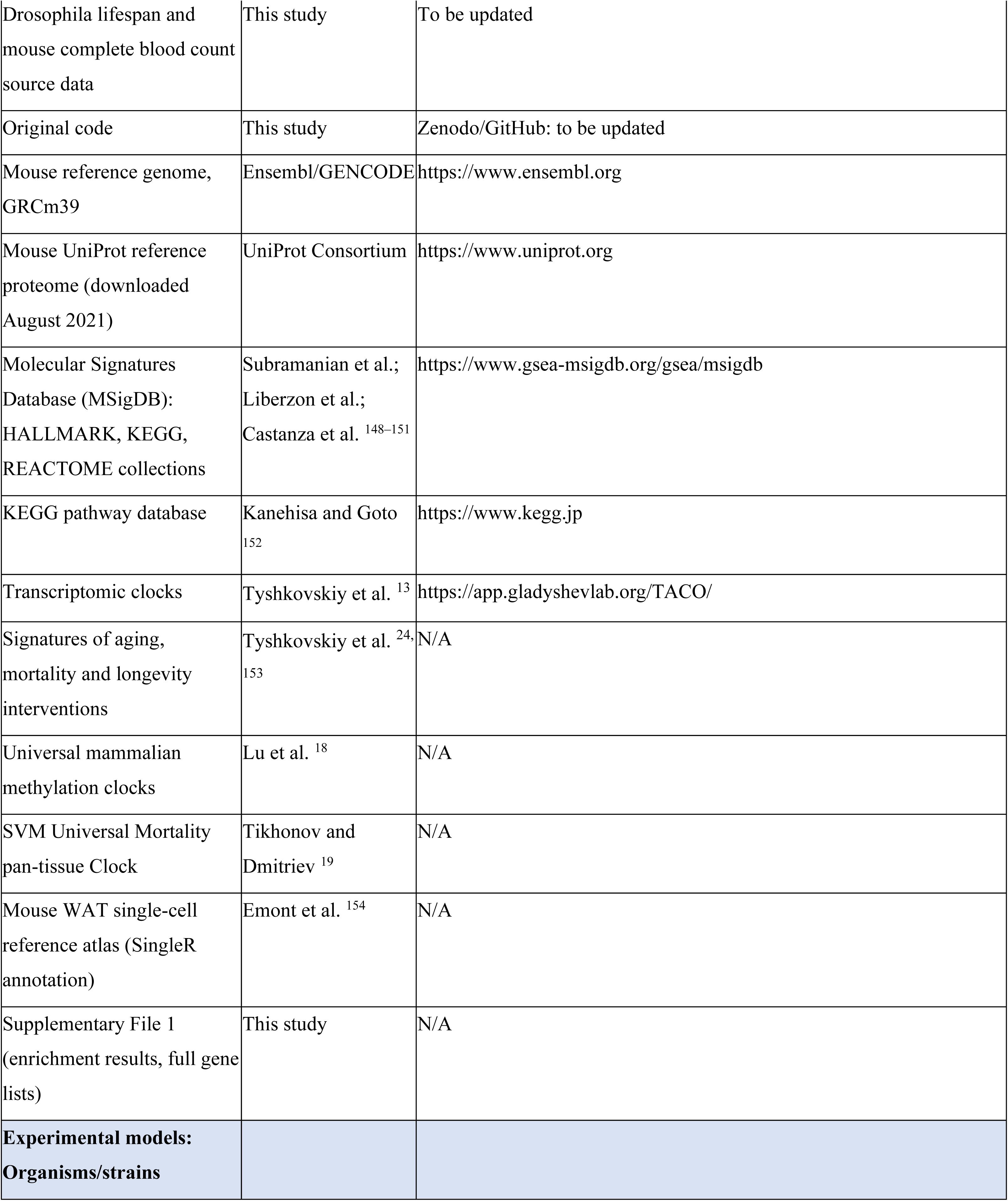
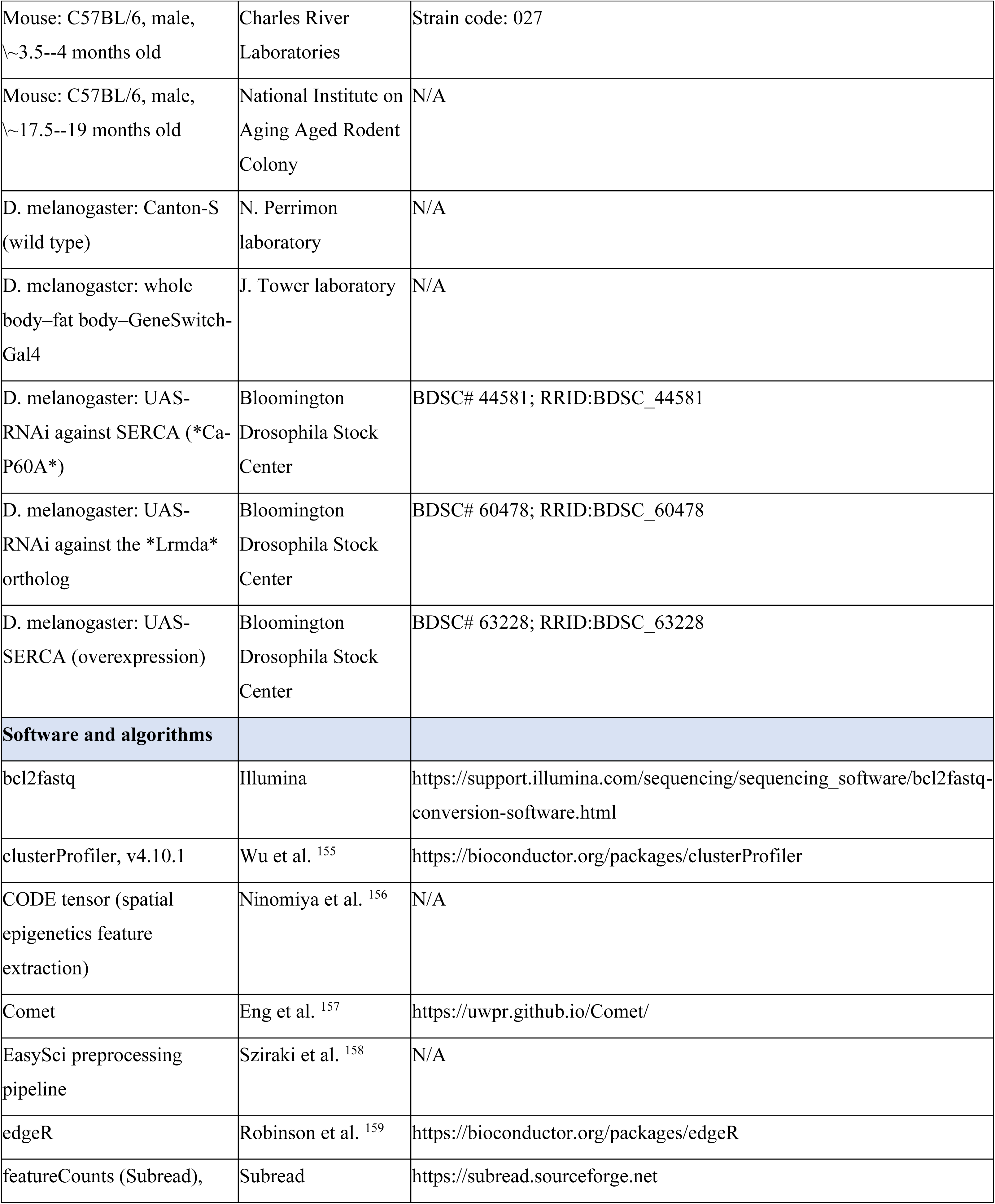
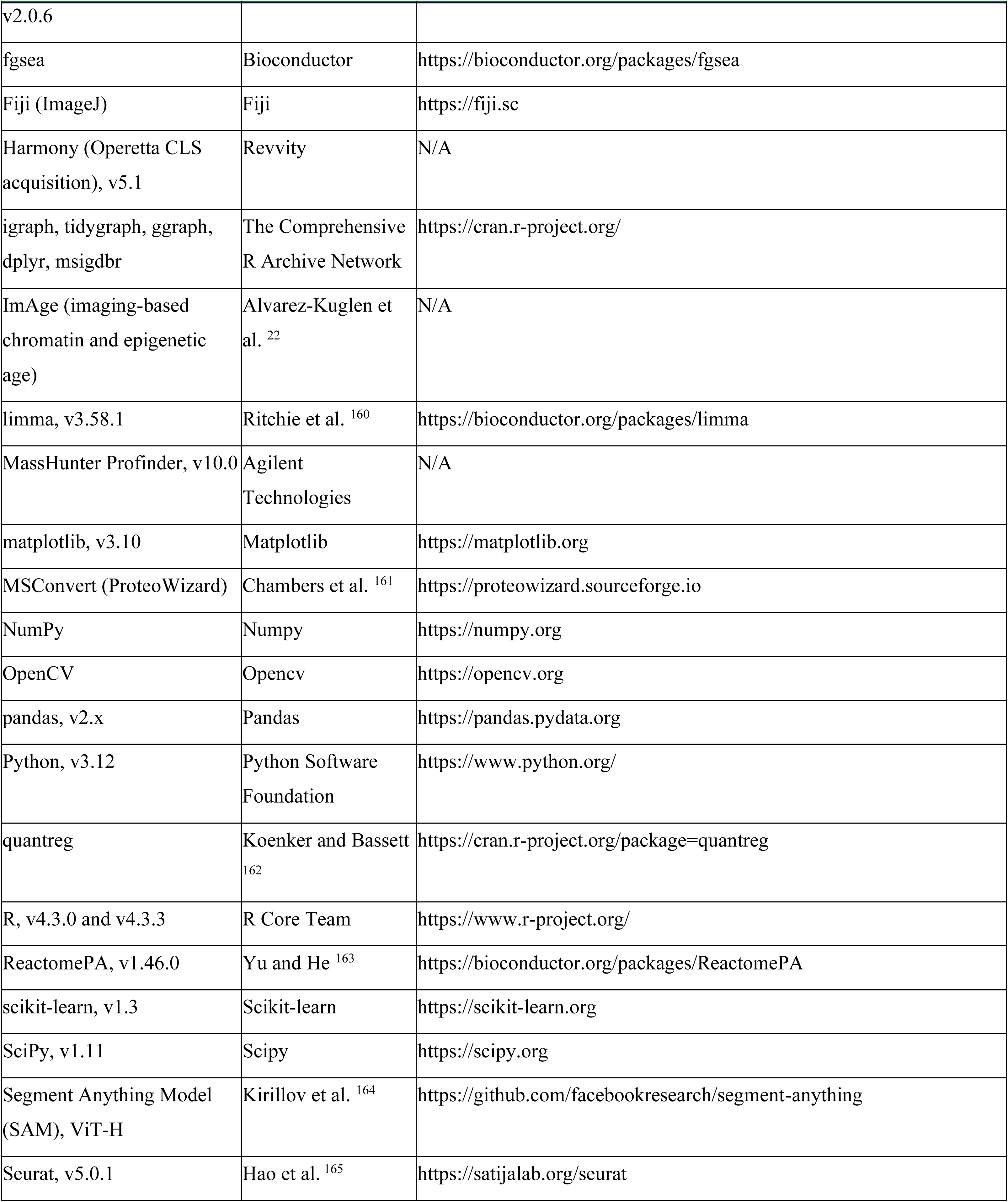
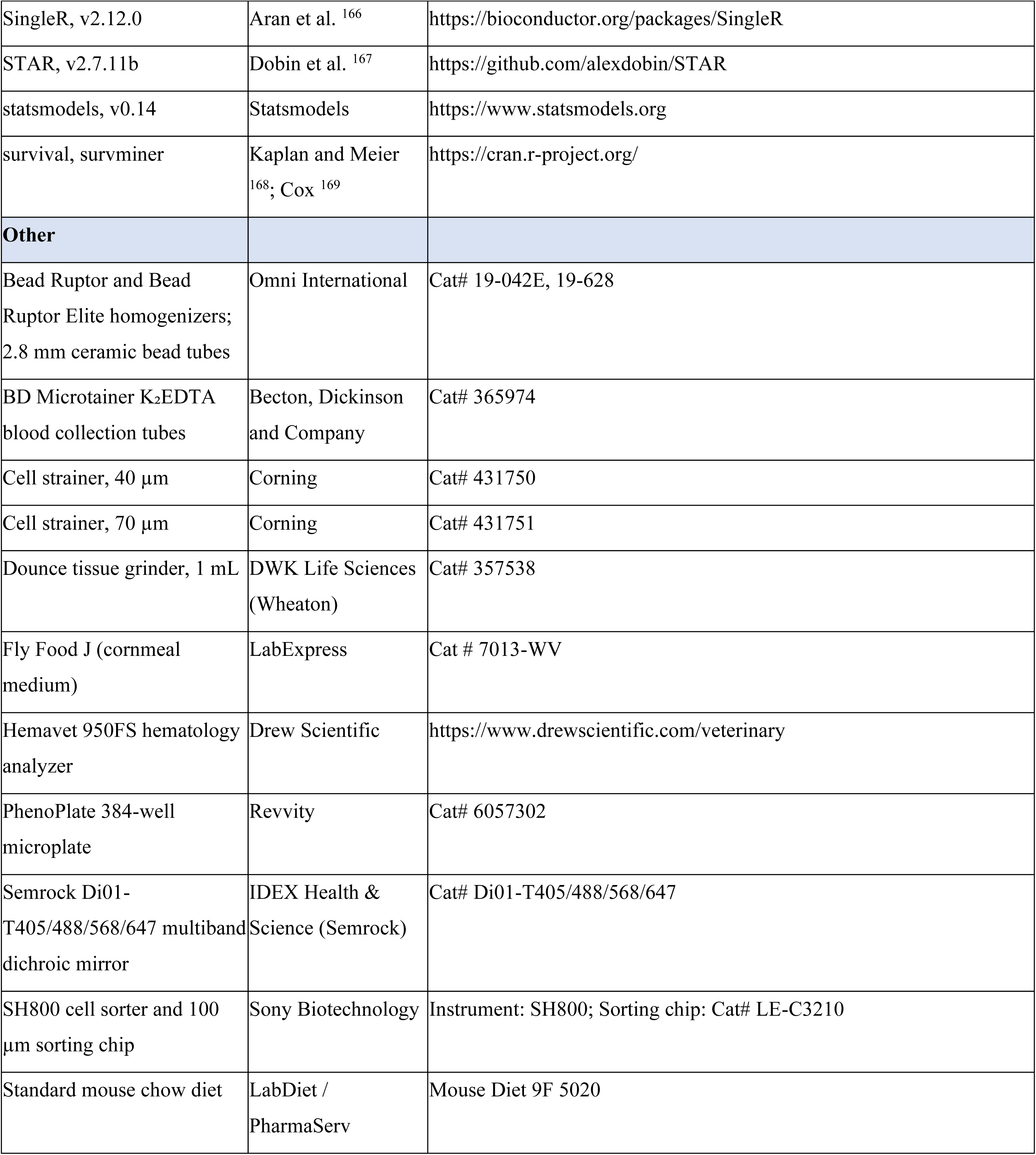

## Method details

### Animals

All experiments were conducted following NIH guidelines and protocols approved by the IACUC at Joslin Diabetes Center (JDC). All experiments were performed using ∼3.5- to ∼4.3-month-old male C57BL/6J mice purchased from Charles River Laboratories, MA, and ∼17.5 to ∼19.3-month-old male C57BL/6JN mice obtained from the National Institute of Aging aged colony. All mice were housed on a 12/12 h light/dark cycle. Standard mouse chow diet (9F 5020 Lab Diet, PharmaServ, Inc.) and water were provided ad libitum.

### Adipose tissue transplantation

All mice were randomly assigned to experimental groups (sham, donor, or recipient) one week prior to surgery. All animals underwent identical preparatory and surgical procedures, differing only in the manipulation of the inguinal white adipose tissue (iWAT): sham animals retained their native fat pads and received no transplants; donors underwent surgical excision of their iWAT pads, and recipients received donor fat implanted subcutaneously in the interscapular region, while their own iWAT remained intact. Surgeries were performed during a fixed 3-hour window to control for circadian variation, with group assignments randomized across surgical days. On the day of surgery, anesthesia was induced with intraperitoneal sodium pentobarbital (75 mg/kg for young and 65 mg/kg for old mice), followed by maintenance with 3.5% isoflurane administered via nose cone. Surgical sites were shaved and sterilized using Betadine. Core body temperature was maintained at 36°C throughout using a heating pad and monitored with a rectal probe. Inguinal fat pads were harvested from donors, weighed, and immediately transferred into 15 mL sterile tubes filled with 1% Penicillin-Streptomycin (100 U/mL Penicillin and 100 µg/mL Streptomycin; Gibco, Cat. No. 15140-122) dissolved in PBS and maintained at 37°C in a water bath. Donor incisions were closed using sterile sutures, and mice were allowed to recover on a 37°C heating pad. Recipient mice underwent a dorsal incision over the interscapular region, where donor iWAT was inserted subcutaneously. The amount of implanted tissue was normalized to 1% of the recipient’s body weight to ensure consistent graft load across animals. A portion of donor tissue was retained for histological and omics analyses, with samples for omics being flash-frozen in liquid nitrogen immediately after collection, and samples for histology fixed in 3.5% paraformaldehyde. Sham-operated animals were further randomized to receive surgical incisions at either the donor or the recipient sites. Following surgery, all mice were singly housed for the duration of the study. At study endpoint, the following tissues were collected for analysis after isoflurane-induced anesthesia followed by cardiac puncture to collect blood and plasma followed by removal of vital organs, such as: heart, liver, kidney, visceral white adipose tissue (vWAT), native subcutaneous white adipose tissue (except in donors), transplanted fat (in recipients), triceps, gastrocnemius muscle, cortex, skin at the surgical site, and skin from an unaffected area. Blood was collected into 500 µL K2EDTA-coated microtainer tubes (BD Microtainer with BD Microgard Closure, catalog no. 365974; Becton, Dickinson and Company). Complete blood counts were acquired within 5 minutes of collection on a Hemavet 950FS (Drew Scientific, Waterbury, CT). Blood was then centrifuged at 2,000 × g for 10 minutes at 4 °C to separate plasma.

### Isolation of nucleic acids

DNA and RNA were isolated using the Chemagic360 system (Revvity) with the appropriate kits (CMG-723 for tissue DNA; CMG-1212 for tissue RNA). Frozen adipose tissue samples were homogenized in the appropriate lysis buffer from Chemagic360 kits using a Bead Ruptor (Omni International) with 2.8mm ceramic bead filled tubes, using the 2 ml liver pre-set program. The homogenates were centrifuged (18,000 × g, 10 min) to remove insoluble material then processed according to the appropriate Chemagic360 protocol. DNA and RNA were eluted into molecular biology grade water (HyClone HyPure, Cytiva). When necessary, DNA samples were concentrated using a Centrivap (Labconco). Concentration of DNA/RNA samples was determined using Qubit or Quant-it kits (Invitrogen). Isolated DNA was stored at –20°C and isolated RNA was stored at –80°C until further processing.

### DNA methylation profiling

Methylation data were generated through the Epigenetic Clock Development Foundation using the HorvathMammalMethyl40 array (Illumina) ^16^ at AKESOgen Inc. Samples were randomized to avoid introduction of batch/chip effects. All sample preparation/processing was carried out according to the Illumina kit protocols. Epigenetic age (eAge) was estimated by applying Universal chronological clock and SVM-based multi-species multi-tissue mortality clock models to methylation β values. CpG sites with a detection p-value > 0.01 in at least 25% of samples were excluded. For the remaining CpGs, the mean methylation level and Shannon entropy were calculated individually for each sample. To compare eAge, mean methylation, and Shannon entropy among different WAT types (donor WAT Before transplantation, donor WAT After transplantation, and Native WAT), we used an ANOVA model with WAT type and donor ID as covariates within each experimental group (young-to-young, young-to-old, old-to-young, and old-to-old). Model coefficients and p-values for the WAT type factor were extracted, and p-values were adjusted for multiple comparisons using the Benjamini–Hochberg (BH) method. To evaluate differences in the dynamics of donor WAT eAge during transplantation across experimental groups, we applied an ANOVA model for each pair of experimental groups, including transplantation stage (“Before” and “After”), experimental group, and their interaction as covariates. The coefficient of the interaction term and its corresponding p-value were extracted, and p-values were adjusted for multiple testing using the BH method.

### RNA sequencing

RNA sequencing was performed at Novogene. Total RNA isolated as described above was checked for quality using an Agilent 2100 Bioanalyzer. Samples that passed QC were paired-end sequenced on an Illumina NovaSeq 6000 with 150 bp read length. Reads were mapped to a mouse genome (GRCm39) with STAR ^167^ (version 2.7.11b) and counted via featureCounts (version 2.0.6). To filter out non-expressed genes, we required at least 10 reads in at least 20% of samples. Differentially expressed genes between donor WAT before and after transplantation were identified separately for each profiled experimental group using ANOVA model with donor’s ID included as a covariate through edgeR package ^159^. Genes with differential change of expression after transplantation in donor WAT across experimental groups were identified for each pair of experimental groups using ANOVA model with stage of transplantation (“before” and “after”), experimental group, and their interaction included as covariates. Model coefficient of interaction term together with the corresponding p-value was used to characterize genes with distinct effect induced by transplantation across groups. P-values were adjusted for multiple testing with the BH method.

### Transcriptomic signature analysis

We conducted functional enrichment analysis to characterize the transcriptome changes in donor WAT induced by transplantation across different experimental groups, and compare them to established molecular signatures of aging, mortality, and lifespan regulation. We applied reference signatures from tissue-specific aging biomarkers of liver, kidney, and brain, and from multi-tissue biomarkers of chronological age and expected mortality adjusted for chronological age ^13,24^. Additionally, we included hepatic signatures of expected maximum and median lifespan in rodents as well as signatures of individual longevity interventions, such as caloric restriction, genetic models of growth hormone deficiency, and rapamycin ^153^. For each experimental transplantation group, we ranked gene expression changes during transplantation using a signed log-transformed p-value metric estimated through differential expression analysis:

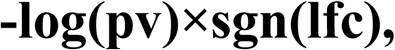

where pv and lfc are p-value and logFC of a certain gene, respectively, and sgn is the signum function (equal to 1, -1 and 0 if value is positive, negative or equal to 0, respectively). Ranked gene lists were then subjected to gene set enrichment analysis (GSEA) using the fgsea package in R, with 10,000 permutations and multilevel Monte Carlo sampling. Gene sets were drawn from the HALLMARK, KEGG, and REACTOME collections of the Molecular Signatures Database (MSigDB) ^148–152^. The same enrichment pipeline was applied to reference signatures of aging, mortality, and longevity interventions. Individual p-values were adjusted for multiple testing with the Benjamini-Hochberg (BH) method ^170^. We computed Spearman correlations between normalized enrichment scores (NES) to quantify similarities between signatures of transplantation and reference gene expression biomarkers.

### Transcriptomic age analysis

Filtered RNA-seq data were processed using Relative Log Expression (RLE) normalization, log transformation, and scaling. Missing expression values for clock genes absent from the dataset were imputed using their precomputed average values. Normalized gene expression profiles were then centered to the median profile of young samples derived from donor WAT before transplantation. Transcriptomic age (tAge) for each sample was estimated using Bayesian Ridge-based rodent multi-tissue transcriptomic clocks of chronological age and expected mortality ^13^. Pairwise differences in tAge across WAT groups were evaluated separately for each experimental group using a mixed-effects model, with both the mean and standard deviation of tAge predictions included, and donor ID and batch included as technical covariates. To assess differences in donor WAT tAge dynamics following transplantation across experimental groups, tAges were first adjusted for chronological age. Specifically, a model mapping chronological age to tAge was fitted using sham control samples and then applied to other samples to generate expected tAge values. The difference between observed and expected tAge was calculated for each sample, and for each pair of experimental groups, a mixed-effects model was fitted with transplantation stage (Before vs. After), experimental group, and their interaction as fixed effects, and batch as a technical covariate. P-values for the interaction terms were adjusted for multiple testing using the BH method, and these adjusted values were used to determine statistical significance of differences in tAge dynamics across groups.

To estimate the contribution of each gene to tAge changes during transplantation, we multiplied its log fold change (logFC) by the corresponding coefficient from Elastic Net-based transcriptomic clock, calculated separately for each experimental group. The statistical significance of each gene’s contribution was derived from the corresponding differential expression analysis. Comparisons of gene contributions to tAge dynamics across experimental groups were performed using Pearson correlation analysis.

To identify genes whose expression changes during transplantation were significantly associated with tAge dynamics, we analyzed samples from donor WAT collected before and after transplantation. A linear model was fitted using the edgeR package, with tAge difference, transplantation stage, and donor ID included as covariates. Model coefficients and p-values for the tAge difference term were extracted, and p-values were adjusted for multiple testing using the BH method. Genes with an adjusted p-value < 0.05 were considered significant signatures of tAge dynamics deviation during donor WAT transplantation. Pathway enrichment analysis for these genes was performed separately for those positively and negatively associated with tAge dynamics using Fisher’s exact test in the clusterProfiler package ^155^, based on KEGG ontology. Resulting p-values were adjusted using the BH method, and KEGG terms with an adjusted p-value < 0.05 were considered statistically significant pathway signatures of tAge-associated genes.

Module-specific transcriptomic clocks of chronological age and expected mortality were applied to the scaled relative gene expression profiles using the same framework. The resulting tAge estimates from each module-specific clock were standardized within each experimental group. For each module-specific clock and experimental group, an ANOVA model with donor ID as a technical covariate was used to compare tAge between WAT samples collected before and after transplantation. Resulting p-values were adjusted for multiple comparisons using the BH method.

### Adipose tissue nuclei extraction for single-nucleus RNA-sequencing

The processing of samples was conducted in accordance with the established PanSci methodology with minor adjustments ^171^. Briefly, a 10X PBS-hypotonic stock solution was prepared as previously described. Fresh 1X hypotonic lysis buffer was prepared on the day of nuclei extraction by diluting the stock solution with RNase-free water, supplemented with 0.33 M sucrose and 3 mM MgCl2. For each adipose tissue sample, a working lysis buffer was freshly prepared by adding 1% diethyl pyrocarbonate and 0.01% digitonin to 10 mL of the 1X hypotonic lysis buffer. Dry tissue powder, maintained in liquid nitrogen, was rapidly transferred into the prepared working solution. Following a 10-second vortex to disperse tissue aggregates, the mixture was incubated at 4°C for 15 minutes with constant rotation. The suspension was filtered through a 40 μm cell strainer, and the filter was rinsed with an additional 5 mL of lysis solution. Extracted nuclei were recovered via centrifugation at 500 × g for 5 minutes at 4°C and resuspended in a nuclei suspension buffer (10 mM Tris-HCl pH 7.5, 10 mM NaCl, 3 mM MgCl2, and 0.2 mg/mL recombinant albumin). For fluorescence-activated cell sorting (FACS), the suspension was supplemented with DRAQ5. FACS was performed using a SH800 Cell Sorter with a 100 μm sorting chip, specifically gating for DRAQ5-positive singlet nuclei to ensure global cell population recovery and the removal of debris and doublets. Sorted nuclei were collected into a 1.5 mL tube containing 100 μL of nuclei suspension buffer and subsequently concentrated by centrifugation at 500g for 5 minutes at 4°C.

### EasySci library construction and sequencing data preprocessing

Sequencing libraries were generated according to the previously described EasySci RNA protocol with specific parameters ^158^. Briefly, sorted nuclei were distributed into 96-well plates for reverse transcription (RT). The first round of indexing was achieved by utilizing indexed oligo-dT and indexed random hexamer primers. Nuclei were pooled, washed, and redistributed into subsequent 96-well plates for the introduction of a second index via ligation. Following an additional pooling and washing cycle, nuclei were diluted and distributed for second-strand synthesis, followed by purification with AMPure XP beads. Eluted cDNA was subjected to tagmentation using Tn5 transposase and amplified via a 15-cycle PCR reaction with indexed P7 primers. The final PCR products were pooled and underwent two rounds of purification using a 0.8X volume of AMPure XP beads. Library concentration and fragment size distribution were assessed using an Agilent TapeStation, and sequencing was performed on an Illumina NovaSeq X System. Base call (BCL) files were demultiplexed based on the final PCR indexing round using Illumina bcl2fastq conversion software to generate FASTQ files. For single-nucleus RNA-seq analysis, read alignment and the generation of gene/exon count matrices were performed using the established EasySci pipeline ^158^. Nuclei were filtered from the count matrix based on the following criteria: an unmatched rate below 0.4, a combined oligo-dT and random hexamer UMI (Unique Molecular Identifier) count of at least 200, and a minimum of 100 detected genes.

### Single-nucleus RNA sequencing analysis

snRNA-seq data were processed with Cell Ranger (v7.0.0) and Seurat (v5.0.1) package. Cells were annotated with SingleR (v2.12.0) ^166^ using a mouse WAT single-cell atlas as the reference ^154^. snRNA-seq data were normalized, subjected to PCA, and visualized with UMAP in Seurat ^165^. Within each annotated cell type, nuclei were aggregated into metacells to achieve a target coverage of ≥500,000 reads per metacell. Genes with at least 5 reads in at least 25% of metacells within each cell type were retained for downstream analysis. Metacell data were then processed using the bulk RNA-seq pipeline as described in the ‘Transcriptomic age analysis’. Normalized gene expression profiles were centered relative to the median profile across all samples. tAge for each metacell was estimated using Bayesian Ridge-based mouse multi-tissue transcriptomic clocks of chronological age and expected mortality ^13^. Statistical analyses were performed separately for mature adipocytes, B cells and T cells, and monocytes and macrophages, as described for the bulk RNA-seq data.

### Isolation of nuclei from white adipose tissue for ImAge

Nuclei were isolated from mouse inguinal white adipose tissue (iWAT) samples using the Nuclei EZ Prep Nuclei Isolation Kit (Sigma-Aldrich, NUC101) with minor modifications from previously published protocol ^156^. Approximately 100 mg of frozen scWAT was thawed on ice, finely minced, and homogenized in 500 μL of ice-cold EZ Nuclei Lysis Buffer supplemented with Protease Inhibitor and RNase inhibitor using a 1-mL Dounce homogenizer (10 loose and 10 tight strokes). The homogenate was incubated on ice for 5 min to complete cell lysis. The lysate was filtered through a 70-μm cell strainer into a 15-mL conical tube, and the strainer was rinsed with 1 mL of additional ice-cold EZ Nuclei Lysis Buffer to maximize nuclei recovery. An additional 1 mL of EZ Nuclei Lysis Buffer was added to the filtrate before centrifugation at 600 × g for 4 min at 4°C. Following centrifugation, the floating lipid layer was carefully removed while preserving the nuclei-containing suspension. The suspension was then passed through a 40-μm cell strainer, which was rinsed with an additional 1 mL of EZ Nuclei Lysis Buffer. The filtered nuclei suspension was centrifuged at 500 × g for 10 min at 4°C, and the supernatant was discarded. The nuclei pellet was resuspended in 1 mL of Nuclei Wash and Resuspension Buffer (PBS supplemented with 0.5% fatty acid-free bovine serum albumin (BSA)) and washed by centrifugation at 500 × g for 5 min at 4°C. The final nuclei pellet was resuspended in Nuclei Wash and Resuspension Buffer and quantified using the Acridine Orange/Propidium Iodide (AO/PI) Viability Assay (DeNovix) 1:1 volume prior to seeding nuclei into 384-well plate.

Nuclei were plated in a 384-well plate by seeding 7,000 nuclei/well in a 20 µL volume. Nuclei were centrifuged in a 384-well plate at 1,000 x g (acceleration 9, deceleration 4) for 10 min at 4 °C. Nuclei were then fixed with 4% paraformaldehyde (PFA) in PBS for 10 min at room temperature. The PFA solution was removed, and 30 µL of 0.1 M glycine in PBS was added for 5 min at room temperature. The solution was removed and samples were washed with PBS 2x and 50 µL of PBS was added for storage at 4 °C until immunofluorescence labeling. Nuclei were blocked with 2% BSA in PBS 0.5% Triton X-100 for 1 hr at room temperature and then incubated with primary antibody overnight at 4 °C and washed with PBS 3x (5 min each at room temperature). The following primary antibodies were used to label at least two wells per experimental animal. Mouse monoclonal anti-H3K27ac antibody (clone MABI 0309; Active Motif, Cat# 39685, RRID:AB_2793305) at a dilution of 1:250, Rabbit polyclonal anti-C/EBPα antibody (Cell Signaling Technology, Cat# 2295, RRID:AB_10692506) at a dilution of 1:250. and Rat monoclonal anti-PU.1 (clone 823123; Novus Biologicals, Cat# MAB7124; RRID: AB_3658704) at a dilution of 1:1,000.

Next, nuclei were incubated with secondary antibody for 1 hour at room temperature and then washed with PBS 3x (5 min each at room temperature). Primary antibodies were detected using the following secondary antibodies for their appropriate hosts. Donkey anti-rabbit IgG (H+L), Alexa Fluor™ 488 highly cross-adsorbed secondary antibody (Invitrogen, Cat# A21206, RRID:AB_2535792) used at 1:500. Donkey anti-mouse IgG (H+L), Alexa Fluor™ 555 highly cross-adsorbed secondary antibody (Invitrogen, Cat# A-31570, RRID:AB_2536180) used at 1:250. Donkey anti-rat IgG (H+L), Alexa Fluor™ Plus 647 highly cross-adsorbed secondary antibody (Invitrogen, Cat# A48272, RRID:AB_2893138) used at 1:500 for immunofluorescence. All wells were washed with 100 µL PBS 3x (5 min each at room temperature) and stain with Hoechst 33342 in 30 µL of 0.01 mg/mL concentration. Wash with 100 µL of PBS and add 50 µL of PBS per well before imaging.

### Confocal imaging of isolated white adipose nuclei for ImAge

Isolated nuclei were imaged in 384 wells plate using an Operetta CLS High-Content Analysis System (Revvity; formerly PerkinElmer) operated in confocal spinning-disk mode and controlled using Harmony software (v5.1). Images were acquired using a 63x water-immersion objective (NA 1.15). Hoechst 33342 fluorescence was acquired using the 370-nm excitation channel with a 430–500 nm emission filter. Alexa Fluor™ 488 fluorescence was acquired using the 475-nm excitation channel with a 500–550 nm emission filter. Alexa Fluor™ 555 fluorescence was acquired using the 550-nm excitation channel with a 570–650 nm emission filter. Alexa Fluor™ 647 fluorescence was acquired using the 630-nm excitation channel with a 655–760 nm emission filter. Autofocus was performed using the Harmony default Two-Peak autofocus mode for microplates.

For each well, at least 27 non-overlapping fields of view were acquired. At least two wells were imaged for each mouse sample, corresponding to a minimum of 54 fields per mouse sample. At each field, a z-stack comprising 20 optical sections was acquired with a 0.8-µm z-step, corresponding to a total imaging depth of 15.2 µm. Tiff image files were exported from Harmony software (v5.1), and a minimum of 300 nuclei per experimental condition were included in downstream analyses.

### Imaging-based chromatin and epigenetic age (ImAge)

To quantify biological age at the level of chromatin organization within individual nuclei, we applied ImAge, an imaging-based measure of chromatin and epigenetic age ^17^. WAT samples were imaged at single-nucleus resolution for DNA (Hoechst 33342) and active chromatin (H3K27ac, histone H3 lysine 27 acetylation), together with C/EBPα (CCAAT/enhancer-binding protein α) and PU.1 (SPI1). Analysis was restricted to mature adipocytes, defined as nuclei positive for C/EBPα, a marker of mature adipocytes ^172,173^, and negative for PU.1, a transcription factor restricted to hematopoietic lineages ^174^. For samples in which fewer than 25 nuclei passed this gate, the 25 nuclei with the highest C/EBPα signal were retained.

Each nucleus was represented by a set of image features computed with the Coordination of Chromatin Determinants (CODE) tensor, a machine-learning framework for spatial epigenetics that quantifies the co-localization and three-dimensional spatial organization of multiple chromatin determinants within single nuclei in situ [Ninomiya et al., companion manuscript]. Here, the CODE tensor was applied to the two imaged marks, Hoechst 33342 and H3K27ac; full feature definitions and implementation are provided in the companion manuscript.

The ImAge aging axis was defined as a linear support vector machine (SVM) classifier trained to separate young from old reference nuclei, using CODE tensor features as input, with the signed distance of each nucleus from the classifier decision boundary serving as a continuous single-cell aging score ^17^. A single reference axis was used, trained on pre-transplant young versus old donor tissue. Because grafts originate from donor tissue, this donor axis provides a common reference for both pre-transplant samples and grafts, and the use of a single axis keeps the model simple and interpretable.

All scores were obtained by leave-one-mouse-out cross-validation, with observations grouped by animal. For each held-out mouse, the SVM and all feature normalization statistics were fit exclusively on nuclei from the remaining mice, and nuclei of the held-out mouse were scored by projection only, so that no mouse contributed to its own prediction. Grafted samples were excluded from all training and were scored solely by projection onto the reference axis. Per-mouse aging scores were summarized as the median of the single-cell scores of that mouse within each condition. The axis was affine-calibrated to chronological age using the young and old donor references, and the resulting scores were rescaled to a relative range of 0 to 1 across pre-transplant and grafted samples (0, youngest sample; 1, oldest sample). The reported readout is therefore the median relative ImAge per sample and condition, and values are comparable within, but not beyond, this dataset.

Changes in ImAge between WAT Before and WAT After were assessed within each transplantation condition at three levels of comparison. At the pairwise sample level, a paired t-test was applied across transplants, pairing each graft with the pre-transplant value of its own donor. At the population sample level, a Mann–Whitney U test compared graft sample medians with the pooled sample medians of all donors of the corresponding age, comparing young-graft conditions with the young donor pool and old-graft conditions with the old donor pool. At the single-cell level, a Mann–Whitney U test compared individual graft nuclei with the nuclei of the matched pre-transplant donor samples. Tests were one-sided for the conditions in which the direction of change was specified in advance by the experimental hypothesis (old-to-young, rejuvenation; young-to-old, aging), and two-sided for the isochronic conditions (young-to-young and old-to-old), for which no directional hypothesis was defined.

Differences in the magnitude of change between conditions were assessed on the change in ImAge (Δ, median ImAge After minus median ImAge Before) at two levels. At the sample level, a two-sided Mann–Whitney U test was applied to the per-transplant Δ. At the single-cell level, a two-sided Mann–Whitney U test was applied to the per-nucleus Δ, in which each graft nucleus was expressed relative to the median of the pre-transplant nuclei of its own donor. P-values were adjusted for multiple comparisons using the BH method within each test type, across the four transplantation conditions for the within-condition tests and across the two contrasts for the between-condition tests. All reported values are leave-one-mouse-out cross-validated estimates.

### Sample processing for quantitative TMT-proteomic analysis

Adipose tissue was prepared for tandem mass tag (TMT)-based proteomics by following a previously published procedure with minor modifications ^175,176^. ∼20 mg of flash-frozen adipose tissue was placed in 2 ml tubes pre-filled with 2.8 mm ceramic beads and homogenized in modified RIPA buffer (50 mM KPO4 (pH 7.5), 150 mM NaCl, 1% Triton X-100, 0.5% deoxycholate, 2% SDS, and 1X protease inhibitors) on a Bead Ruptor Elite homogenizer (OMNI International, Kennesaw, GA). The samples were then spun at 20,000 × g for 10 minutes (4°C) and the supernatants were transferred to fresh microfuge tubes. Protein concentration was measured by BCA assay (Thermo-Fisher, Waltham, MA), and then 100 µg of protein from each sample were reduced with 5 mM TCEP for 30 minutes, alkylated with 10 mM iodoacetamide for 30 minutes, and quenched with 10 mM DTT for 15 minutes at room temperature. After this, the protein samples were chloroform-methanol precipitated, dried under speed-vac, and resuspended in 100 µl of 200 mM EPPS, pH 8.5. Protein samples were then digested with 1:100 Lys-C:protein overnight at room temperature, followed by 1:100 trypsin:protein for 6 hours at 37°C. The samples were then labeled with tandem mass tag (TMTpro) reagents. Acetonitrile was added to a final volume of 30% prior to adding the TMTpro labeling reagent. For protein level analysis, ∼100 µg of peptides were labeled with 100 µg of TMT. Labeling occurred at room temperature for 1h. Once labeling efficiency was verified (here, >97%), hydroxylamine was added at a final concentration of ∼0.3% and incubated for 15 min. at room temperature. TMTpro-labeled samples were pooled at a 1:1 ratio across all samples. The pooled sample was vacuum centrifuged to near dryness and subjected to C18 solid-phase extraction (SPE) (Sep-Pak, Waters).

#### Off-line basic pH reversed-phase (BPRP) fractionation (whole proteome)

We fractionated the pooled TMT-labeled peptide via BPRP HPLC with an Agilent 1260 pump. Peptides were subjected to a 50 min linear gradient from 5% to 35% acetonitrile in 10 mM ammonium bicarbonate pH 8 at a flow rate of 0.25 mL/min over an Agilent 300Extend C18 column (3.5 μm particles, 2.1 mm ID and 25 cm long). The peptide mixture was fractionated into a total of 96 fractions, which were consolidated into 24 super-fractions. These fractions were subsequently acidified with 1% formic acid and vacuum centrifuged to near dryness. Each fraction was also desalted via StageTip, dried via vacuum centrifugation, and reconstituted in 5% acetonitrile, 5% formic acid for LC-MS/MS processing.

### Liquid chromatography and mass spectrometry data acquisition for whole proteome analysis

Mass spectrometry data for all 24 super-fractions were collected using a Orbitrap Exploris480 mass spectrometer (Thermo Fisher Scientific, San Jose, CA) coupled nLC-1200 liquid chromatograph. Peptides were separated on a 100 μm inner diameter microcapillary column packed with ∼35cm of Accucore C18 resin (2.6 μm, 150 Å, Thermo Fisher Scientific). For each analysis, we loaded ∼2 μg onto the column. Peptides were separated using a 120 min gradient of 5 to 29% acetonitrile in 0.125% formic acid with a flow rate of 450 nL/min. The scan sequence began with an Orbitrap MS1 spectrum with the following parameters: resolution 60K, scan range 350-1350, automatic gain control (AGC) target 100%, maximum injection time 50ms, and centroid spectrum data type. We use a cycle time of 1s for MS2 analysis which consisted of HCD high-energy collision dissociation with the following parameters: resolution 50K, AGC 200%, maximum injection time 120ms, isolation window 0.6 Th, normalized collision energy (NCE) 35%, and centroid spectrum data type. Dynamic exclusion was set to automatic. The FAIMS compensation voltages (CV) used were -40V/-60V/-80V and -30V/-50V/-70V for each set of 12 non-adjacent superfractions, and additional injection was made for the first 12 non-adjacent superfractions using a CV set of -35/-55/-75V. A 1sec TopSpeed cycle was used for each CV.

### Proteomic data processing

Spectra were converted to mzXML via MSconvert ^161^. Database searching included all entries from the mouse UniProt reference Database (downloaded: August 2021). The database was concatenated with one composed of all protein sequences for that database in the reversed order. Searches were performed using a 50-ppm precursor ion tolerance for total protein level profiling. The product ion tolerance was set to 0.03 Da. These wide mass tolerance windows were chosen to maximize sensitivity in conjunction with Comet searches and linear discriminant analysis ^177,178^. TMTpro labels on lysine residues and peptide N-termini +304.207 Da, as well as carbamidomethylation of cysteine residues (+57.021 Da) were set as static modifications, while oxidation of methionine residues (+15.995 Da) was set as a variable modification. Peptide-spectrum matches (PSMs) were adjusted to a 1% false discovery rate (FDR) ^179,180^. PSM filtering was performed using a linear discriminant analysis, as described previously ^178^ and then assembled further to a final protein-level FDR of 1% ^179^. Proteins were quantified by summing reporter ion counts across all matching PSMs, also as described previously ^181^. Reporter ion intensities were adjusted to correct for the isotopic impurities of the different TMTpro reagents according to manufacturer specifications. The signal-to-noise (S/N) measurements of peptides assigned to each protein were summed and these values were normalized so that the sum of the signal for all proteins in each channel was equivalent to account for equal protein loading. Finally, each protein abundance measurement was scaled, such that the summed signal-to-noise for that protein across all channels equals 100, thereby generating a relative abundance (RA) measurement.

### Proteomic analysis

Protein-level abundances were derived from TMT reporter ion intensities (single TMT 18-plex). Proteins identified as contaminants were excluded, and proteins with any missing values across samples were removed, yielding 7,162 quantified proteins for downstream analysis. Remaining protein abundances were mean-centered across samples to correct for differences in total loading. Two samples were identified as having elevated expression of skeletal muscle marker proteins, likely reflecting procedural contamination from adjacent tissue during dissection. To mitigate this, a binary contamination indicator was regressed out from the protein expression matrix via linear residualization prior to differential expression analysis. Differentially abundant proteins between donor’s WAT before and after transplantation were identified separately for each profiled experimental group using the limma package ^160^, with donor identity included as a fixed-effect covariate to account for within-donor variability. For old-to-young and old-to-old groups, three paired donor–recipient sets (n = 3 pairs per group) were used. To identify proteins with differential transplantation responses between experimental groups, a limma interaction model was fit to all old-to-young and old-to-old samples jointly, with donor pair, transplantation state (before/after), host age group (young/old), and the state × age group interaction as terms. The interaction coefficient captures the differential transplant effect between host ages. P-values were adjusted for multiple testing using the Benjamini-Hochberg method within each contrast.

We conducted functional enrichment analysis to characterize the proteome changes in donor’s WAT induced by transplantation across different experimental groups, and compare them to established transcriptomic signatures of aging and mortality. We applied reference signatures from tissue-specific aging biomarkers of brain, kidney, and liver, and from multi-tissue signatures of chronological age and expected mortality in rodents. For each experimental transplantation group, proteins were ranked by their log2 fold change (logFC) during transplantation. Ranked protein lists were then subjected to gene set enrichment analysis (GSEA) using the clusterProfiler and ReactomePA ^163^ packages in R, with gene sets drawn from the HALLMARK collection of the Molecular Signatures Database (MSigDB), KEGG pathways, and Reactome pathways. The same enrichment pipeline was applied in parallel to the corresponding WAT RNA-seq signatures to enable direct comparison of pathway-level responses across platforms. P-values were adjusted for multiple testing with the Benjamini-Hochberg method (FDR). We computed Pearson correlations between protein-level logFC values and transcript-level logFC values across overlapping genes to quantify similarities between proteomic signatures and transcriptomic ones.

To evaluate cross-platform concordance, a concordance analysis was performed on the WAT rejuvenation signature, defined as the 172 genes whose expression changes correlated significantly with tAge changes according to the Mortality Transcriptomic Clock (BH-adjusted p < 0.05). For each gene, delta-delta values were computed as the difference between old-to-young and old-to-old logFC at both the protein and transcript levels. Genes were classified as concordant if the sign of the delta-delta was the same in both platforms.

To infer shifts in cellular composition from bulk proteomic data, cell-type marker scores were computed for each sample. Adipocyte identity was assessed using the mean Z-scored residualized protein abundance of manually curated adipocyte markers (Plin1, Adipoq, Cidec, Lpl, Aqp7, Retn), and macrophage infiltration was assessed using the mean Z-scored residualized protein abundance of macrophage markers (Adgre1, Cd68, Ctss, Lyz2, Itgam, Csf1r, Lgals3). Residualized values were obtained after regressing out the muscle contamination indicator. Differences between donor WAT before and after transplantation were assessed using paired t-tests within donor–recipient pairs.

Analyses were performed in R v4.3.3 using limma v3.58.1, clusterProfiler v4.10.1, ReactomePA v1.46.0, and msigdbr v10.0.2.

### Metabolite profiling and mass spectrometry

Snap-frozen tissues were weighed and pulverized prior to metabolite extraction. Metabolites were extracted using 1:1:1 methanol:water:chloroform containing d8-phenylalanine, 16:0 lyso-phosphatidylethanolamine (LPE 16:0; Avanti Polar Lipids, cat. no. 856705), and N-heptadecanoyl-D-erythro-sphingosylphosphorylcholine (SM d18:1/17:0; Avanti Polar Lipids, cat. no. 860585), which were spiked into the extraction solvent as internal standards to monitor extraction efficiency and instrument performance. Extraction and reconstitution volumes were scaled to tissue wet weight. Samples were vortexed for 30 seconds, sonicated in ice-cold water for 20 minutes, and centrifuged at 2,030 × g for 30 minutes at 4 °C. The upper aqueous phase was transferred to a clean tube and dried under vacuum using a SpeedVac. The lower non-polar phase was transferred to a clean glass tube using a glass Pasteur pipette and dried under a stream of nitrogen.

Dried polar extracts were resuspended in 1:1 acetonitrile:water at 1.5 µL per mg of starting tissue and centrifuged at 21,000 × g for 15 minutes at 4 °C. Dried non-polar extracts were resuspended in 9:1 methanol:chloroform at 1.5 µL per mg of starting tissue, vortexed for 30 seconds, sonicated for 5 minutes on ice, and centrifuged at 2,030 × g for 5 minutes at 4 °C. Because extraction and reconstitution volumes were scaled to starting tissue mass, peak areas were treated as normalized to tissue wet weight, and no additional tissue-weight normalization was applied after acquisition. Supernatants were transferred to vials for LC-MS analysis.

Polar metabolites were separated using an Infinity 1290 HPLC system coupled to a 6546 Q-TOF mass spectrometer equipped with an electrospray ionization source (Agilent Technologies) and operated in negative ion mode. Chromatographic separation was performed using a 150 × 2.1 mm iHILIC-(P) Classic polymeric column equipped with a 2.1 × 20 mm iHILIC-(P) Classic Guard column (both 5 μm, 200 Å; HILICON AB). Mobile phase A consisted of 20 mM ammonium carbonate in water with 0.1% ammonium hydroxide, and mobile phase B consisted of 100% acetonitrile. The gradient was performed at a flow rate of 0.15 mL/min unless otherwise noted, as follows: 0–23 minutes, a linear gradient from 95% to 5% B; 23–25 minutes, a hold at 5% B; 25–25.5 minutes, a gradient to 95% B at 0.20 mL/min with flow directed to waste; 25.5–32.5 minutes, a hold at 95% B; and 32.5–33 minutes, a hold at 95% B at 0.15 mL/min. The injection volume was 3 μL, and mass spectrometric data were acquired in full-scan mode.

Non-polar metabolites were separated using an Infinity 1290 HPLC system coupled to a 6546 Q-TOF mass spectrometer equipped with an electrospray ionization source (Agilent Technologies). Chromatographic separation was performed using a 150 × 2.1 mm Kinetex EVO C18 column (2.6 μm, 100 Å; Phenomenex). Mobile phase A consisted of 95:5 water:methanol containing 5 mM ammonium acetate at pH 7.5, and mobile phase B consisted of 90:10 isopropanol:methanol. The gradient was performed at a flow rate of 0.15 mL/min unless otherwise noted, as follows: 0–3 minutes, a linear gradient from 15% to 40% B; 3–20 minutes, a linear gradient from 40% to 95% B; 20–25 minutes, a hold at 95% B; 25–26 minutes, a linear gradient from 95% to 15% B; 26–29 minutes, a hold at 15% B at 0.20 mL/min; and 29–30 minutes, a hold at 15% B at 0.15 mL/min. The injection volume was 3 μL.

Each sample was analyzed using two separate RPLC-MS injections with the same column, mobile phases, and chromatographic gradient: one acquisition in negative ion mode and one acquisition in positive ion mode. Data were acquired in full-scan mode for both injections. Positive-mode data were used to profile phosphatidylcholine, phosphatidylethanolamine, diacylglycerol, triacylglycerol, cholesteryl ester, sphingomyelin, ceramide, and acylcarnitine species, together with selected steroids. Negative-mode data were used to profile phosphatidylinositol, phosphatidylglycerol, phosphatidylserine, phosphatidic acid, and cardiolipin species.

Feature extraction and peak integration were performed using MassHunter Profinder version 10.0 (Agilent Technologies) for both polar and non-polar data. Compounds were annotated by matching measured accurate mass within a 5 ppm search tolerance and retention time against in-house libraries of pure chemical standards. Peaks were manually curated to exclude low-quality signals, including noise, improperly integrated or multiple peaks, and compounds for which no valid peaks were identified.

### Metabolomic analysis

#### Data preprocessing

Raw peak area data were extracted from three LC-MS platforms: reversed-phase LC in positive ionization mode (RPLC-POS), reversed-phase LC in negative ionization mode (RPLC-NEG), and hydrophilic interaction LC in negative ionization mode (pHILIC-NEG), capturing lipid and polar metabolite fractions. Internal standards and all reagent-derived features (RX_ prefix) were removed prior to analysis. Zero values were treated as below-detection-limit and replaced with NA. Features were retained if at least two of three samples within every experimental group had valid measurements. Remaining missing values were imputed using half the minimum observed value for that feature across all samples, consistent with a missing-not-at-random mechanism. Peak areas were log2-transformed to stabilize variance. For the RPLC-NEG platform, probabilistic quotient normalization (PQN) ^182^ was applied prior to log2 transformation to correct for sample-level loading differences identified during quality control. For multivariate analyses (PCA), log2-transformed values were additionally Pareto-scaled ^183^. Analyses were performed in R v4.3.0.

#### Fold-change computation and correlation of transplantation responses

For each metabolite, per-pair log2 fold changes (deltas) were computed across four transplantation arms: old fat into young host (O→Y, the rejuvenation arm), old fat into old host (O→O), young fat into young host (Y→Y), and young fat into old host (Y→O). Each delta was defined as the difference between the post-transplantation graft and its matched pre-transplantation baseline (recipient minus donor), averaged across the three pairs. To compare the overall metabolic response across arms, pairwise Pearson correlations were computed between the arm-level delta profiles — the vectors of per-metabolite mean log2 fold changes across all measured metabolites — for the four transplantation arms, with p-values corrected for multiple testing using the BH procedure.

#### Identification of rejuvenation-associated metabolites

Differential metabolite analysis was performed using the limma package, which applies empirical Bayes shrinkage to stabilize per-feature variance estimates across all measured metabolites, an approach particularly suited to small-sample metabolomics (n = 3 per group, 24 total). Log2-transformed, unscaled intensity values from the combined platform matrix (320 metabolites after quality filtering and internal standards removal) were used as input. A linear model was fitted with experimental group as a categorical factor using a means parameterization (no intercept), and donor-recipient pairing was accounted for via the duplicateCorrelation function in limma, which estimates the average within-pair correlation and incorporates it as a weighting term in the model fit. Four transplant arm contrasts were defined, each representing the metabolic shift from donor to recipient: old-to-young (O→Y), old-to-old (O→O), young-to-young (Y→Y), and young-to-old (Y→O). Moderated t-statistics were computed using the eBayes function, and p-values were corrected for multiple testing using the Benjamini-Hochberg procedure across all 320 metabolites within each contrast. Metabolites with adjusted p < 0.10 in the O→Y contrast were considered significantly altered. To characterize the specificity of O→Y metabolic changes, each significant metabolite was classified by its behavior in the Y→O arm: metabolites showing the opposite direction of change in Y→O were classified as divergent, those significant in Y→O with the same direction were classified as a shared transplant response, and those not reaching significance in Y→O (adjusted p >= 0.10) with the same direction were classified as absent. Results were visualized as a scatter plot of O→Y versus Y→O log2 fold changes, and as a heatmap of log2 fold changes across all four arms with significance annotations to display the overall structure of metabolomic changes.

### Tissue preparation for histology and immunostaining

Adipose tissue samples fixed in 3.5% paraformaldehyde overnight were washed with PBS and transferred to 70% ethanol for storage. Samples were paraffin-embedded and subjected to histopathological staining at the Harvard Rodent Histopathology Core. Hematoxylin and eosin (H&E) and Masson’s Trichrome staining were performed. Automated image analysis was performed using the Segment Anything Model (SAM). Adipocyte sizes were obtained by averaging the sizes of 50–200 randomly selected adipocytes per sample (similar yields per fixed field of view, resembling biological differences in adipocyte sizes). Collagen deposition was quantified using SAM applied to Masson’s Trichrome-stained slides. Images for both assays were captured at 10× magnification, with three random fields per sample analyzed.

### Adipocyte lipid droplet quantification

Adipocyte lipid droplet size was quantified from H&E-stained sections using automated segmentation with the Segment Anything Model (SAM) ^164^ employing the ViT-H architecture. Images acquired at 10× magnification were processed using custom Python scripts with the following parameters: minimum mask region area of 0 pixels, predicted intersection over union threshold of 0.5, and stability score threshold of 0.8. Adipocyte lipid droplets were identified as regions with mean RGB values ≥190 for all color channels and areas exceeding 10,000 pixels². Masks touching image borders within 20 pixels were excluded to eliminate incomplete adipocytes. Overlapping masks were resolved by retaining the largest mask. Cell nuclei were distinguished from adipocytes based on their characteristic purple staining (RGB values: R ≤150, G ≤150, B ≤200) and smaller size (<5,000 pixels²). The cross-sectional area of each adipocyte was recorded in pixels². Adipocyte droplet size values were log-transformed prior to statistical analysis to approximate a normal distribution. For each mouse, the mean and standard deviation of droplet size (in the log scale) were calculated, along with the corresponding standard errors. A mixed-effects model using the REML method was applied to assess statistical differences in both the mean and the standard deviation of droplet size before and after transplantation individually for each group. Pairwise differences in adipocyte size dynamics across experimental groups during transplantation were evaluated using a mixed-effects model with transplantation stage, experimental group, and their interaction as fixed effects. The p-value for the interaction term, adjusted for multiple comparisons using the BH method, was used to determine the significance of differences in changes in adipocyte size mean and standard deviation following transplantation between experimental groups.

### Collagen deposition quantification

Collagen content was quantified from Masson’s trichrome-stained sections by identifying blue-stained collagen fibers using automated segmentation with the Segment Anything Model (SAM). Pixels were classified as collagen-positive when RGB values fell within the range of 170-220 for red, 190-240 for green, and 210-255 for blue channels. The total number of collagen-positive pixels was calculated for each image field. Collagen content was expressed as the proportion of collagen-positive pixels relative to the total tissue area after excluding white spaces corresponding to adipocyte lipid droplets. All image analyses were performed using Python with OpenCV and NumPy libraries, and morphometric data were exported as CSV files for subsequent statistical analysis. Collagen content values were log-transformed prior to statistical analysis to approximate a normal distribution. Changes in collagen content during transplantation, as well as differences in these dynamics across experimental groups, were assessed using the same statistical approach described previously for adipocyte droplet size.

### Mitochondria immunostaining

For the mitochondria morphology analyses, we performed an immunofluorescence assay in paraffin slices of 100 and 15 μm, using the previously described protocol ^184^. The WAT sections were blocked in 5% Bovine Serum Albumin (BSA) with 0.5% Triton X-100 for 1 hour at room temperature before being incubated with the primary antibody. For the mitochondria staining, we used the primary antibody Tomm20 (Novus, NBP2-67501, lot number HP0128) diluted 1:100 in PBS Triton X-100 0.25%, incubated overnight at 4°C in a humidified chamber. Sections were washed in PBS 1x before being incubated with secondary antibody Alexa Fluor anti-rabbit 647 (1:500, Thermo, A21244, lot 2659317) with DAPI (1:1000, Sigma-Aldrich, D9542) diluted in 0.1% Triton X-100. Slides were mounted with ProLong Glass Antifade Mountant (Invitrogen, P36980, Lot 2801574) and kept at 4 °C overnight before imaging. All tissues from the same group (WAT Before, After and Native) were stained within the same batch to ensure maintained consistency for image acquisition and analysis.

### Imaging and mitochondria morphology quantification

To assess mitochondrial properties, we performed immunostaining of paraffin-embedded sections using an antibody against Tomm20. Initial imaging of 100 μm sections confirmed homogeneous mitochondrial labeling throughout the adipocyte cytoplasm, and after that, we used 15 μm sections for quantitative analysis (**Figure 5E)**. To exclude infiltrating immune cells, we focused exclusively on mitochondria within the cytoplasmic compartment, omitting those near nuclei (**Supplementary Figure 5D**). Following segmentation of cytoplasmic mitochondria, we quantified their numbers, as well as total volume and sphericity as readouts of mitochondrial morphology. Because cell boundaries were not always clearly discernible, we estimated mitochondrial numbers by quantifying mitochondrial density by dividing the total number of segmented mitochondria by the total cytoplasmic volume of adipocytes in the region of interest. Three regions per tissue sample were randomly selected and imaged using a Yokogawa CSU-W1 spinning disk confocal unit (50 µm pinhole size) mounted on a fully motorized Nikon Ti2 inverted microscope equipped with a Nikon linear-encoded motorized XY stage and a Physik Instrumente piezo Z-motor integrated into the nosepiece drive. An Andor Zyla 4.2 Plus sCMOS (6.5 µm pixel size) monochrome camera was used together with a Nikon Plan Fluor 40×/1.3 NA oil-immersion objective (Cargille Type 37 immersion oil). Images were acquired at 1×1 binning, 12-bit depth (Gain 1). Excitation for confocal imaging was provided by solid-state, directly modulated diode lasers launched through a Nikon LUN-F laser combiner, using the 405 nm, 488 nm, or 640 nm laser lines depending on the fluorophore. A hard-coated Semrock Di01-T405/488/568/647 multiband dichroic mirror was used for all laser-based channels. Green fluorescence excited with the 488 nm laser was collected using a Chroma ET535/36 nm emission filter, and far-red fluorescence excited with the 640 nm laser was collected through a Chroma ET705/72 nm emission filter. No software-based autofocus routines were used. For image acquisition, Nikon Elements AR (version 5.41.02) was used. Z-stacks were collected using either the Nikon nosepiece drive or the PI piezo Z-motor. For each field of view, images were acquired sequentially by channel, with the shutter remaining open during exposure. For each sample, a complete optical volume was acquired, typically between 41 and 50 Z-slices collected at a 0.3 µm step size. Data were saved in ND2 format. Prior to morphological analysis, all images were pre-processed in Fiji. For standardization of three-dimensional dimensions across samples, each image was cropped to include 41 optical slices, corresponding to a total depth of 12.3 μm. Image segmentation and object detection, such as mitochondria, nuclei, and cell membranes, were performed using Arivis software. Cell membranes were identified by the autofluorescence signal captured with the 488 nm laser. The three-dimensional mitochondrial morphology was assessed in terms of mitochondrial volume and sphericity. Mitochondrial volume values were log-transformed before statistical analysis to approximate a normal distribution. Changes in mitochondrial volume and sphericity during transplantation were assessed using the same statistical approach described previously for adipocyte droplet size.

### Fibrosis assessment

White adipose tissue (WAT) samples from all experimental groups were submitted to the Rodent Histopathology Core at Harvard Medical School for blinded fibrosis evaluation. A trained veterinary histopathologist, blinded to donor age, recipient age, and transplantation group, performed qualitative and semi-quantitative scoring of fibrosis. All slides were evaluated under identical conditions, and fibrosis scores were recorded and returned to the research team without disclosure of sample identity until the completion of analysis.

### Drosophila stocks and husbandry

Wild-type *Canton-S* (a gift from Norbert Perrimon lab) and whole body–fat body–GeneSwitch-Gal4 (WB-FB-GS-GAL4) flies (a gift from John Tower lab) were maintained on standard cornmeal medium (“Fly Food J”; LabExpress, Ann Arbor, MI) containing yeast, dextrose, sucrose, and molasses, at 25 °C under a 12-h light/dark cycle. Drug-inducible, fat body-specific transgene expression was achieved by crossing a whole-body fat-body GS-GAL4 driver to the following UAS responder lines, all obtained from the Bloomington *Drosophila* Stock Center (BDSC): UAS-RNAi against SERCA (the single *Drosophila* SERCA orthologue; BDSC #44581) and against *Lrmda* (BDSC #60478) for knockdown, and a UAS-SERCA line for overexpression (BDSC #63228).

### Lifespan assays

Flies were collected within 24–48 h of eclosion, allowed to mate for 1–2 days, and sorted by sex under CO₂ anesthesia. Sorted flies were housed at 20–25 per vial at 25 °C and 60% relative humidity under a 12-h light/dark cycle, transferred to fresh vials every two days, and scored for deaths at each transfer. Transgene expression in GS lines was induced by adding RU-486 (mifepristone; Cayman Chemical, no. 10006317) dissolved in ethanol to the food at a final concentration of 200 µM (86 µg/mL) (GeneSwitch system ^185^); control flies received food with an equal volume of ethanol vehicle. For constitutive experiments, RU-486 was present throughout adult life. For intermittent experiments, RU-486 was provided to females continuously over the first 30 days of adult life (days 0–30) and to males in two 10-day pulses separated by a 30-day withdrawal (days 0–10 and 40–50), with no further induction thereafter; matched control flies received vehicle on the same schedule.

### Survival analysis

Each fly was recorded as a single observation with its day of death and event indicator; flies alive at the final recorded time point were treated as right-censored. Survival distributions were estimated by the Kaplan–Meier method ^168^ for each sex separately and for both sexes combined, and overall survival was compared with the log-rank test; sex was included as a covariate in the combined-sex models. Median lifespan and its 95% confidence interval were obtained from the Kaplan–Meier estimate using the Greenwood formula. Maximum lifespan, defined as the 90^th^ percentile of the survival-time distribution, was compared between groups by quantile regression for survival data (quantreg ^162^) with standard errors from 1,000 bootstrap iterations.

Summary hazard ratios were estimated with Cox proportional-hazards models ^169^ with sex included as a covariate in combined-sex analyses. In the constitutive SERCA overexpression experiment, which was carried out across multiple batches, Cox and log-rank models were stratified by batch to account for clustering within replicates; the remaining experiments were performed as single batches and were not stratified. To characterize the temporal dynamics of each effect and to assess the proportional-hazards assumption, Cox models with time-varying coefficients were fitted using the tt() interface, specifying the interaction between group and log-transformed time [log(t + 1)]; a positive interaction coefficient denotes a hazard ratio that increases over follow-up (an early protective effect that attenuates, or an early detrimental effect that accumulates), and a negative coefficient the converse. P-values across sexes and lifespan metrics were adjusted by the BH procedure, with significance defined at p.adjusted < 0.05. Analyses were performed in R v4.3.0 using the survival, survminer, quantreg, and dplyr packages.

### Complete Blood Count (CBC) analysis

Data were pooled from three independent studies (Study 1: n = 54; Study 2: n = 10; Study 3: n = 47; combined N = 111). Animals were assigned to one of eight groups: Young sham, Young donor, Young recipient of young fat, Young recipient of old fat, Old sham, Old donor, Old recipient of young fat, and Old recipient of old fat. Young animals had a chronological age of approximately 177–201 days and old animals approximately 603–657 days at the time of measurement.

#### Outlier detection

Prior to statistical analysis, a principal component analysis (PCA) was performed on all 20 complete blood count (CBC) parameters to identify multivariate outliers. CBC values were standardised (z-scored) before PCA. Mahalanobis distances were computed in the space of the first five principal components (accounting for 83% of total variance) and compared against a chi-squared distribution (df = 5) at a threshold of p < 0.001. Three animals exceeded this threshold and were excluded from all subsequent analyses: OOR4 (Study 2, Old recipient of old fat; extreme on PC3, characterised by markedly low RBC), YD5 (Study 1, Young donor; extreme on PC2, characterised by markedly elevated WBC and neutrophil counts), and YYR4 (Study 1, Young recipient of young fat; borderline outlier on PC2). The final analytical dataset comprised N = 108 animals.

#### The dynamical frailty index parameters

The dynamical Frailty Index (dFI) and derived parameters (dFI_loghaz, dFI_perc, dFI_age) were computed from CBC measurements as described previously^53^.

#### Group comparisons

For all pairwise group comparisons, analysis of covariance (ANCOVA) was used with study batch and chronological age as fixed-effect covariates (Type II sums of squares). Chronological age was included as a covariate in all comparisons among young groups (age range 177–201 days) and all comparisons among old groups (age range 603–657 days), given the modest within-category variation in age. For comparisons between young and old sham animals, chronological age was not included as an additional covariate because it was nearly perfectly collinear with group membership (Pearson’s r = 0.998, VIF > 350), precluding independent estimation of the group effect; study batch alone was used as the covariate in those models. Effect sizes are reported as Cohen’s d, calculated as the difference in group means divided by the square root of the residual mean square error from the ANCOVA model. Multiple testing correction was applied across all pairwise comparisons within each analysis using the BH false discovery rate (FDR) procedure.

#### Body weight change

Body weight change was computed as the difference between weight recorded at week 10 and weight recorded at week 0 (baseline). The same ANCOVA framework (study + chronological age as covariates, FDR correction) was applied to body weight change within each age cohort.

#### CBC multivariate analysis

The effect of intervention on the 20 CBC parameters (WBC, NE, LY, MO, EO, BA, NE%, LY%, MO%, EO%, BA%, RBC, Hb, HCT%, MCV, MCH, MCHC, RDW%, PLT, MPV) was assessed in two stages. First, a multivariate analysis of covariance (MANOVA) was conducted with all 20 parameters as simultaneous outcomes and group, study, and chronological age as predictors; significance was assessed using Pillai’s trace. Second, regardless of the omnibus result, individual univariate ANCOVAs were run for each parameter, with FDR correction applied across all 20 tests within each age cohort (k = 20). Effect sizes for each parameter relative to the sham reference group are presented as Cohen’s d in a heatmap.

#### Correlations with dFI_perc

Spearman rank correlations were computed between dFI_perc and each of the 20 CBC parameters for all animals combined and separately within young and old cohorts. FDR correction (Benjamini–Hochberg) was applied within each subset (k = 20). The RBC–dFI_perc relationship was further examined via scatter plots with linear regression lines overlaid.

#### Software

All analyses were performed in Python 3.12. Key packages: pandas (v2.x) for data management; statsmodels (v0.14) for ANCOVA and MANOVA; scipy (v1.11) for Spearman correlations; scikit-learn (v1.3) for PCA; matplotlib (v3.10) for figure generation. Multiple testing correction used the BH procedure implemented in statsmodels.stats.multitest.

---

**Supplementary Figure 1.**
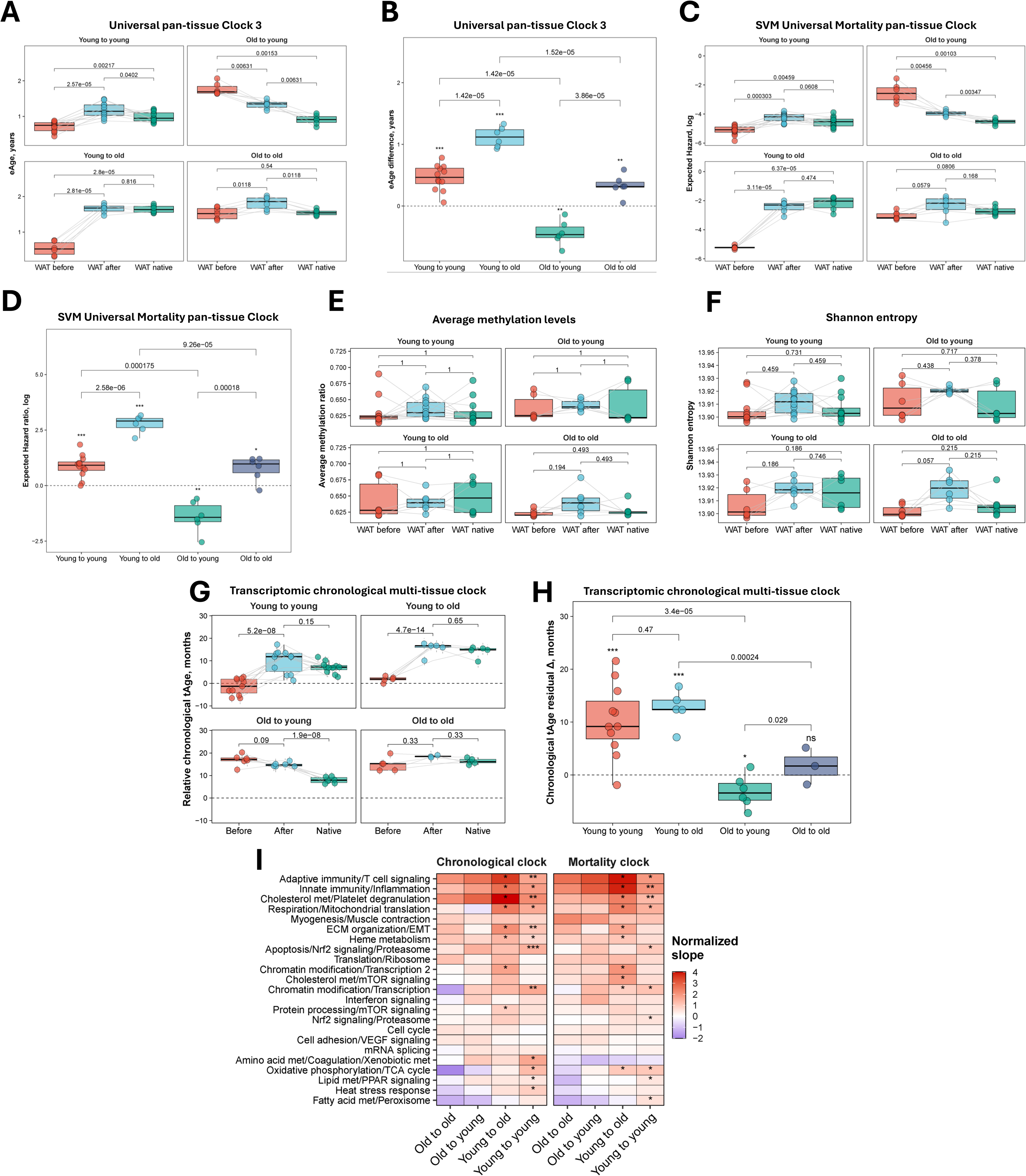
Epigenetic and transcriptomic aging clocks reveal rejuvenation of aged WAT in young hosts. **(A)** Epigenetic age (eAge) of WAT samples before and after transplantation across all experimental groups according to the Universal Clock 3 predictions. Mouse ID was included in the statistical model as a covariate to ensure paired comparison of eAge dynamics. Each dot represents a sample collected from individual mice, n = 6-12 per group. **(B)** Comparison of epigenetic age (eAge) changes of WAT samples before and after transplantation across all experimental groups according to Universal Clock 3. Asterisks reflect statistical significance (BH-adjusted p-values) of eAge changes during transplantation within individual groups, whereas the significance of pairwise comparisons in eAge dynamics between groups is denoted in text. **(C)** Expected hazard ratios (in log scale) of WAT samples before and after transplantation across all experimental groups according to the SVM Universal Mortality pan-tissue Clock. Mouse ID was included in the statistical model as a covariate to ensure paired comparison of expected hazard ratio dynamics. Each dot represents a sample collected from individual mice, n = 6-12 per group. **(D)** Comparison of expected hazard ratio (in log scale) changes of WAT samples before and after transplantation across all experimental groups according to the SVM Universal Mortality pan-tissue Clock. Asterisks reflect statistical significance (BH-adjusted p-values) of expected hazard ratio changes during transplantation within individual groups, whereas the significance of pairwise comparisons in expected hazard ratio dynamics between groups is denoted in text. **(E-F)** Average methylation levels **(E)** and Shannon entropy **(F)** of WAT samples before and after transplantation across all experimental groups. Mouse ID was included in the statistical model as a covariate to ensure paired comparison of average methylation levels and Shannon entropy. Each dot represents a sample collected from individual mice, n = 6-12 per group. **(G)** Transcriptomic age (tAge) of WAT samples before and after transplantation across all experimental groups according to the Bayesian Ridge (BR) rodent multi-tissue chronological clock. Statistical significance was assessed with mixed-effects models. Mouse ID was included in the statistical model as a covariate to ensure paired comparison of tAge dynamics. Data are tAges ± s.d. Each dot represents a sample collected from individual mice, n = 6-11 per group. **(H)** Comparison of age-adjusted changes in transcriptomic age (tAge) of WAT samples before and after transplantation across all experimental groups, according to the rodent multi-tissue transcriptomic chronological clock. Statistical significance was assessed with mixed-effects models. Asterisks reflect statistical significance (BH-adjusted p-values) of tAge changes during transplantation within individual groups, whereas the significance of pairwise comparisons in tAge dynamics between groups is denoted in text. **(I)** Standardized change of tAge in WAT samples following transplantation across all experimental conditions, according to module-specific rodent multi-tissue transcriptomic chronological and mortality clocks. Red color indicates increased tAge, while blue color indicates decreased tAge after transplantation relative to their respective pre-transplantation baselines. Asterisks reflect statistical significance (BH-adjusted p-values) of changes, assessed with ANOVA model with donor fat identity included as a covariate. Mice per group = 6-11. Boxplots: center line indicates median; box limits, interquartile range; whiskers, ±1.5× IQR. Unless specified otherwise, statistical differences between groups are assessed with ANOVA and adjusted for multiple comparisons with the Benjamini-Hochberg approach. *** p.adjusted< 0.001, ** p.adjusted < 0.01, * p.adjusted < 0.05, ^ p.adjusted < 0.1.

**Supplementary figure 2.**
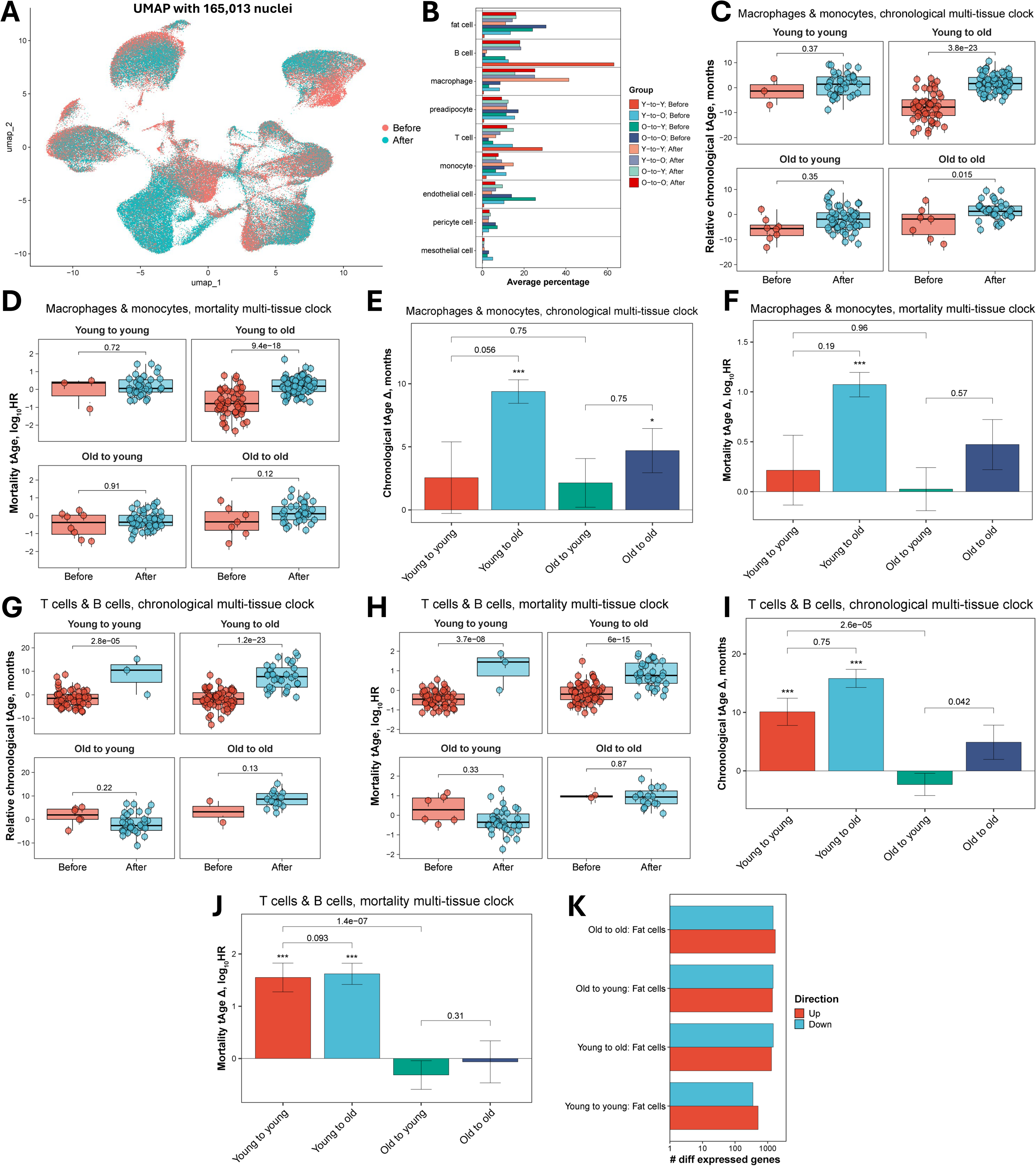
Single-nucleus profiling and module clocks localize and characterize rejuvenation in adipocytes. **(A)** The UMAP from Figure 2A recolored by transplantation timepoint. **(B)** Average percentage of most abundant cell types among all sequenced nuclei across the eight experimental groups, shown as horizontal bars. Cell types are arranged along the y-axis; bar length indicates the average proportional abundance within each group. The eight groups represent the four transplantation conditions (young-to-young, Y-to-Y; young-to-old, Y-to-O; old-to-young, O-to-Y; old-to-old, O-to-O) at each of the two timepoints (Before and After transplantation), distinguished by color as indicated in the legend. **(C)** Transcriptomic age (tAge) of pooled macrophages and monocytes before and after transplantation across all experimental groups according to the Bayesian Ridge (BR) mouse multi-tissue chronological clock. Statistical significance was assessed with mixed-effects models. Mouse ID was included in the statistical model as a covariate to ensure paired comparison of tAge dynamics. Data are tAges ± s.d. Each dot represents a meta-cell compiled from individual mice samples, n = 1-3 per group. **(D)** Transcriptomic age (tAge) of pooled macrophages and monocytes before and after transplantation across all experimental groups according to the BR mouse multi-tissue mortality clock. Statistical significance was assessed with mixed-effects models. Mouse ID was included in the statistical model as a covariate to ensure paired comparison of tAge dynamics. Data are tAges ± s.d. Each dot represents a meta-cell compiled from individual mice samples, n = 1-3 per group. **(E)** Comparison of chronological tAge dynamics in pooled macrophages and monocytes over 10 weeks after transplantation across groups. For each transplantation group, the slope was estimated using a mixed-effects model with donor fat identity included as a covariate; p-values testing whether the slope differs from 0 are indicated with asterisks. Pairwise comparisons of tAge changes between groups were performed using mixed-effects model, with the p-value for the interaction term shown in the plot. P-values were adjusted for multiple comparisons using the Benjamini-Hochberg approach. Data are slope estimate ± SE. **(F)** Comparison of mortality tAge dynamics in pooled macrophages and monocytes over 10 weeks after transplantation across groups. For each transplantation group, the slope was estimated using a mixed-effects model with donor fat identity included as a covariate; p-values testing whether the slope differs from 0 are indicated with asterisks. Pairwise comparisons of tAge changes between groups were performed using mixed-effects model, with the p-value for the interaction term shown in the plot. P-values were adjusted for multiple comparisons using the Benjamini-Hochberg approach. Data are slope estimate ± SE. **(G)** Transcriptomic age (tAge) of pooled T cells and B cells before and after transplantation across all experimental groups according to the BR mouse multi-tissue chronological clock. Statistical significance was assessed with mixed-effects models. Mouse ID was included in the statistical model as a covariate to ensure paired comparison of tAge dynamics. Data are tAges ± s.d. Each dot represents a meta-cell compiled from individual mice samples, n = 1-3 per group. **(H)** Transcriptomic age (tAge) of pooled T cells and B cells before and after transplantation across all experimental groups according to the BR mouse multi-tissue mortality clock. Statistical significance was assessed with mixed-effects models. Mouse ID was included in the statistical model as a covariate to ensure paired comparison of tAge dynamics. Data are tAges ± s.d. Each dot represents a meta-cell compiled from individual mice samples, n = 1-3 per group. **(I)** Comparison of chronological tAge dynamics in pooled T cells and B cells over 10 weeks after transplantation across groups. For each transplantation group, the slope was estimated using a mixed-effects model with donor fat identity included as a covariate; p-values testing whether the slope differs from 0 are indicated with asterisks. Pairwise comparisons of tAge changes between groups were performed using mixed-effects model, with the p-value for the interaction term shown in the plot. P-values were adjusted for multiple comparisons using the Benjamini-Hochberg approach. Data are slope estimate ± SE. **(J)** Comparison of mortality tAge dynamics in pooled T cells and B cells over 10 weeks after transplantation across groups. For each transplantation group, the slope was estimated using a mixed-effects model with donor fat identity included as a covariate; p-values testing whether the slope differs from 0 are indicated with asterisks. Pairwise comparisons of tAge changes between groups were performed using mixed-effects model, with the p-value for the interaction term shown in the plot. P-values were adjusted for multiple comparisons using the Benjamini-Hochberg approach. Data are slope estimate ± SE. **(K)** Number of differentially expressed genes (p.adjusted < 0.05) in a donor’s adipocytes after transplantation relative to their respective pre-transplantation baselines for each experimental group. Barplots indicate the numbers of differentially expressed genes, and colors denote the direction of changes. Mice per group = 1-3. Boxplots: center line indicates median; box limits, interquartile range; whiskers, ±1.5× IQR. Unless specified otherwise, statistical differences between groups are assessed with ANOVA and adjusted for multiple comparisons with the Benjamini-Hochberg approach. *** p.adjusted< 0.001, ** p.adjusted < 0.01, * p.adjusted < 0.05, ^ p.adjusted < 0.1

**Supplementary Figure 3.**
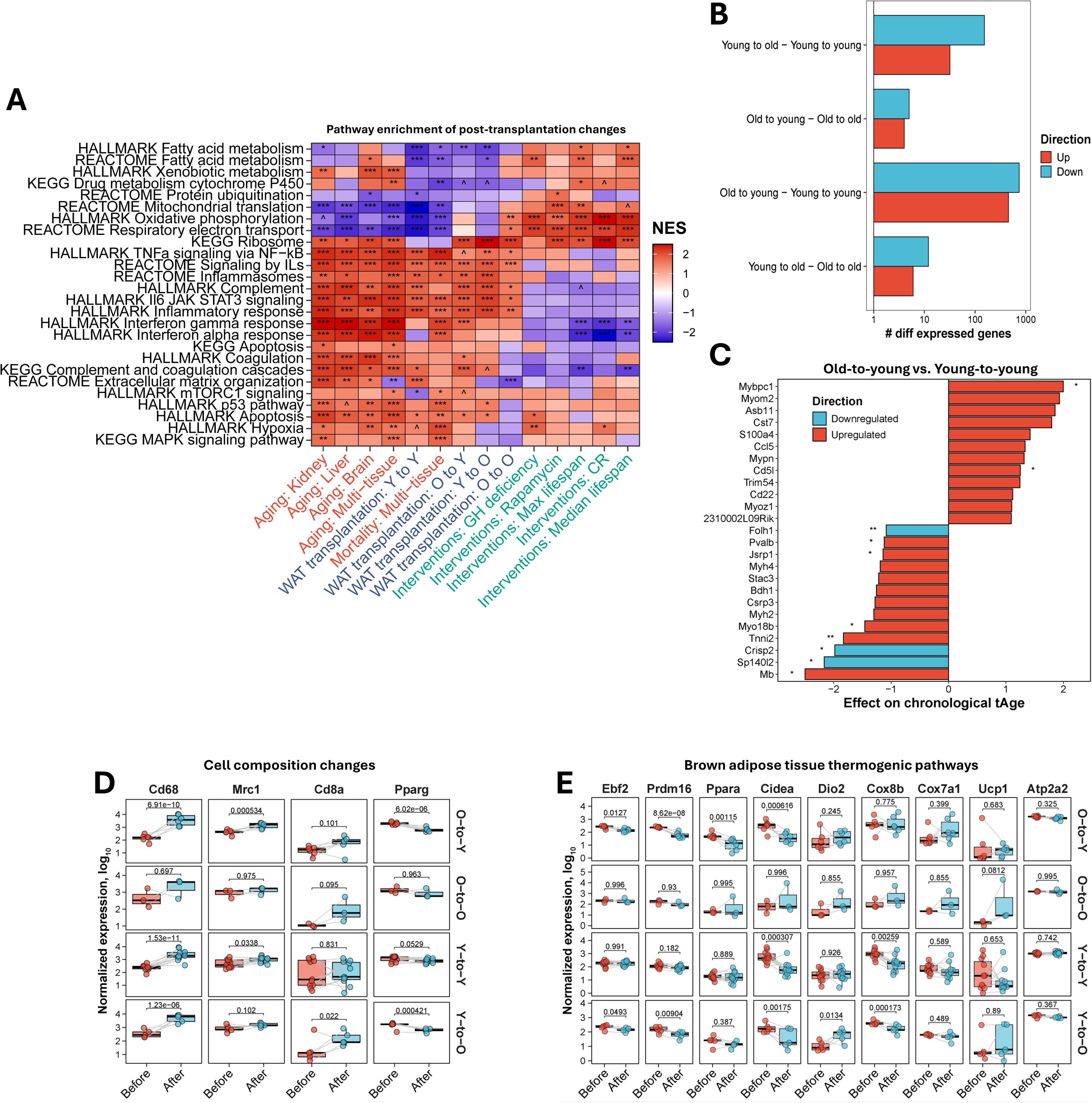
WAT rejuvenation engages thermogenic and muscle-like transcriptional programs. **(A)** Pathway enrichment analysis (GSEA) of gene expression changes induced by transplantation in a donor’s WAT across all experimental groups (darkblue), along with the established signatures of lifespan-extending interventions (green), aging, and mortality in rodents (red). The direction of expression change is reflected with color, and asterisks reflect statistical significance (BH-adjusted p-values). See Supplementary File 1 for full enrichment results. **(B)** Number of genes with differential change of expression in a donor’s WAT samples after transplantation relative to their respective pre-transplantation baselines across all experimental groups. Barplots indicate the numbers of differentially expressed genes, and colors denote the direction of changes. **(C)** Top genes contributing to increased and decreased tAge during transplantation in the old-to-young vs young-to-young group, according to the rodent multi-tissue transcriptomic chronological clock. Color and asterisks denote direction and statistical significance (BH-adjusted p-value) of difference in expression dynamics between groups, respectively. **(D)** Normalized expression dynamics (in log scale) of established immune (Cd68, Mrc1, Cd8a) and adipocyte (Pparg) cell-type markers in WAT samples before and after transplantation across all transplantation models. **(E)** Normalized log-expression dynamics of established brown adipocyte thermogenic markers in WAT samples before and after transplantation across all transplantation models. Boxplots: center line indicates median; box limits, interquartile range; whiskers, ±1.5× IQR. Number of samples from individual mice = 3-11 per group. Unless specified otherwise, statistical differences between groups are assessed with ANOVA and adjusted for multiple comparisons with Benjamini-Hochberg approach. *** p.adjusted< 0.001, ** p.adjusted < 0.01, * p.adjusted < 0.05, ^ p.adjusted < 0.1.

**Supplementary figure 4.**
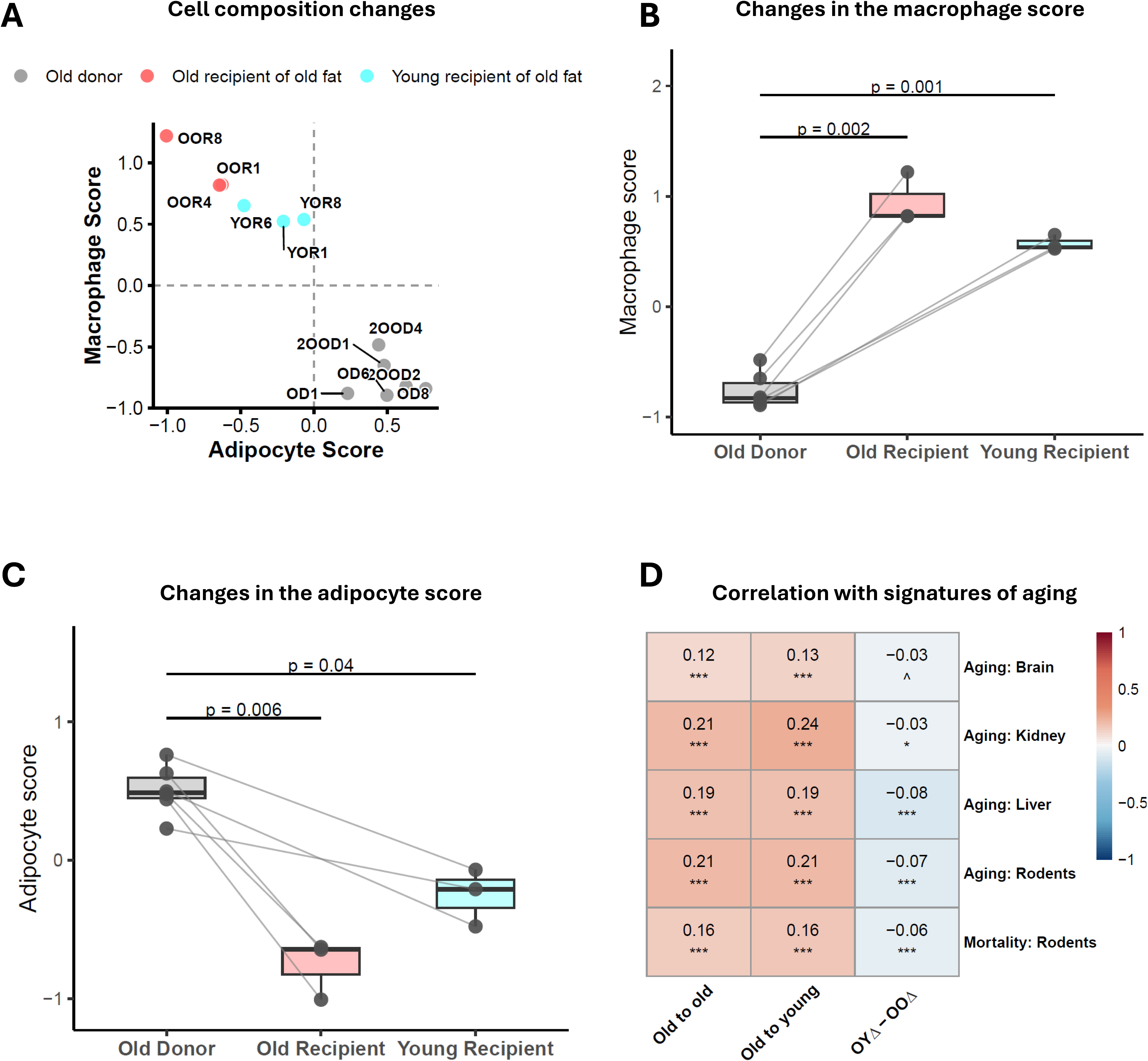
Proteomic and metabolomic profiling corroborate and extend the transcriptomic signature of WAT rejuvenation. **(A)** Scatter plot of adipocyte marker score (x-axis) versus macrophage marker score (y-axis). Grey points represent old donor WAT before transplantation; red points, old WAT recovered from old recipients; cyan points, old WAT recovered from young recipients. **(B)** Macrophage compositional scores in old donor WAT before and after transplantation into old or young recipients. Macrophage marker score (mean Z-score of Adgre1, Cd68, Ctss, Lyz2, Itgam, Csf1r, and Lgals3 protein abundances) in old donor WAT before transplantation (Old Donor) compared to WAT after transplantation into old recipients (Old Recipient) or young recipients (Young Recipient). Lines connect paired donor–recipient samples. Statistical comparisons between paired samples were performed using paired t-tests. **(C)** Adipocyte compositional scores in old donor WAT before and after transplantation into old or young recipients. Adipocyte marker score (mean Z-score of Plin1, Adipoq, Cidec, Lpl, Aqp7, and Retn protein abundances) in old donor WAT before transplantation (Old Donor) compared to WAT after transplantation into old recipients (Old Recipient) or young recipients (Young Recipient). Lines connect paired donor–recipient samples. Statistical comparisons between paired samples were performed using paired t-tests. **(D)** Pearson correlation heatmap of protein-level log2 fold changes (logFC) from paired proteomics signatures (Old to old, Old to young, and the interaction contrast OYΔ–OOΔ) against established transcriptomic signatures of aging and mortality across tissues and species. Values indicate correlation coefficients; asterisks indicate statistical significance (Pearson correlation test with BH-adjusted p-values). Boxplots: center line indicates median; box limits, interquartile range; whiskers, ±1.5× IQR. Number of samples from individual mice = 3 per group. Unless specified otherwise, statistical differences between groups are assessed with ANOVA and adjusted for multiple comparisons with the Benjamini-Hochberg approach. *** p.adjusted< 0.001, ** p.adjusted < 0.01, * p.adjusted < 0.05, ^ p.adjusted < 0.1.

**Supplementary Figure 5.**
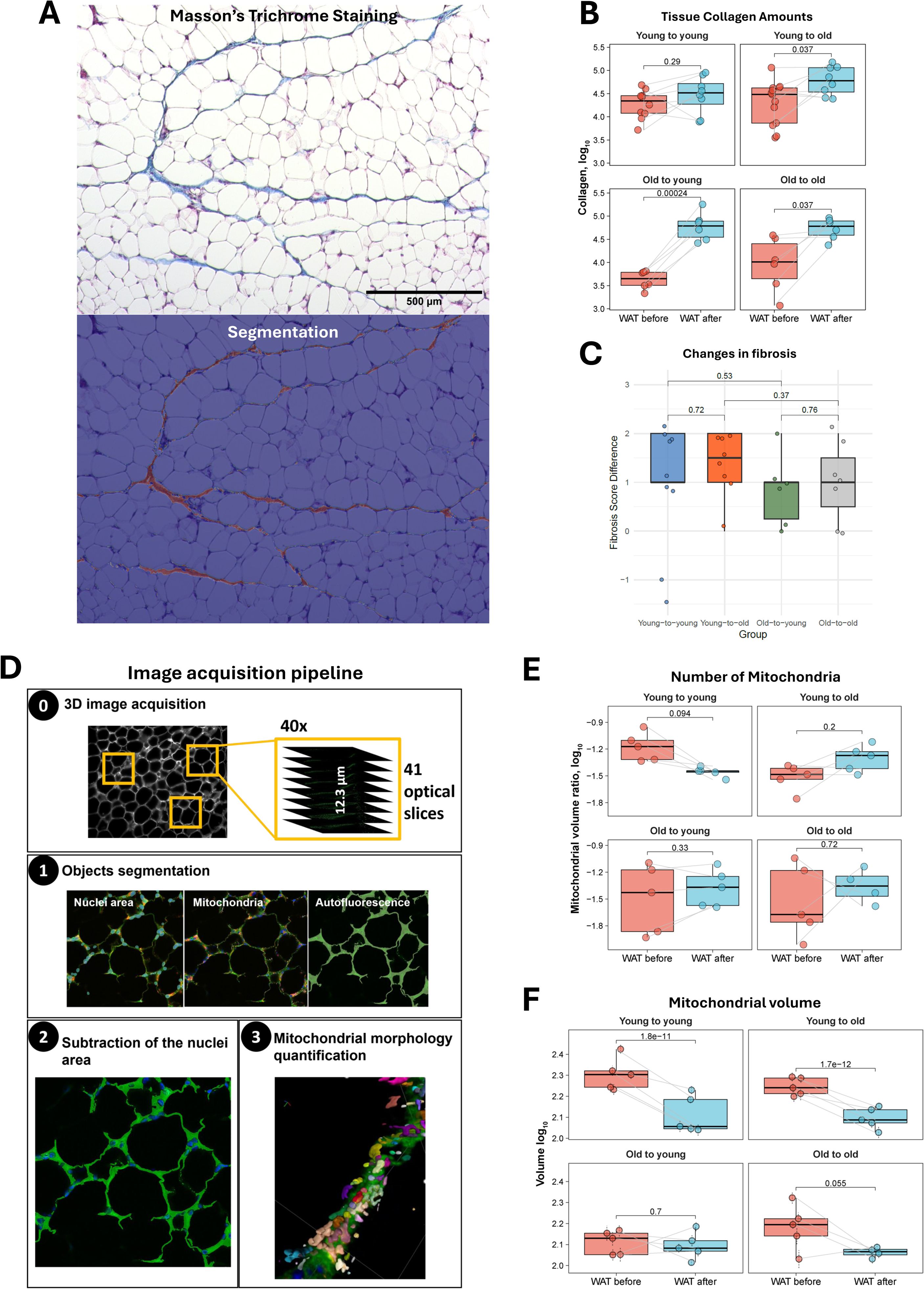
WAT rejuvenation model and validation of findings at the structural level of organization. **(A)** Example image of adipose tissue stained with Masson’s Trichrome (top) and computationally segmented collagen-positive regions used for quantification (bottom). **(B)** Quantification of collagen deposition ratio (in log scale) in WAT samples before and after transplantation across all experimental groups. Changes in collagen deposition ratio were assessed with ANOVA model. Mouse ID was included in the model as a covariate to ensure paired comparison of collagen deposition ratios. Each dot represents a sample collected from individual mice, n = 6-13 per group. **(C)** Comparison of fibrosis score changes in WAT samples before and after transplantation across all experimental groups. Pairwise comparisons were performed using the two-sided Wilcoxon rank-sum test with the Benjamini–Hochberg correction for multiple comparisons. Mice per group = 6-9. **(D)** Workflow for three-dimensional mitochondrial morphology analysis. (0) Three fields per tissue sample were randomly selected and imaged using spinning disk confocal microscopy, acquiring 41 optical slices per field to generate a 12.3 μm-deep 3D image stack. (1) Image segmentation was performed using Arivis software to identify nuclei (DAPI), mitochondria (Tomm20), and cytoplasm (autofluorescence signal from the 488 nm channel). (2) To exclude mitochondria from non-adipocyte cells, nuclear regions were subtracted, and only cytoplasmic regions distant from nuclei were analyzed. (3) Mitochondria were segmented in 3D and their morphological features quantified. **(E)** Quantification of mitochondrial density ratio (number per cytoplasmic volume, in log scale) in all WAT samples before and after transplantation across all experimental groups. Changes in mitochondrial density ratios were assessed with ANOVA model. Mouse ID was included in the model as a covariate to ensure paired comparison of mitochondrial density ratios. Each dot represents a sample collected from individual mice, n = 4-5 per group. **(F)** Quantification of mitochondrial volume (in log scale) in WAT samples before and after transplantation across all experimental groups. Changes of mitochondrial volume following transplantation were assessed with mixed-effects models. Mouse ID was included in the model as a covariate to ensure paired comparison of mitochondrial volume ratios. Data are mean mitochondrial volume (in log scale) ± s.e.m. Each dot represents a sample collected from individual mice, n = 4-5 per group. Boxplots: center line indicates median; box limits, interquartile range; whiskers, ±1.5× IQR. Unless specified otherwise, statistical differences between groups are assessed with ANOVA and adjusted for multiple comparisons with the Benjamini-Hochberg approach. *** p.adjusted< 0.001, ** p.adjusted < 0.01, * p.adjusted < 0.05, ^ p.adjusted < 0.1.

**Supplementary figure 6.**
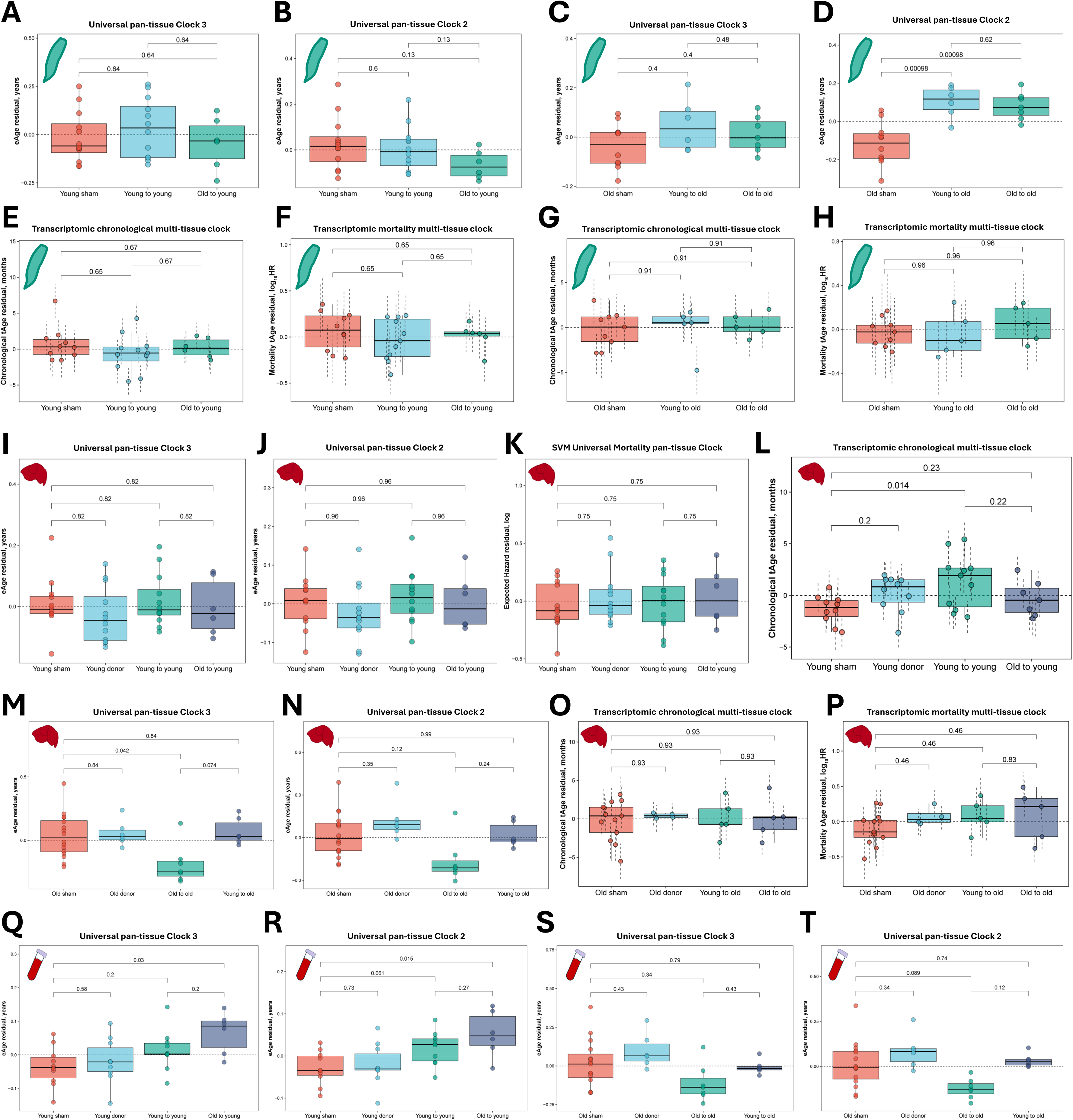
Heterochronic WAT transplantation has a limited impact on host systemic aging. **(A)** Epigenetic age residuals (eAge residual, years) of native WATs from young recipient mice (young-to-young and old-to-young transplantation groups) compared to young sham controls, estimated using Universal pan-tissue Clock 3. Residuals were obtained by regressing out chronological age, sequencing batch, and study as covariates; positive values indicate epigenetic age acceleration relative to the value expected for the given chronological age. BH-adjusted p-values reflect pairwise comparisons between groups. Each dot represents a sample collected from individual mice, n = 6-15 per group. **(B)** Epigenetic age residuals (eAge residual, years) of native WATs from young recipient mice (young-to-young and old-to-young transplantation groups) compared to young sham controls, estimated using Universal pan-tissue Clock 2. Residuals were obtained by regressing out chronological age, sequencing batch, and study as covariates; positive values indicate epigenetic age acceleration relative to the value expected for the given chronological age. BH-adjusted p-values reflect pairwise comparisons between groups. Each dot represents a sample collected from individual mice, n = 6-15 per group. **(C)** Epigenetic age residuals (eAge residual, years) of native WATs from old recipient mice (young-to-old and old-to-old transplantation groups) compared to old sham controls, estimated using Universal pan-tissue Clock 3. Residuals were obtained by regressing out chronological age, sequencing batch, and study as covariates; positive values indicate epigenetic age acceleration relative to the value expected for the given chronological age. BH-adjusted p-values reflect pairwise comparisons between groups. Each dot represents a sample collected from individual mice, n = 6-10 per group. **(D)** Epigenetic age residuals (eAge residual, years) of native WATs from old recipient mice (young-to-old and old-to-old transplantation groups) compared to old sham controls, estimated using Universal pan-tissue Clock 2. Residuals were obtained by regressing out chronological age, sequencing batch, and study as covariates; positive values indicate epigenetic age acceleration relative to the value expected for the given chronological age. BH-adjusted p-values reflect pairwise comparisons between groups. Each dot represents a sample collected from individual mice, n = 6-10 per group. **(E)** Transcriptomic age residuals (tAge residual, months) of native WATs from young recipient mice (young-to-young and old-to-young transplantation groups) compared to young sham controls, estimated using Bayesian Ridge (BR) rodent multi-tissue transcriptomic chronological clock. Residuals were obtained by regressing out chronological age, sequencing batch, and study as covariates; positive values indicate transcriptomic age acceleration relative to the value expected for the given chronological age. BH-adjusted p-values reflect pairwise comparisons between groups, assessed with mixed-effects model. Each dot represents a sample collected from individual mice, n = 6-10 per group. **(F)** Expected hazard ratio residuals (in log scale) (Mortality tAge residual, log_10_HR) of native WATs from young recipient mice (young-to-young and old-to-young transplantation groups) compared to young sham controls, estimated using BR rodent multi-tissue transcriptomic mortality clock. Residuals were obtained by regressing out chronological age, sequencing batch, and study as covariates; positive values indicate transcriptomic age acceleration relative to the value expected for the given chronological age. BH-adjusted p-values reflect pairwise comparisons between groups, assessed with mixed-effects model. Each dot represents a sample collected from individual mice, n = 6-10 per group. **(G)** Transcriptomic age residuals (tAge residual, months) of native WATs from old recipient mice (young-to-old and old-to-old transplantation groups) compared to old sham controls, estimated using BR rodent multi-tissue transcriptomic chronological clock. Residuals were obtained by regressing out chronological age, sequencing batch, and study as covariates; positive values indicate transcriptomic age acceleration relative to the value expected for the given chronological age. BH-adjusted p-values reflect pairwise comparisons between groups, assessed with mixed-effects model. Each dot represents a sample collected from individual mice, n = 5-9 per group. **(H)** Expected hazard ratio residuals (in log scale) (Mortality tAge residual, log_10_HR) of native WATs from old recipient mice (young-to-old and old-to-old transplantation groups) compared to old sham controls, estimated using BR rodent multi-tissue transcriptomic mortality clock. Residuals were obtained by regressing out chronological age, sequencing batch, and study as covariates; positive values indicate transcriptomic age acceleration relative to the value expected for the given chronological age. BH-adjusted p-values reflect pairwise comparisons between groups, assessed with mixed-effects model. Each dot represents a sample collected from individual mice, n = 5-9 per group. **(I)** Epigenetic age residuals (eAge residual, years) of livers from young donor and recipient mice (young-to-young and old-to-young transplantation groups) compared to young sham controls, estimated using Universal pan-tissue Clock 3. Residuals were obtained by regressing out chronological age and study as covariates; positive values indicate epigenetic age acceleration relative to the value expected for the given chronological age. BH-adjusted p-values reflect pairwise comparisons between groups. Each dot represents a sample collected from individual mice, n = 6-12 per group. **(J)** Epigenetic age residuals (eAge residual, years) of livers from young donor and recipient mice (young-to-young and old-to-young transplantation groups) compared to young sham controls, estimated using Universal pan-tissue Clock 2. Residuals were obtained by regressing out chronological age and study as covariates; positive values indicate epigenetic age acceleration relative to the value expected for the given chronological age. BH-adjusted p-values reflect pairwise comparisons between groups. Each dot represents a sample collected from individual mice, n = 6-12 per group. **(K)** Expected hazard ratio residuals (in log scale) (eAge residual, logHR) of livers from young donor and recipient mice (young-to-young and old-to-young transplantation groups) compared to young sham controls, estimated using SVM Universal Mortality pan-tissue Clock. Residuals were obtained by regressing out chronological age and study as covariates; positive values indicate epigenetic age acceleration relative to the value expected for the given chronological age. BH-adjusted p-values reflect pairwise comparisons between groups. Each dot represents a sample collected from individual mice, n = 6-12 per group. **(L)** Transcriptomic age residuals (tAge residual, months) of livers from young donor and recipient mice (young-to-young and old-to-young transplantation groups) compared to young sham controls, estimated using BR rodent multi-tissue transcriptomic chronological clock. Residuals were obtained by regressing out chronological age, sequencing batch, and study as covariates; positive values indicate transcriptomic age acceleration relative to the value expected for the given chronological age. BH-adjusted p-values reflect pairwise comparisons between groups, assessed with mixed-effects model. Each dot represents a sample collected from individual mice, n = 7-11 per group. **(M)** Epigenetic age residuals (eAge residual, years) of livers from old donor and recipient mice (young-to-old and old-to-old transplantation groups) compared to old sham controls, estimated using Universal pan-tissue Clock 3. Residuals were obtained by regressing out chronological age and study as covariates; positive values indicate epigenetic age acceleration relative to the value expected for the given chronological age. BH-adjusted p-values reflect pairwise comparisons between groups. Each dot represents a sample collected from individual mice, n = 6-16 per group. **(N)** Epigenetic age residuals (eAge residual, years) of livers from old donor and recipient mice (young-to-old and old-to-old transplantation groups) compared to old sham controls, estimated using Universal pan-tissue Clock 2. Residuals were obtained by regressing out chronological age and study as covariates; positive values indicate epigenetic age acceleration relative to the value expected for the given chronological age. BH-adjusted p-values reflect pairwise comparisons between groups. Each dot represents a sample collected from individual mice, n = 6-16 per group. **(O)** Transcriptomic age residuals (tAge residual, months) of livers from old donor and recipient mice (young-to-young and old-to-young transplantation groups) compared to old sham controls, estimated using BR rodent multi-tissue transcriptomic chronological clock. Residuals were obtained by regressing out chronological age, sequencing batch, and study as covariates; positive values indicate transcriptomic age acceleration relative to the value expected for the given chronological age. BH-adjusted p-values reflect pairwise comparisons between groups, assessed with mixed-effects model. Each dot represents a sample collected from individual mice, n = 4-13 per group. **(P)** Expected hazard ratio residuals (in log scale) (Mortality tAge residual, log_10_HR) of livers from old donor and recipient mice (young-to-old and old-to-old transplantation groups) compared to old sham controls, estimated using BR rodent multi-tissue mortality clock. Residuals were obtained by regressing out chronological age, sequencing batch, and study as covariates; positive values indicate transcriptomic age acceleration relative to the value expected for the given chronological age. BH-adjusted p-values reflect pairwise comparisons between groups, assessed with mixed-effects model. Each dot represents a sample collected from individual mice, n = 4-13 per group. **(Q)** Epigenetic age residuals (eAge residual, years) of blood from young donor and recipient mice (young-to-young and old-to-young transplantation groups) compared to young sham controls, estimated using Universal pan-tissue Clock 3. Residuals were obtained by regressing out chronological age and study as covariates; positive values indicate epigenetic age acceleration relative to the value expected for the given chronological age. BH-adjusted p-values reflect pairwise comparisons between groups. Each dot represents a sample collected from individual mice, n = 6-10 per group. **(R)** Epigenetic age residuals (eAge residual, years) of blood from young donor and recipient mice (young-to-young and old-to-young transplantation groups) compared to young sham controls, estimated using Universal pan-tissue Clock 2. Residuals were obtained by regressing out chronological age and study as covariates; positive values indicate epigenetic age acceleration relative to the value expected for the given chronological age. BH-adjusted p-values reflect pairwise comparisons between groups. Each dot represents a sample collected from individual mice, n = 6-10 per group. **(S)** Epigenetic age residuals (eAge residual, years) of blood from old donor and recipient mice (young-to-old and old-to-old transplantation groups) compared to old sham controls, estimated using Universal pan-tissue Clock 3. Residuals were obtained by regressing out chronological age and study as covariates; positive values indicate epigenetic age acceleration relative to the value expected for the given chronological age. BH-adjusted p-values reflect pairwise comparisons between groups. Each dot represents a sample collected from individual mice, n = 6-15 per group. **(T)** Epigenetic age residuals (eAge residual, years) of blood from old donor and recipient mice (young-to-old and old-to-old transplantation groups) compared to old sham controls, estimated using Universal pan-tissue Clock 2. Residuals were obtained by regressing out chronological age and study as covariates; positive values indicate epigenetic age acceleration relative to the value expected for the given chronological age. BH-adjusted p-values reflect pairwise comparisons between groups. Each dot represents a sample collected from individual mice, n = 6-15 per group. Boxplots: center line indicates median; box limits, interquartile range; whiskers, ±1.5× IQR. Unless specified otherwise, statistical differences between groups are assessed with ANOVA and adjusted for multiple comparisons with the Benjamini-Hochberg approach.

**Supplementary figure 7.**
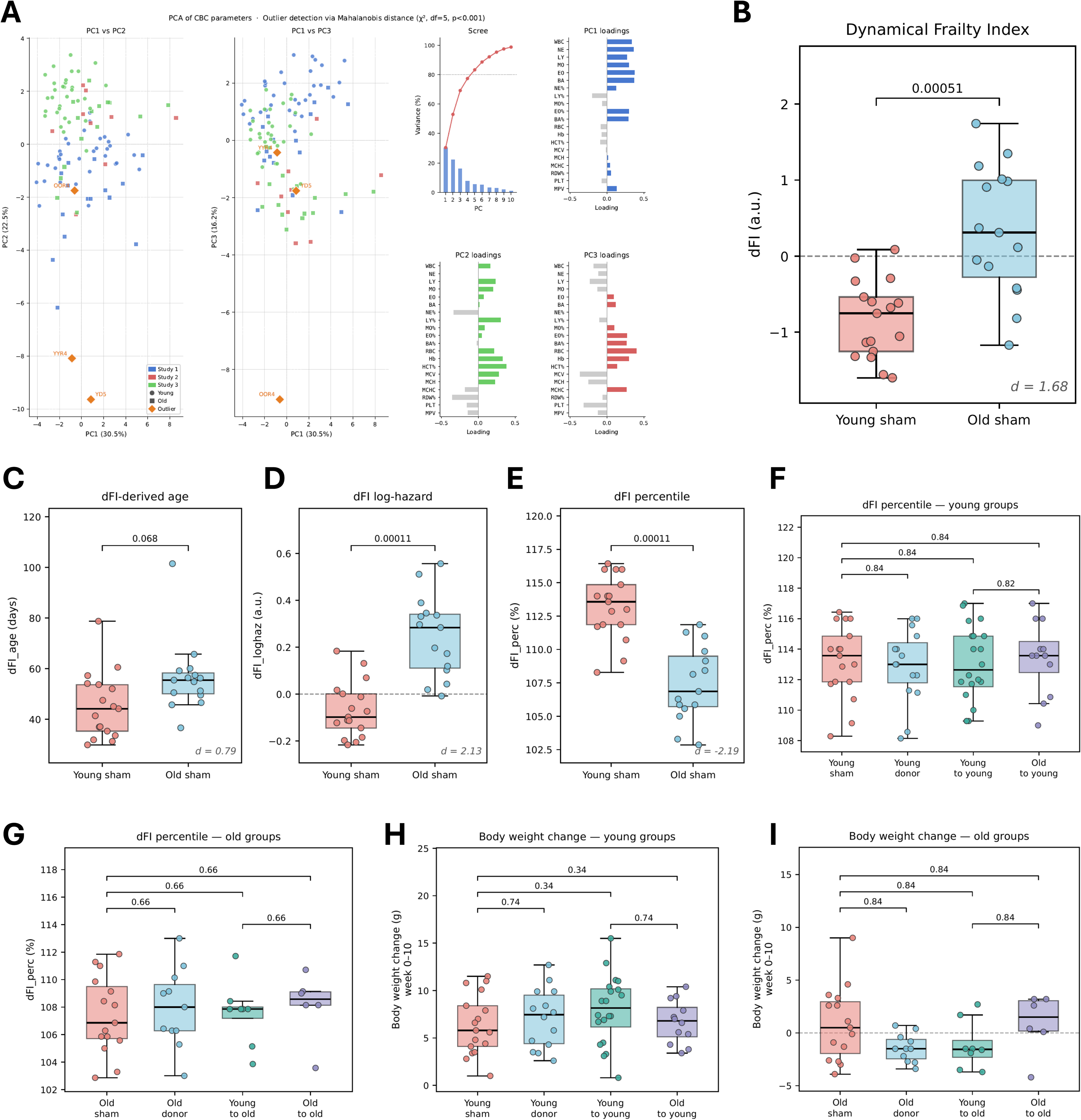
Heterochronic WAT transplantation has a limited impact on host systemic aging. **(A) (**Left) Scores plot of PC1 vs PC2. (Centre-left) Scores plot of PC1 vs PC3. Points are colored by study and shaped by age group (circles: young; squares: old). Animals flagged as multivariate outliers (Mahalanobis distance exceeding the chi-squared threshold at p < 0.001, df = 5) are shown as orange diamonds and labelled with their sample identifier. (Centre-right) Scree plot showing the percentage of variance explained by each principal component (bars) and cumulative variance (red line); the dashed line marks 80% cumulative variance. (Right panels) Loading bar charts for PC1, PC2, and PC3, showing the contribution of each CBC parameter. Positive loadings (coloured bars) and negative loadings (grey bars) are shown. **(B-E) (B)** dynamic Frailty Indicator (dFI); **(C)** dFI embeddings-based predictor of chronological age (dFI_age); **(D)** dFI embeddings-based predictor of log-hazard (dFI_loghaz); **(E)** dFI embeddings-based predictor of hazard percentile (dFI_perc). Comparisons are between young sham (n = 17) and old sham (n = 15) animals pooled across three independent studies. ANCOVA covariate: study batch. BH-adjusted p-values reflect pairwise comparisons between groups. Data points are individual mice. Cohen’s d is shown in italics (bottom right of each panel). **(F)** Hazard percentile scores (dFI_perc) of young donor and recipient mice (young-to-young and old-to-young transplantation groups) compared to young sham controls. BH-adjusted p-values reflect pairwise comparisons between groups. ANCOVA covariates: study batch and chronological age. Data points are individual mice. **(G)** Hazard percentile scores (dFI_perc) of old donor and recipient mice (young-to-old and old-to-old transplantation groups) compared to old sham controls. BH-adjusted p-values reflect pairwise comparisons between groups. ANCOVA covariates: study batch and chronological age. Data points are individual mice. **(H)** Body weight changes (week 10 minus week 0, in grams) of young donor and recipient mice (young-to-young and old-to-young transplantation groups) compared to young sham controls. BH-adjusted p-values reflect pairwise comparisons between groups. ANCOVA covariates: study batch and chronological age. Data points are individual mice. **(I)** Body weight changes (week 10 minus week 0, in grams) of old donor and recipient mice (young-to-old and old-to-old transplantation groups) compared to old sham controls. The dashed horizontal line marks zero (no change). BH-adjusted p-values reflect pairwise comparisons between groups. ANCOVA covariates: study batch and chronological age. Data points are individual mice. Boxplots: center line indicates median; box limits, interquartile range; whiskers, ±1.5× IQR. Mice per group: young sham = 17; young donor = 15 (14 for weight data); young recipient of young fat = 20; young recipient of old fat = 12; old sham = 15; old donor = 11; old recipient of young fat = 8; old recipient of old fat = 6. Unless specified otherwise, statistical differences between groups are assessed with ANOVA and adjusted for multiple comparisons with the Benjamini-Hochberg approach. *** p.adjusted< 0.001, ** p.adjusted < 0.01, * p.adjusted < 0.05, ^ p.adjusted < 0.1.

**Supplementary figure 8.**
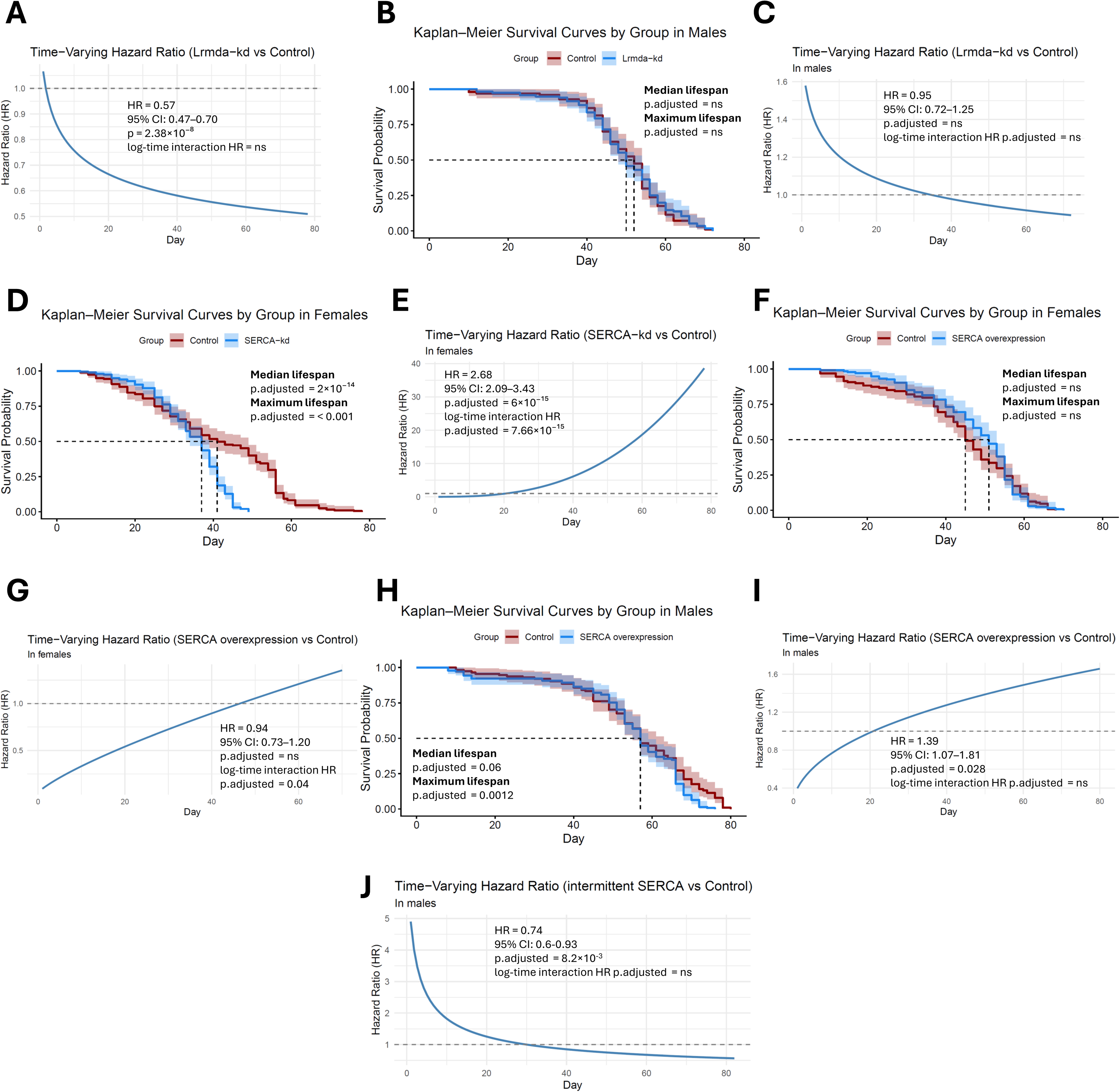
Rejuvenation-associated genes *Lrmda* and *SERCA* modulate lifespan in Drosophila. **(A)** Time-varying hazard ratio of Lrmda knockdown (Lrmda-kd) males and females relative to control flies, with sexes pooled and sex included as a covariate. Hazards were estimated using a Cox proportional hazards model and a Cox model with a time-varying coefficient. Values above 1 indicate increased mortality risk, and values below 1 indicate decreased mortality risk relative to controls at the corresponding timepoint. HR, 95% CI, and p-value reflect the overall Cox proportional hazards model estimates; the log-time interaction HR tests whether the hazard ratio changes significantly over time. **(B)** Kaplan-Meier survival curves for Lrmda knockdown (Lrmda-kd) and control males. Median lifespan was compared using the log-rank test; maximum lifespan was compared using the quantile regression (90th percentile) test. BH-adjusted p-values are shown. Shaded bands indicate 95% confidence intervals. **(C)** Time-varying hazard ratio of Lrmda knockdown (Lrmda-kd) males relative to control flies. Hazards were estimated using a Cox proportional hazards model and a Cox model with a time-varying coefficient. Values above 1 indicate increased mortality risk, and values below 1 indicate decreased mortality risk relative to controls at the corresponding timepoint. HR, 95% CI, and BH-adjusted p-value reflect the overall Cox proportional hazards model estimates; the log-time interaction HR tests whether the hazard ratio changes significantly over time. **(D)** Kaplan-Meier survival curves for SERCA knockdown (SERCA-kd) and control females. Median lifespan was compared using the log-rank test; maximum lifespan was compared using the quantile regression (90th percentile) test. BH-adjusted p-values are shown. Shaded bands indicate 95% confidence intervals. **(E)** Time-varying hazard ratio of SERCA knockdown (SERCA-kd) females relative to control flies. Hazards were estimated using a Cox proportional hazards model and a Cox model with a time-varying coefficient. Values above 1 indicate increased mortality risk, and values below 1 indicate decreased mortality risk relative to controls at the corresponding timepoint. HR, 95% CI, and BH-adjusted p-value reflect the overall Cox proportional hazards model estimates; the log-time interaction HR tests whether the hazard ratio changes significantly over time. **(F)** Kaplan-Meier survival curves for SERCA overexpression and control females. Median lifespan was compared using the log-rank test; maximum lifespan was compared using the quantile regression (90th percentile) test. BH-adjusted p-values are shown. Shaded bands indicate 95% confidence intervals. **(G)** Time-varying hazard ratio of SERCA overexpression females relative to control flies. Hazards were estimated using a Cox proportional hazards model and a Cox model with a time-varying coefficient. Values above 1 indicate increased mortality risk, and values below 1 indicate decreased mortality risk relative to controls at the corresponding timepoint. HR, 95% CI, and BH-adjusted p-value reflect the overall Cox proportional hazards model estimates; the log-time interaction HR tests whether the hazard ratio changes significantly over time. **(H)** Kaplan-Meier survival curves for SERCA overexpression and control males. Median lifespan was compared using the log-rank test; maximum lifespan was compared using the quantile regression (90th percentile) test. BH-adjusted p-values are shown. Shaded bands indicate 95% confidence intervals. **(I)** Time-varying hazard ratio of SERCA overexpression males relative to control flies. Hazards were estimated using a Cox proportional hazards model and a Cox model with a time-varying coefficient. Values above 1 indicate increased mortality risk, and values below 1 indicate decreased mortality risk relative to controls at the corresponding timepoint. HR, 95% CI, and BH-adjusted p-value reflect the overall Cox proportional hazards model estimates; the log-time interaction HR tests whether the hazard ratio changes significantly over time. **(J)** Time-varying hazard ratio of intermittent SERCA overexpression males relative to control flies. Hazards were estimated using a Cox proportional hazards model and a Cox model with a time-varying coefficient. Values above 1 indicate increased mortality risk, and values below 1 indicate decreased mortality risk relative to controls at the corresponding timepoint. HR, 95% CI, and BH-adjusted p-value reflect the overall Cox proportional hazards model estimates; the log-time interaction HR tests whether the hazard ratio changes significantly over time. Number of flies per experimental condition = 97-204; number of vials per condition = 5-9.

## References

1. Gladyshev, V. N. The Origin of Aging: Imperfectness-Driven Non-Random Damage Defines the Aging Process and Control of Lifespan. Trends Genet. 29, 506 (2013).

2. Gladyshev, V. N. Aging: progressive decline in fitness due to the rising deleteriome adjusted by genetic, environmental, and stochastic processes. Aging Cell vol. 15 594–602 Preprint at 10.1111/acel.12480 (2016).

3. Moldakozhayev, A., Tskhay, A. & Gladyshev, V. N. Applying deductive reasoning and the principles of particle physics to aging research. Aging 13, 22611–22622 (2021).

4. Moldakozhayev, A. & Gladyshev, V. N. Metabolism, homeostasis, and aging. Trends in Endocrinology & Metabolism 34, 158–169 (2023).

5. Baht, G. S. et al. Exposure to a youthful circulation rejuvenates bone repair through modulation of β-catenin. Nature Communications 2015 6:1 6, 1–10 (2015).

6. Conboy, I. M. et al. Rejuvenation of aged progenitor cells by exposure to a young systemic environment. Nature 2004 433:7027 433, 760–764 (2005).

7. Loffredo, F. S. et al. Growth differentiation factor 11 is a circulating factor that reverses age-related cardiac hypertrophy. Cell 153, 828–839 (2013).

8. Ruckh, J. M. et al. Cell Stem Cell Rejuvenation of Regeneration in the Aging Central Nervous System. 10.1016/j.stem.2011.11.019 (2012) doi:10.1016/j.stem.2011.11.019.

9. Villeda, S. A. et al. Young blood reverses age-related impairments in cognitive function and synaptic plasticity in mice. Nature Medicine 2014 20:6 20, 659–663 (2014).

10. Vi, L. et al. Macrophage cells secrete factors including LRP1 that orchestrate the rejuvenation of bone repair in mice. Nature Communications 2018 9:1 9, 1–12 (2018).

11. Buckley, M. T. et al. Cell-type-specific aging clocks to quantify aging and rejuvenation in neurogenic regions of the brain. Nature Aging 2022 3:1 3, 121–137 (2022).

12. Zhang, B. et al. Multi-omic rejuvenation and lifespan extension on exposure to youthful circulation. Nature Aging 2023 3:8 3, 948–964 (2023).

13. Tyshkovskiy, A. et al. Universal transcriptomic hallmarks of mammalian ageing and mortality. Nature 2026 654:8117 654, 173–188 (2026).

14. Wang, G., Song, A. & Wang, Q. A. Adipose tissue ageing: implications for metabolic health and lifespan. Nature Reviews Endocrinology 2025 1–15 (2025) doi:10.1038/s41574-025-01142-8.

15. Moreno-Mendez, E., Quintero-Fabian, S., Fernandez-Mejia, C. & Lazo-De-La-Vega-Monroy, M. L. Early-life programming of adipose tissue. Nutr. Res. Rev. 33, 244–259 (2020).

16. Arneson, A. et al. A mammalian methylation array for profiling methylation levels at conserved sequences. Nat. Commun. 13, 783 (2022).

17. Alvarez-Kuglen, M. et al. ImAge quantitates aging and rejuvenation. Nat. Aging 4, 1308–1327 (2024).

18. Lu, A. T. et al. Universal DNA methylation age across mammalian tissues. Nature Aging 2023 3:9 3, 1144–1166 (2023).

19. Tikhonov, S. & Dmitriev, S. E. Heterogeneous epigenetic regulatory patterns link mammalian aging, development, and mortality. bioRxiv 2026.09.08.750031 (2026) doi:10.64898/2026.09.08.750031.

20. Mozhui, K. et al. Genetic loci and metabolic states associated with murine epigenetic aging. Elife 11, (2022).

21. Tomusiak, A. et al. Development of an epigenetic clock resistant to changes in immune cell composition. Commun. Biol. 7, 934 (2024).

22. Moqri, M., Poganik, J. R., Horvath, S. & Gladyshev, V. N. What makes biological age epigenetic clocks tick. Nat. Aging 5, 335 (2025).

23. Shchukina, I. et al. Inflammaging in aged tissues drives remodeling of the CD8+T cell compartment. Cell Rep. 45, (2026).

24. Tyshkovskiy, A. et al. Distinct longevity mechanisms across and within species and their association with aging. Cell 186, 2929–2949.e20 (2023).

25. Kurokawa, J. et al. Macrophage-derived AIM Is endocytosed into adipocytes and decreases lipid droplets via inhibition of fatty acid synthase activity. Cell Metab. 11, 479–492 (2010).

26. Iwamura, Y. et al. Apoptosis inhibitor of macrophage (AIM) diminishes lipid droplet-coating proteins leading to lipolysis in adipocytes. Biochem. Biophys. Res. Commun. 422, 476–481 (2012).

27. Kurokaw, J. et al. Apoptosis inhibitor of macrophage (AIM) is required for obesity-associated recruitment of inflammatory macrophages into adipose tissue. Proc. Natl. Acad. Sci. U. S. A. 108, 12072–12077 (2011).

28. Uhlén, M. et al. Tissue-based map of the human proteome. Science (1979). 347, (2015).

29. Jakobsson, P. J., Mancini, J. A., Riendeau, D. & Ford-Hutchinson, A. W. Identification and characterization of a novel microsomal enzyme with glutathione-dependent transferase and peroxidase activities. Journal of Biological Chemistry 272, 22934–22939 (1997).

30. MGST3 microsomal glutathione S-transferase 3 [Homo sapiens (human)] - Gene - NCBI. https://www.ncbi.nlm.nih.gov/gene?Db=gene&Cmd=ShowDetailView&TermToSearch=4259.

31. Alnasser, S. M. The role of glutathione S-transferases in human disease pathogenesis and their current inhibitors. Genes Dis. 12, 101482 (2024).

32. De Victoria, E. O. M. et al. Macrophage Content in Subcutaneous Adipose Tissue: Associations With Adiposity, Age, Inflammatory Markers, and Whole-Body Insulin Action in Healthy Pima Indians. Diabetes 58, 385 (2009).

33. Moreno-Navarrete, J. M. et al. The MRC1/CD68 Ratio Is Positively Associated with Adipose Tissue Lipogenesis and with Muscle Mitochondrial Gene Expression in Humans. PLoS One 8, e70810 (2013).

34. Kiran, S., Kumar, V., Murphy, E. A., Enos, R. T. & Singh, U. P. High Fat Diet-Induced CD8+ T Cells in Adipose Tissue Mediate Macrophages to Sustain Low-Grade Chronic Inflammation. Front. Immunol. 12, 680944 (2021).

35. Rosen, E. D. et al. C/EBPα induces adipogenesis through PPARγ: a unified pathway. Genes Dev. 16, 22 (2002).

36. Vegiopoulos, A. et al. Cyclooxygenase-2 controls energy homeostasis in mice by de novo recruitment of brown adipocytes. Science (1979). 328, 1158–1161 (2010).

37. Wu, J. et al. Beige adipocytes are a distinct type of thermogenic fat cell in mouse and human. Cell 150, 366–376 (2012).

38. Pilkington, A. C., Paz, H. A. & Wankhade, U. D. Beige Adipose Tissue Identification and Marker Specificity—Overview. Front. Endocrinol. (Lausanne). 12, 599134 (2021).

39. Rajakumari, S. et al. EBF2 determines and maintains brown adipocyte identity. Cell Metab. 17, 562 (2013).

40. Ikeda, K. et al. UCP1-independent signaling involving SERCA2b-mediated calcium cycling regulates beige fat thermogenesis and systemic glucose homeostasis. Nature Medicine 2017 23:12 23, 1454–1465 (2017).

41. Li, J. et al. TMTpro-18plex: The Expanded and Complete Set of TMTpro Reagents for Sample Multiplexing. J. Proteome Res. 20, 2964–2972 (2021).

42. Alam, M. S. et al. Genomic Expression Analyses Reveal Lysosomal, Innate Immunity Proteins, as Disease Correlates in Murine Models of a Lysosomal Storage Disorder. PLoS One 7, e48273 (2012).

43. Houseknecht, K. L. & Spurlock, M. E. Leptin regulation of lipid homeostasis: dietary and metabolic implications. Nutr. Res. Rev. 16, 83–96 (2003).

44. Gupta, A. et al. Human CIDEC transgene improves lipid metabolism and protects against high-fat diet–induced glucose intolerance in mice. J. Biol. Chem. 298, 102347 (2022).

45. Singer, J. M. et al. Monoacylglycerol O-acyltransferase 1 lowers adipocyte differentiation capacity in vitro but does not affect adiposity in mice. Obesity (Silver Spring) 30, 2122 (2022).

46. Von Bank, H., Kirsh, C. & Simcox, J. Aging adipose: Depot location dictates age-associated expansion and dysfunction. Ageing Res. Rev. 67, 101259 (2021).

47. Divoux, A. et al. Mast Cells in Human Adipose Tissue: Link with Morbid Obesity, Inflammatory Status, and Diabetes. J. Clin. Endocrinol. Metab. 97, E1677–E1685 (2012).

48. Sun, K., Tordjman, J., Clément, K. & Scherer, P. E. Fibrosis and Adipose Tissue Dysfunction. Cell Metab. 18, 470–477 (2013).

49. Liu, X. et al. Obesity induces adipose fibrosis and collagen cross-linking through suppressing AMPK and enhancing lysyl oxidase expression. Biochim. Biophys. Acta Mol. Basis Dis. 1868, 166454 (2022).

50. Halberg, N. et al. Hypoxia-Inducible Factor 1α Induces Fibrosis and Insulin Resistance in White Adipose Tissue. Mol. Cell. Biol. 29, 4467–4483 (2009).

51. Pahal, S., Mainali, N., Balasubramaniam, M., Shmookler Reis, R. J. & Ayyadevara, S. Mitochondria in aging and age-associated diseases. Mitochondrion 82, 102022 (2025).

52. Traa, A. et al. Overexpression of mitochondrial fission or mitochondrial fusion genes enhances resilience and extends longevity. Aging Cell 23, e14262 (2024).

53. Avchaciov, K. et al. Unsupervised learning of aging principles from longitudinal data. Nat. Commun. 13, (2022).

54. Scialo, F., Sriram, A., Stefanatos, R. & Sanz, A. Practical Recommendations for the Use of the GeneSwitch Gal4 System to Knock-Down Genes in Drosophila melanogaster. PLoS One 11, e0161817 (2016).

55. Osterwalder, T., Yoon, K. S., White, B. H. & Keshishian, H. A conditional tissue-specific transgene expression system using inducible GAL4. Proc. Natl. Acad. Sci. U. S. A. 98, 12596–12601 (2001).

56. Konige, M., Wang, H. & Sztalryd, C. Role of adipose specific lipid droplet proteins in maintaining whole body energy homeostasis. Biochimica et Biophysica Acta (BBA) - Molecular Basis of Disease 1842, 393–401 (2014).

57. Olzmann, J. A. & Carvalho, P. Dynamics and functions of lipid droplets. Nat. Rev. Mol. Cell Biol. 20, 137 (2019).

58. Walther, T. C. & Farese, R. V. Lipid Droplets And Cellular Lipid Metabolism. Annu. Rev. Biochem. 81, 687 (2012).

59. Zhang, W. et al. Lipid Droplets, the Central Hub Integrating Cell Metabolism and the Immune System. Front. Physiol. 12, 746749 (2021).

60. Reyes-Farias, M., Fos-Domenech, J., Serra, D., Herrero, L. & Sánchez-Infantes, D. White adipose tissue dysfunction in obesity and aging. Biochem. Pharmacol. 192, 114723 (2021).

61. Ou, M. Y., Zhang, H., Tan, P. C., Zhou, S. B. & Li, Q. F. Adipose tissue aging: mechanisms and therapeutic implications. Cell Death & Disease 2022 13:4 13, 1–10 (2022).

62. Mancuso, P. & Bouchard, B. The Impact of Aging on Adipose Function and Adipokine Synthesis. Front. Endocrinol. (Lausanne). 10, 137 (2019).

63. Nguyen, T. T. & Corvera, S. Adipose tissue as a linchpin of organismal ageing. Nature Metabolism 2024 6:5 6, 793–807 (2024).

64. Nishimoto, Y. & Tamori, Y. CIDE Family-Mediated Unique Lipid Droplet Morphology in White Adipose Tissue and Brown Adipose Tissue Determines the Adipocyte Energy Metabolism. J. Atheroscler. Thromb. 24, 989 (2017).

65. D’Silva, L., Cheng, C. & Barrow, J. J. Unlocking thermogenic silencers for the treatment of metabolic disease. Trends in Endocrinology & Metabolism 0, (2025).

66. Perez-Campo, R., López-Torres, M., Cadenas, S., Rojas, C. & Barja, G. The rate of free radical production as a determinant of the rate of aging: Evidence from the comparative approach. J. Comp. Physiol. B 168, 149–158 (1998).

67. Poganik, J. R. et al. Aging vs. biological age: Conceptual considerations for age reversal claims. Cell Metab. 38, 1746–1750 (2026).

68. Olaisen, B. F. et al. Replacement-Based Ageing Interventions for Systemic Rejuvenation: Shaping Longevity Science and Clinical Directions. Aging Cell 25, e70516 (2026).

69. Kong, X. et al. Brown Adipose Tissue Controls Skeletal Muscle Function via the Secretion of Myostatin. Cell Metab. 28, 631–643.e3 (2018).

70. Chen, Y. et al. Thermal stress induces glycolytic beige fat formation via a myogenic state. Nature 2018 565:7738 565, 180–185 (2018).

71. Miyano, C. A., Orezzoli, S. F., Loren Buck, C. & Nishikawa, K. C. Severe thermoregulatory deficiencies in mice with a deletion in the titin gene TTN. Journal of Experimental Biology 222, (2019).

72. Behrens, J. et al. Single-nucleus mRNA-sequencing reveals dynamics of lipogenic and thermogenic adipocyte populations in murine brown adipose tissue in response to cold exposure. Mol. Metab. 101, 102252 (2025).

73. Di Paola, J. N. & Schöck, F. The varied functions of the giant muscle scaffold protein obscurin. Front. Cell Dev. Biol. 13, 1689326 (2026).

74. Adekeye, T. E. et al. Myosin light chain proteins cooperatively promote sarcomere growth in fast-twitch muscle. Nature Communications 2026 10.1038/s41467-026-73861-z (2026) doi:10.1038/s41467-026-73861-z.

75. Waldén, T. B., Hansen, I. R., Timmons, J. A., Cannon, B. & Nedergaard, J. Recruited vs. nonrecruited molecular signatures of brown, ‘brite,’ and white adipose tissues. Am. J. Physiol. Endocrinol. Metab. 302, 19–31 (2012).

76. Zhang, Z. et al. Transcriptomics profiling reveal the heterogeneity of white and brown adipocyte. J. Bioenerg. Biomembr. 55, 423–433 (2023).

77. Diao, Z. et al. Possibility of uncoupling protein 1 expression in bovine fast-twitch muscle fibers. Journal of Veterinary Medical Science 85, 587–591 (2023).

78. Tharp, K. M. et al. Actomyosin-Mediated Tension Orchestrates Uncoupled Respiration in Adipose Tissues. Cell Metab. 27, 602–615.e4 (2018).

79. Park, G. et al. The transcriptional modulator Ifrd1 controls PGC-1α expression under short-term adrenergic stimulation in brown adipocytes. FEBS J. 284, 784–795 (2017).

80. Strehlau, C. et al. Adipocyte Myoglobin Is a Determinant of Energy Expenditure and a Potential Target to Limit Obesity. Advanced Science e76191 (2026) doi:10.1002/ADVS.76191.

81. Christen, L. et al. Myoglobin-mediated lipid shuttling increases adrenergic activation of brown and white adipocyte metabolism and is as a marker of thermogenic adipocytes in humans. Clin. Transl. Med. 12, e1108 (2022).

82. Aboouf, M. A. et al. Myoglobin, expressed in brown adipose tissue of mice, regulates the content and activity of mitochondria and lipid droplets. Biochimica et Biophysica Acta (BBA) - Molecular and Cell Biology of Lipids 1866, 159026 (2021).

83. Aboouf, M. A., Gorr, T. A., Hamdy, N. M., Gassmann, M. & Thiersch, M. Myoglobin in Brown Adipose Tissue: A Multifaceted Player in Thermogenesis. Cells 12, 2240 (2023).

84. Sun, Y. et al. Genome-wide survey identifies TNNI2 as a target of KLF7 that inhibits chicken adipogenesis via downregulating FABP4. Biochimica et Biophysica Acta (BBA) - Gene Regulatory Mechanisms 1866, 194899 (2023).

85. Ma, X. et al. ORMDL3 and its implication in inflammatory disorders. Int. J. Rheum. Dis. 21, 1154–1162 (2018).

86. Song, Y. et al. Ablation of ORMDL3 impairs adipose tissue thermogenesis and insulin sensitivity by increasing ceramide generation. Mol. Metab. 56, (2022).

87. Mooli, R. G. R. et al. Epigenetically active chromatin in neonatal iWAT reveals GABPα as a potential regulator of beige adipogenesis. Front. Endocrinol. (Lausanne). 15, 1385811 (2024).

88. Murari, A. et al. Dissecting the concordant and disparate roles of NDUFAF3 and NDUFAF4 in mitochondrial complex I biogenesis. iScience 24, 102869 (2021).

89. Zou, M. et al. Txnip deficiency causes a susceptibility to acute cold stress with brown fat dysfunction in mice. J. Biol. Chem. 301, 108293 (2025).

90. Chutkow, W. A. & Lee, R. T. Thioredoxin Regulates Adipogenesis through Thioredoxin-interacting Protein (Txnip) Protein Stability. Journal of Biological Chemistry 286, 29139–29145 (2011).

91. Rahbani, J. F. et al. ADRA1A–Gαq signalling potentiates adipocyte thermogenesis through CKB and TNAP. Nature Metabolism 2022 4:11 4, 1459–1473 (2022).

92. Kazak, L. & Spiegelman, B. M. Mechanism of futile creatine cycling in thermogenesis. 10.1152/ajpendo.00444.2020 319, E947–E949 (2020).

93. Bal, N. C. et al. Sarcolipin is a newly identified regulator of muscle-based thermogenesis in mammals. Nature Medicine 2012 18:10 18, 1575–1579 (2012).

94. Auger, C. et al. Identification of a molecular resistor that controls UCP1-independent Ca2+ cycling thermogenesis in adipose tissue. Cell Metab. 37, 1311–1325.e9 (2025).

95. Aquilano, K. et al. Low-protein/high-carbohydrate diet induces AMPK-dependent canonical and non-canonical thermogenesis in subcutaneous adipose tissue. Redox Biol. 36, 101633 (2020).

96. Yu, X. et al. The GIP receptor activates futile calcium cycling in white adipose tissue to increase energy expenditure and drive weight loss in mice. Cell Metab. 37, 187–204.e7 (2025).

97. Georgiou, D. K. et al. Ca2+ binding/permeation via calcium channel, CaV1.1, regulates the intracellular distribution of the fatty acid transport protein, CD36, and fatty acid metabolism. Journal of Biological Chemistry 290, 23751–23765 (2015).

98. Weninger, G. et al. Structural insights into the regulation of RyR1 by S100A1. Proc. Natl. Acad. Sci. U. S. A. 121, e2400497121 (2024).

99. Ni, X. et al. Gegen Huazhuo Tang (GHT) inhibits 7-KDCA and increases TGR5 to promote white fat browning. J. Ethnopharmacol. 352, 120266 (2025).

100. Hossain, M. A., Poojari, A. & Rabiee, A. Thermogenesis in Adipose Tissue: Adrenergic and Non-Adrenergic Pathways. Cells 2026, Vol. 15, Page 131 15, 131 (2026).

101. Kazak, L. et al. A Creatine-Driven Substrate Cycle Enhances Energy Expenditure and Thermogenesis in Beige Fat. Cell 163, 643–655 (2015).

102. Wu, Q. et al. Intestinal hypoxia-inducible factor 2α regulates lactate levels to shape the gut microbiome and alter thermogenesis. Cell Metab. 33, 1988–2003.e7 (2021).

103. Groza, T. et al. The International Mouse Phenotyping Consortium: comprehensive knockout phenotyping underpinning the study of human disease. Nucleic Acids Res. 51, D1038–D1045 (2023).

104. Wyss, M. & Kaddurah-Daouk, R. Creatine and creatinine metabolism. Physiol. Rev. 80, 1107–1213 (2000).

105. Willis, J., Jones, R., Nwokolo, N. & Levy, J. Protein and creatine supplements and misdiagnosis of kidney disease. BMJ 340, 210 (2010).

106. Chen, Y. et al. Creatine ameliorates high-fat diet-induced obesity by regulation of lipolysis and lipophagy in brown adipose tissue and liver. Biochimie 209, 85–94 (2023).

107. Furuhashi, M., Saitoh, S., Shimamoto, K. & Miura, T. Fatty Acid-Binding Protein 4 (FABP4): Pathophysiological Insights and Potent Clinical Biomarker of Metabolic and Cardiovascular Diseases. Clin. Med. Insights Cardiol. 8, 23 (2015).

108. Bahrami-Nejad, Z. et al. Early enforcement of cell identity by a functional component of the terminally differentiated state. PLoS Biol. 20, e3001900 (2022).

109. Chawla, A. & Lazar, M. A. Induction of Rev-ErbAα, an orphan receptor encoded on the opposite strand of the α-thyroid hormone receptor gene, during adipocyte differentiation. Journal of Biological Chemistry 268, 16265–16269 (1993).

110. Wang, J. & Lazar, M. A. Bifunctional Role of Rev-erbα in Adipocyte Differentiation. Mol. Cell. Biol. 28, 2213 (2008).

111. Kumar, N. et al. Regulation of Adipogenesis by Natural and Synthetic REV-ERB Ligands. Endocrinology 151, 3015–3025 (2010).

112. Gerhart-Hines, Z. et al. The nuclear receptor Rev-erbα controls circadian thermogenic plasticity. Nature 2013 503:7476 503, 410–413 (2013).

113. Kuuskmäe, C. et al. Negr1 deficiency alters glutamate signalling and kynurenine pathway in a mouse model of psychiatric disorders. Scientific Reports 2026 16:1 16, 5317- (2026).

114. Nigro, P. et al. Exercise training remodels inguinal white adipose tissue through adaptations in innervation, vascularization, and the extracellular matrix. CellReports 42, 112392 (2023).

115. Kim, S. Y., Park, S. Y. & Kim, J. E. GULP1 deficiency reduces adipogenesis and glucose uptake via downregulation of PPAR signaling and disturbing of insulin/ERK signaling in 3T3-L1 cells. J. Cell. Physiol. 239, e31173 (2024).

116. Li, G. H., Arora, P. D., Chen, Y., Mcculloch, C. A. & Liu, P. Multifunctional roles of gelsolin in health and diseases. Med. Res. Rev. 32, 999–1025 (2012).

117. Mukherjee, R. & Yun, J. W. Long chain acyl CoA synthetase 1 and gelsolin are oppositely regulated in adipogenesis and lipogenesis. Biochem. Biophys. Res. Commun. 420, 588–593 (2012).

118. Kawaji, A., Ohnaka, Y., Osada, S., Nishizuka, M. & Imagawa, M. Gelsolin, an Actin Regulatory Protein, is Required for Differentiation of Mouse 3T3-L1 Cells into Adipocytes. Biol. Pharm. Bull. 33, 773–779 (2010).

119. Costa, I. P. D. et al. AQP7 deficiency drives adipose tissue remodeling and disrupts homeostasis. npj Metabolic Health and Disease 2025 3:1 3, 44- (2025).

120. Ji, Y. K., Tillison, K., Lee, J. H., Rearick, D. A. & Smas, C. M. The adipose tissue triglyceride lipase ATGL/PNPLA2 is downregulated by insulin and TNF-α in 3T3-L1 adipocytes and is a target for transactivation by PPARγ. Am. J. Physiol. Endocrinol. Metab. 291, 115–127 (2006).

121. Roy, D. et al. Coordinated transcriptional control of adipocyte triglyceride lipase (Atgl) by transcription factors Sp1 and peroxisome proliferator-activated receptor γ (PPARγ) during adipocyte differentiation. Journal of Biological Chemistry 292, 14827–14835 (2017).

122. Sakaue, H. et al. Requirement of fibroblast growth factor 10 in development of white adipose tissue. Genes Dev. 16, 908–912 (2002).

123. Asaki, T. et al. Roles of fibroblast growth factor 10 (Fgf10) in adipogenesis in vivo. Mol. Cell. Endocrinol. 218, 119–128 (2004).

124. Fatima, L. A. et al. Estradiol stimulates adipogenesis and Slc2a4/GLUT4 expression via ESR1-mediated activation of CEBPA. Mol. Cell. Endocrinol. 498, 110447 (2019).

125. Kadota, Y. et al. Metallothioneins regulate the adipogenic differentiation of 3T3-L1 cells via the insulin signaling pathway. PLoS One 12, e0176070 (2017).

126. Kawakami, T. et al. Regulatory role of metallothionein-1/2 on development of sex differences in a high-fat diet-induced obesity. Life Sci. 226, 12–21 (2019).

127. Yang, X. et al. Analysis of the different characteristics between omental preadipocytes and differentiated white adipocytes using bioinformatics methods. Adipocyte 11, 227 (2022).

128. Yamashiro, S. et al. Citron Kinase, a Rho-dependent Kinase, Induces Di-phosphorylation of Regulatory Light Chain of Myosin II. Mol. Biol. Cell 14, 1745–1756 (2003).

129. Gai, M. et al. Citron kinase controls abscission through RhoA and anillin. Mol. Biol. Cell 22, 3768–3778 (2011).

130. Shou, J., Yu, C., Zhang, D. & Zhang, Q. Overexpression of Citron Rho-Interacting Serine/Threonine Kinase Associated with Poor Outcome in Bladder Cancer. J. Cancer 11, 4173 (2020).

131. Ji, Y., et al. Rock signaling control PPARγ expression and actin polymerization during adipogenesis. Saudi J. Biol. Sci. 24, 1866 (2017).

132. Diep, D. T. V. et al. Anti-adipogenic effects of KD025 (SLx-2119), a ROCK2-specific inhibitor, in 3T3-L1 cells. Scientific Reports 2018 8:1 8, 2477- (2018).

133. Dimitrova, N., Chen, Y. C. M., Spector, D. L. & De Lange, T. 53BP1 promotes non-homologous end joining of telomeres by increasing chromatin mobility. Nature 2008 456:7221 456, 524–528 (2008).

134. Zimmermann, M. & De Lange, T. 53BP1: Pro choice in DNA repair. Trends Cell Biol. 24, 108–117 (2014).

135. Molchadsky, A. et al. p53 is required for brown adipogenic differentiation and has a protective role against diet-induced obesity. Cell Death Differ. 20, 774 (2013).

136. Park, C. H., Rhyu, D. Y., Sharma, B. R. & Yokozawa, T. Inhibition of preadipocyte differentiation and lipid accumulation by 7-O-galloyl-d-sedoheptulose treatment in 3T3-L1 adipocytes. Biomedicine & Preventive Nutrition 3, 319–324 (2013).

137. Li, X. Glutathione and Glutathione-S-Transferase in Detoxification Mechanisms. General, Applied and Systems Toxicology: Ballantyne/Toxicology 1–13 (2009) doi:10.1002/9780470744307.GAT166.

138. De Paepe, B., Merckx, C., Jarošová, J., Cannizzaro, M. & De Bleecker, J. L. Myo-Inositol Transporter SLC5A3 Associates with Degenerative Changes and Inflammation in Sporadic Inclusion Body Myositis. Biomolecules 2020, Vol. 10, Page 521 10, 521 (2020).

139. Banerjee, D. & Girirajan, S. Discovery of obesity genes through cross-ancestry analysis. Nature Communications 2025 16:1 16, 9319- (2025).

140. Taylor, J. A. et al. Prenatal Exposure to Bisphenol A Disrupts Naturally Occurring Bimodal DNA Methylation at Proximal Promoter of fggy, an Obesity-Relevant Gene Encoding a Carbohydrate Kinase, in Gonadal White Adipose Tissues of CD-1 Mice. Endocrinology 159, 779–794 (2018).

141. Attia, M. et al. Interaction between Nucleosome Assembly Protein 1-like Family Members. J. Mol. Biol. 407, 647–660 (2011).

142. Lee, M. tse G., et al. Integrative Genome-Wide Association Studies of eQTL and GWAS Data for Gout Disease Susceptibility. Scientific Reports 2019 9:1 9, 4981- (2019).

143. Brotman, S. M. et al. Cell-Type Composition Affects Adipose Gene Expression Associations With Cardiometabolic Traits. Diabetes 72, 1707 (2023).

144. TUBE1 Gene - GeneCards. https://www.genecards.org/card/TUBE1.

145. Zhu, Y. et al. Adipose tissue hyaluronan production improves systemic glucose homeostasis and primes adipocytes for CL 316,243-stimulated lipolysis. Nature Communications 2021 12:1 12, 4829- (2021).

146. Robert, F. et al. The Transcriptional Histone Acetyltransferase Cofactor TRRAP Associates with the MRN Repair Complex and Plays a Role in DNA Double-Strand Break Repair. Mol. Cell. Biol. 26, 402–412 (2006).

147. Körner, M. et al. The FAM104 proteins VCF1/2 promote the nuclear localization of p97/VCP. Elife 12, (2023).

148. Subramanian, A. et al. Gene set enrichment analysis: A knowledge-based approach for interpreting genome-wide expression profiles. Proc. Natl. Acad. Sci. U. S. A. 102, 15545–15550 (2005).

149. Liberzon, A. et al. The Molecular Signatures Database (MSigDB) hallmark gene set collection. Cell Syst. 1, 417 (2015).

150. Liberzon, A. et al. Molecular signatures database (MSigDB) 3.0. Bioinformatics 27, 1739–1740 (2011).

151. Castanza, A. S. et al. Extending support for mouse data in the Molecular Signatures Database (MSigDB). Nature Methods 2023 20:11 20, 1619–1620 (2023).

152. Kanehisa, M. & Goto, S. KEGG: Kyoto Encyclopedia of Genes and Genomes. Nucleic Acids Res. 28, 27–30 (2000).

153. Tyshkovskiy, A. et al. Identification and Application of Gene Expression Signatures Associated with Lifespan Extension. Cell Metab. 30, 573–593.e8 (2019).

154. Emont, M. P. et al. A single-cell atlas of human and mouse white adipose tissue. Nature 2022 603:7903 603, 926–933 (2022).

155. Wu, T. et al. clusterProfiler 4.0: A universal enrichment tool for interpreting omics data. Innovation (Cambridge (Mass.)) 2, (2021).

156. Ninomiya, K., Ortaleza, K., Lopez, J. A. & Terskikh, A. V. Measuring Single-Cell Aging with an Imaging-based Biomarker of Chromatin and Epigenetic Aging. J. Vis. Exp. 2026-January, (2026).

157. Eng, J. K., Jahan, T. A. & Hoopmann, M. R. Comet: an open-source MS/MS sequence database search tool. Proteomics 13, 22–24 (2013).

158. Sziraki, A. et al. A global view of aging and Alzheimer’s pathogenesis-associated cell population dynamics and molecular signatures in human and mouse brains. Nature Genetics 2023 55:12 55, 2104–2116 (2023).

159. Robinson, M. D., McCarthy, D. J. & Smyth, G. K. edgeR: a Bioconductor package for differential expression analysis of digital gene expression data. Bioinformatics 26, 139 (2009).

160. Ritchie, M. E. et al. Limma powers differential expression analyses for RNA-sequencing and microarray studies. Nucleic Acids Res. 43, e47 (2015).

161. Chambers, M. C. et al. A cross-platform toolkit for mass spectrometry and proteomics. Nature Biotechnology 2012 30:10 30, 918–920 (2012).

162. Koenker, R. & Bassett, G. Regression Quantiles. Econometrica 46, 33 (1978).

163. Yu, G. & He, Q. Y. ReactomePA: an R/Bioconductor package for reactome pathway analysis and visualization. Mol. Biosyst. 12, 477–479 (2016).

164. Kirillov, A. et al. Segment Anything. Proceedings of the IEEE International Conference on Computer Vision 3992–4003 (2023) doi:10.1109/ICCV51070.2023.00371.

165. Hao, Y. et al. Dictionary learning for integrative, multimodal and scalable single-cell analysis. Nature Biotechnology 2023 42:2 42, 293–304 (2023).

166. Aran, D. et al. Reference-based analysis of lung single-cell sequencing reveals a transitional profibrotic macrophage. Nature Immunology 2019 20:2 20, 163–172 (2019).

167. Dobin, A. et al. STAR: ultrafast universal RNA-seq aligner. Bioinformatics 29, 15 (2012).

168. Kaplan, E. L. & Meier, P. Nonparametric Estimation from Incomplete Observations. J. Am. Stat. Assoc. 53, 457–481 (1958).

169. Cox, D. R. Regression Models and Life-Tables. J. R. Stat. Soc. Series B Stat. Methodol. 34, 187–202 (1972).

170. Benjamini, Y. & Hochberg, Y. Controlling the False Discovery Rate: A Practical and Powerful Approach to Multiple Testing. Journal of the Royal Statistical Society Series B 57, 289–300 (1995).

171. Zhang, Z. et al. A panoramic view of cell population dynamics in mammalian aging. Science (1979). 387, (2025).

172. Farmer, S. R. Transcriptional control of adipocyte formation. Cell Metab. 4, 263–273 (2006).

173. Rosen, E. D. & MacDougald, O. A. Adipocyte differentiation from the inside out. Nature Reviews Molecular Cell Biology 2006 7:12 7, 885–896 (2006).

174. Scott, E. W., Simon, M. C., Anastasi, J. & Singh, H. Requirement of transcription factor PU.1 in the development of multiple hematopoietic lineages. Science 265, 1573–1577 (1994).

175. Navarrete-Perea, J., Yu, Q., Gygi, S. P. & Paulo, J. A. Streamlined Tandem Mass Tag (SL-TMT) Protocol: An Efficient Strategy for Quantitative (Phospho)proteome Profiling Using Tandem Mass Tag-Synchronous Precursor Selection-MS3. J. Proteome Res. 17, 2226–2236 (2018).

176. Mitchell, W. et al. Multi-omics characterization of partial chemical reprogramming reveals evidence of cell rejuvenation. Elife 12, (2024).

177. Beausoleil, S. A., Villén, J., Gerber, S. A., Rush, J. & Gygi, S. P. A probability-based approach for high-throughput protein phosphorylation analysis and site localization. Nat. Biotechnol. 24, 1285–1292 (2006).

178. Huttlin, E. L. et al. A tissue-specific atlas of mouse protein phosphorylation and expression. Cell 143, 1174–1189 (2010).

179. Elias, J. E. & Gygi, S. P. Target-decoy search strategy for increased confidence in large-scale protein identifications by mass spectrometry. Nat. Methods 4, 207–214 (2007).

180. Elias, J. E. & Gygi, S. P. Target-decoy search strategy for mass spectrometry-based proteomics. Methods Mol. Biol. 604, 55–71 (2010).

181. McAlister, G. C. et al. Increasing the multiplexing capacity of TMTs using reporter ion isotopologues with isobaric masses. Anal. Chem. 84, 7469–7478 (2012).

182. Dieterle, F., Ross, A., Schlotterbeck, G. & Senn, H. Probabilistic Quotient Normalization as Robust Method to Account for Dilution of Complex Biological Mixtures. Application in1H NMR Metabonomics. Anal. Chem. 78, 4281–4290 (2006).

183. van den Berg, R. A., Hoefsloot, H. C. J., Westerhuis, J. A., Smilde, A. K. & van der Werf, M. J. Centering, scaling, and transformations: Improving the biological information content of metabolomics data. BMC Genomics 7, 142- (2006).

184. Zaqout, S., Becker, L. L. & Kaindl, A. M. Immunofluorescence Staining of Paraffin Sections Step by Step. Front. Neuroanat. 14, 582218 (2020).

185. Osterwalder, T., Yoon, K. S., White, B. H. & Keshishian, H. A conditional tissue-specific transgene expression system using inducible GAL4. Proc. Natl. Acad. Sci. U. S. A. 98, 12596–12601 (2001).

